# Higher-order topology uncovers the multidimensional dynamics of naturalistic emotion

**DOI:** 10.64898/2026.09.26.754452

**Authors:** Lorenzo Zaffina, Matteo Diano, Flavia Petruso, Maria Giulia Preti, Enrico Amico, Elenor Morgenroth, Giovanni Petri, Dimitri Van De Ville, Patrik Vuilleumier, Andrea Santoro

**Affiliations:** DIAG, Sapienza University of Rome, Rome, Italy; Intesa Sanpaolo A.I. Research, Torino, Italy; Neuro-X Institute, École polytechnique fédérale de Lausanne (EPFL), Geneva, Switzerland; Department of Psychology and Neuroscience Institute of Turin (NIT), University of Torino, Turin, Italy; Department of Radiology and Medical Informatics, University of Geneva, Geneva, 1202, Switzerland; University of Birmingham, School of Mathematics & Centre for Human Brain Health, Birmingham, United Kingdom; Swiss Center for Affective Sciences, University of Geneva, Geneva, 1202, Switzerland; NP Lab, Network Science Institute, Northeastern University London, London, E1W 1LP, United Kingdom; Department of Physics, Northeastern University, Boston, USA; Department of Basic Neurosciences, University of Geneva, Geneva, 1202, Switzerland; ISI Foundation, Turin, Italy

**Keywords:** brain networks, higher-order analyses, fMRI data, naturalistic stimuli, emotions

## Abstract

Emotions are thought to emerge from co-activation among distributed brain systems, yet traditional fMRI analyses typically examine localized responses and pairwise connections, potentially overlooking interactions among groups of regions. Here, we use time-resolved higher-order topology to characterize these group interactions during naturalistic viewing of 14 films totaling over 2.5 h, continuously annotated across 50 affective features. The homological scaffold, representing evolving group-level topology, most closely tracks recurrent emotional states and best predicts fine-grained affective profiles. This sensitivity diminishes when emotion is compressed into dimensions of valence, arousal, and power, where pairwise connectivity captures the dominant arousal signal. Across three independent datasets, arousal predictions transfer most robustly through pairwise connectivity, while the relative geometry of broad affective states remains conserved across film narratives despite poor valence generalization. These findings reveal complementary neural representations of emotion: higher-order topology captures fine-grained affective structure, whereas pairwise connectivity provides a portable readout of broad arousal.

## INTRODUCTION

An emotional episode reorganizes perception, memory, choice, and social behavior within seconds. Yet, uncovering how the brain orchestrates the vivid distinctions between fear, anger, awe, and relief — and how these states evolve as events unfold — remains an outstanding challenge in affective neuroscience [2–5]. Despite divergent perspectives on the exact architecture of emotion, theories of emotion converge on a common view: emotions are not static, isolated reactions, but intrinsically dynamic and multidimensional phenomena that continuously adapt across time and context, enabling organisms to respond to and act upon changes in both their internal and external environments [6–9]. Different theoretical frameworks characterize this complexity in distinct ways. Discrete-emotion theories emphasize recurring affective categories with characteristic expressive and response profiles [10, 11]; dimensional models organize affect along continuous axes, most prominently valence (how pleasant or unpleasant a feeling is) and arousal (the level of physiological activation or energy) [12–17]; appraisal theories emphasize coordinated changes in evaluation, physiology, action tendencies, and subjective feeling [2, 6, 18, 19] ; and constructionist accounts emphasize the context-dependent conceptualization of core affective states generated from internal bodily states [20–23].

For decades, affective neuroimaging has approached emotion through brief, controlled snapshots: a fearful face, an aversive image, a short evocative clip [24–28]. By measuring blood-oxygen-level-dependent (BOLD) signals — the standard hemodynamic fMRI proxy for local neural activity [29–32] — studies using these paradigms identified recurrent affect-related responses across the amygdala, insula, cingulate cortex, and medial prefrontal cortex [33–35]. Yet this regional convergence did not establish one-to-one mappings between emotions and brain areas: no single region is necessary or sufficient for a given emotion [18, 20, 36, 37]. In this context, emotional states are increasingly understood as arising from distributed, partly overlapping cortical–subcortical patterns rather than isolated regional signatures [36, 38–41]. The explanatory target therefore shifts from individual regions to the evolving coordination of context-sensitive, overlapping brain networks [4, 19, 23, 42].

To capture this multifaceted structure, affective neuroscience has increasingly turned to naturalistic film fMRI [43–46]. Films provide a continuous stream of audiovisual, social, and narrative information, allowing affective meaning to unfold through changing events, expectations, and interpretations rather than being reduced to isolated stimulus–response episodes [47–51]. This is particularly important for emotion: states with similar valence or arousal can differ profoundly in their appraisal, bodily, motivational, and social meaning. By aligning ongoing brain activity with continuous, multivariate descriptions of experience, film fMRI therefore brings measurement closer to real-life environments, where emotional experiences are dynamic, context-dependent, and high-dimensional [43, 46].

This shift has moved the field beyond regional activation to dynamic functional connectivity [52–54]. Dynamic interactions among large-scale brain systems track continuously rated emotion, narrative tension, and moment-to-moment arousal across films and participants [54–59]. Edge-centric methods [60, 61] further decompose connectivity into dynamic pairwise co-fluctuations, revealing temporally sparse and behaviorally relevant patterns of coordination [62, 63]. Yet, both conventional and edge-centric connectivity remain dyadic: they describe how two regions relate at a time, not whether distinct collective states emerge only through the joint organization of several regions.

Higher-order network approaches [64–67] provide a natural framework for modeling dependencies among three or more regions, revealing collective states that remain opaque to regional and pairwise analyses. Information-theoretic frameworks distinguish between redundant information, which is shared across regions, and synergistic information, which emerges and is accessible exclusively through their joint activity [68–72]; topological approaches, in turn, characterize the collective shape of group interactions and their persistent mesoscale organization [1, 73–75]. Together, these methods have begun to reveal robust signatures in consciousness [70, 76, 77], learning [78], individual differences [75, 79], development and aging [80–82], as well as in neuromodulation and clinical conditions [83–89]. Crucially, topological higher-order metrics have shown to outperform traditional node- and edge-based methods in task decoding, and brain–behavior prediction [75, 90]. Yet, whether this representational power extends to the rich landscape of naturalistic emotion remains unknown, despite theoretical predictions of reconfigurable architecture made by componential models [2, 6, 19]: do collective brain dynamics reveal affective structures that regional and pairwise measures overlook — or does that advantage collapse when emotion is compressed into the classical, low-dimensional core axes of valence and arousal?

Here, we address these questions using the Emo-FilM dataset [91] (Figure 1**a**), which pairs naturalistic fMRI responses of 30 participants with continuous, multicomponent annotations of 50 distinct affective features [91, 92]. By systematically comparing regional BOLD activity, pairwise co-fluctuations, and higher-order topological descriptors (triangles and the homological scaffold; Figure 1**b**), we demonstrate that higher-order topology [1, 75] most accurately tracks recurrent emotional states and predicts the full, high-dimensional affective profile in held-out films. Crucially, this topological advantage is specific to the full affective profile: once affect is reduced into broad dimensions of valence, arousal, and power, the advantage diminishes, as pairwise connectivity is sufficient to track the dominant arousal signal. Testing these models across three independent film fMRI datasets [59, 93, 94], we find that pairwise connectivity robustly transfers arousal predictions to unseen narratives. Notably, no representation reliably generalizes valence, even though the relative geometry of affective states in representational space remains aligned across datasets. Generalization therefore emerges at complementary levels: higher-order topology is well suited for fine-grained emotional differentiation, while connectivity preserves and transfers only broad arousal dynamics across narratives.

**Figure 1.**
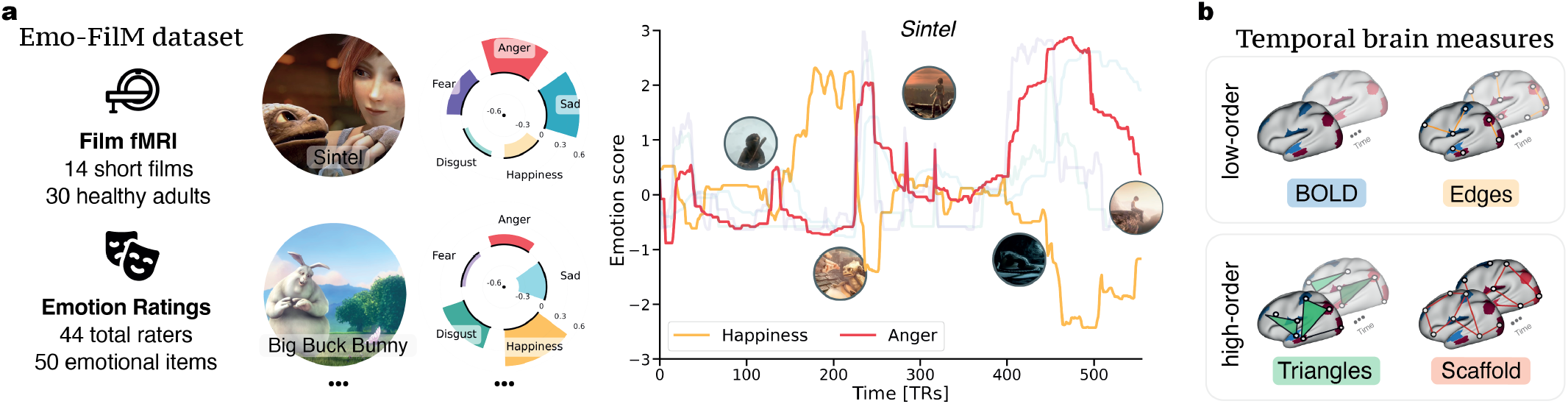
Emo-FilM dataset and time-resolved brain connectivity measures. **(a)** The Emo-FilM dataset comprises fMRI recordings from 30 healthy adults viewing 14 short films, paired with continuous annotations (44 raters) across 50 affective features: 13 discrete emotions and 37 component-process descriptors spanning appraisal, motivation, expression, bodily physiology, and subjective feeling. Average affective profiles differentiate films, as illustrated for *Sintel* and *Big Buck Bunny*, but compress the substantial emotional fluctuations that unfold within each narrative. Scene-by-scene trajectories of happiness and anger during *Sintel* illustrate this dynamic affective structure. **(b)** We tracked time-resolved neural dynamics at three complementary levels: regional BOLD signals reflect local activity; edge co-fluctuations capture transient pairwise coordination between regions; whereas a time-resolved topological framework identifies higher-order coordination across groups of regions. In the latter, hypercoherent triangles identify three-region co-fluctuations beyond their pairwise links, whereas the homological scaffold highlights edges that recurrently support the evolving mesoscopic organization of coherent and incoherent dynamics [1].

## RESULTS

Emotional narratives do not unfold as a sequence of isolated affective events. As a story progresses, feelings, appraisals, bodily responses, and action tendencies recur, recombine, and change, creating a structured but continuously evolving emotional landscape. To determine whether brain dynamics mirror this componential organization, we leveraged the Emo-FilM dataset [91], which features fMRI data from 30 participants viewing 14 short films. An independent cohort of 44 raters provided high-temporal-resolution continuous annotations of each film along 50 affective features, spanning discrete emotion labels and component-process descriptors of appraisal, motivation, expression, bodily physiology, and subjective feeling (see Methods for details on the dataset and Supplementary Figs S1-S3 for annotations).

### Temporal dynamics of emotions and brain activity

As individuals watch a film, specific emotional states naturally arise and change, often recurring across different scenes. We first ask whether moments sharing a similar multivariate emotional configuration mirror analogous neural configurations, even when separated by substantial temporal gaps. To test this, for each film, we describe every time point using its full 50-dimensional affective profile and correlate these profiles across the narrative. This generates an emotion time–time correlation matrix, or recurrence plot [95], in which each entry reflects Pearson’s similarity between the emotional states expressed at two points in the narrative (Fig. 2**a**).

**Figure 2.**
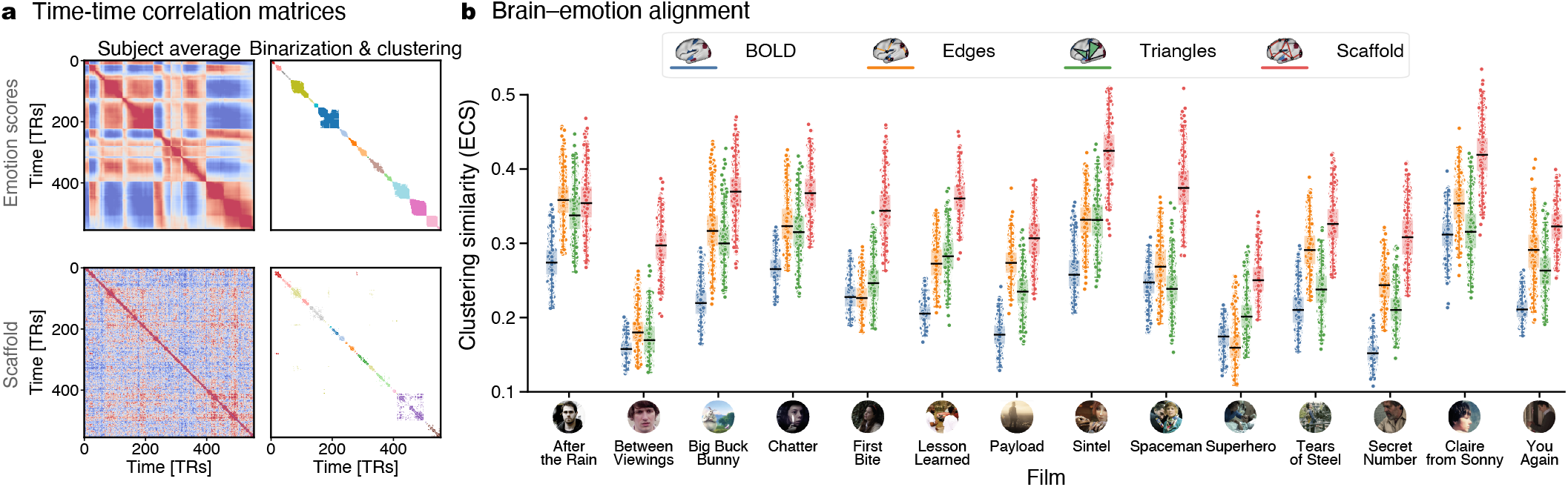
Time-resolved alignment between brain and emotion dynamics. **(a)** For each film, time–time correlation (recurrence) matrices captured emotional and neural dynamics. The emotion recurrence matrix quantifies the similarity between affective profiles across all pairs of time points, whereas brain matrices quantify similarity between instantaneous neural patterns over time. Averaging across subjects, binarizing (95th percentile), and applying Louvain clustering allowed the extraction of recurrent temporal states, yielding a common representation of emotion and brain dynamics. **(b)** Brain–emotion alignment was quantified via element-centric similarity (ECS) between the emotion- and brain-derived partitions for regional activity (BOLD), edge co-fluctuations (edges), hypercoherent triangles (triangles), and the homological scaffold (scaffold). Points denote 1000 paired participant subsamples (*N* = 25 of 30, matched across representations). The scaffold achieved the highest mean ECS in 13 of 14 films (one-sided exact binomial sign test, *p* = 9.2 *×* 10^*−*4^; proportion = 0.93), matched only by edge time series in *After the Rain*. Crucially, the scaffold exceeded all alternative representations in a median of 94% of subsamples, demonstrating robustness to participant composition. Boxes indicate the median and interquartile range.

To link these affective dynamics to brain activity, we parcellate the fMRI data into *N* = 119 regions, comprising 100 cortical parcels [96] and 19 subcortical and cerebellar regions [97]. We then construct analogous time– time similarity matrices from four complementary representations of brain dynamics. While regional BOLD activity describes local fluctuations across these *N* regions, the remaining three representations share a common edge-level space of 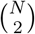 features, allowing direct comparison between pairwise and higher-order descriptions (see Methods for details). Edge time series [60, 61] capture moment-to-moment co-fluctuations between region pairs, whereas a time-resolved topological framework [1, 75] models collective coordination beyond pairs. Within this approach, hyper-coherent triangles isolate three-region interactions exceeding their constituent edges, and the temporal homological scaffold identifies connections that are part of the mesoscale network’s topology, namely, 1-dimensional cycles (see Methods and Supplementary Fig. S4 for a schematic representation).

For each neural representation, the resulting time– time matrix captures when similar whole-brain configurations recur over the course of a film. Following previous work [1], we retain the strongest temporal similarities by thresholding each matrix at its 95th percentile and use consensus Louvain clustering to identify recurrent temporal communities (see Methods for details, and Supplementary Fig. S5 for robustness analysis). These communities group moments with similar affective or neural configurations, regardless of their temporal distance in the narrative. We then quantify brain–emotion alignment using element-centric similarity (ECS) [98], a measure that compares two partitions and ranges from 0 for entirely dissimilar partitions to 1 for identical ones [98]. In other words, higher ECS values indicate a closer match between the recurrent temporal structure of a neural representation and that of the film’s evolving affective content.

Across films, the scaffold approach most closely mirrors the recurrent temporal structure of emotion (Fig. 2**b**). It achieves the highest mean ECS (0.345 *±* 0.047, mean ± s.d. across films), exceeding edge time series (0.279 *±* 0.059), triangles (0.264 *±* 0.052) and regional BOLD activity (0.221 *±* 0.047). The scaffold ranks first in 13 of 14 films (one-sided exact binomial sign test, *p* = 9.2 *×* 10^*−*4^; proportion = 0.93, one-sided 95% Clopper–Pearson lower bound = 0.70 ). This finding is also stable across participant subsamples; across 1000 paired resampled sub-groups of 25 out of the 30 participants, the scaffold yields the highest ECS in a median of 94% of subsamples overall, and in at least 79% of subsamples for 13 of the 14 films (Fig. 2**b**). The only exception is *After the Rain*, where scaffold and edge time series were very similar (ECS, 0.356 versus 0.360; scaffold higher in 39% of sub-samples).

Thus, while local regional activity and pairwise co-fluctuations capture a fraction of the temporal organization of emotion, it is the persistent mesoscale structure highlighted by the scaffold that most consistently decodes it. Crucially, this scaffold sensitivity is tuned specifically to affective, rather than merely audiovisual, organization. When applying the identical temporal-recurrence framework to low-level audiovisual features of the films —such as audio, video, or chromatic variations— the scaffold’s advantage over regional activity largely disappears (see Supplementary Fig. S6). This suggests that higher-order topology tracks the narrative’s complex emotional landscape rather than its stimulus stream.

### Multidimensional emotion–brain alignment

Knowing when emotional states recur does not explain what they contain. To map the intrinsically multidimensional nature of emotion onto brain dynamics, we rely on partial least squares correlation (PLSC) [99]. This is a multivariate approach that identifies paired latent variables (LVs) — weighted patterns of emotion and brain features — whose expression covaries maximally across observations. Each LV thus captures a distinct mode of shared emotion–brain variation. We operationalize this using the 190 film scenes from the consensus clustering of the emotion recurrence matrix across all 14 films. Summarizing each scene by its mean 50-dimensional affective profile and corresponding mean brain pattern, we fit an independent PLSC model for each neural representation. We then assess the number of significant LVs using permutation testing (5000 permutations) and quantify the reliability of individual loadings via a bootstrap ratio (BSR) across 5000 resamples, defining a reliable loading [99] as having |BSR| *≥* 2.0 (see Methods for details). This analysis reveals pronounced differences in how the four representations encode affective dimensionality (Fig. 3). Regional BOLD activity covaries with emotional content through a *single* significant latent variable, which alone captures 49.7% of the emotion–brain covariance. Moving from local activity to network connectivity does not simply strengthen this relationship but unfolds it. Edge time series leads to three significant LVs (43%, *p* = 0.019; 27%, *p* = 0.009; 9%, *p* = 0.001; a total of *∼*79% covariance), while the homological scaffold has four (38%, 20%, 9%, and 6%; all *p <* 0.05; total of *∼*74%). Hypercoherent triangles, by contrast, capture only a fragmented fraction of reliable covariance spread across two LVs (roughly 12% in total - LV2 and LV3). The loadings that define these latent variables reveal a striking spatial dissociation (Fig. 3**b**,**c**). BOLD saliences concentrate in limbic and subcortical regions, consistent with their canonical role in affective processing [20, 33, 35, 38] (see Supplementary Fig. S7 for details on brain regions as a function of different bootstrap ratios). Conversely, connectivity-based representations distribute their loadings across a broad architecture of interacting cortical and subcortical networks, with each significant LV pairing a distinct blend of affective features to specific network-level signatures.

**Figure 3.**
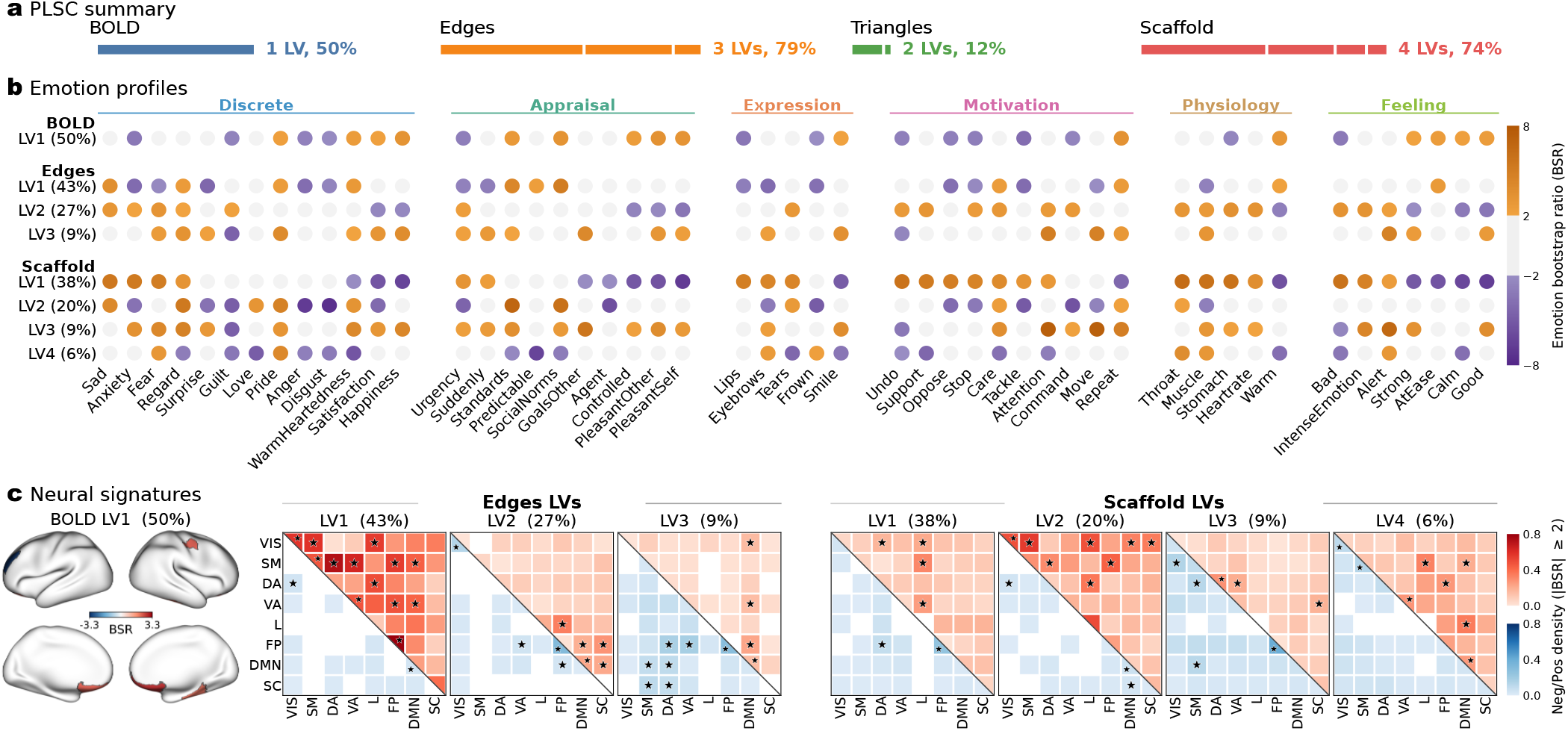
Connectivity representations reveal multidimensional associations between emotional experience and brain dynamics. **(a)** Partial least squares correlation (PLSC) relating emotion-defined scenes to their mean brain patterns for four representations: regional BOLD activity, edge time series (edges), higher-order triangles, and the temporal homological scaffold. Scenes were defined by consensus Louvain clustering of the emotion time-by-time correlation matrix and concatenated across the 14 films. Bars indicate the number of significant latent variables identified for each representation (LVs; permutation test, 5000 permutations, *p <* 0.05) and the covariance explained by these LVs. Regional BOLD activity yielded a single significant mode, whereas the connectivity-based representations resolved several (three for edges and four for the scaffold), revealing a richer, more differentiated emotion–brain mapping. Because cumulative covariance depends on the scene partition, significance and explained covariance are reported per LV. **(b)** Emotion saliences for each significant LV across the 50 affective features, grouped into discrete emotions, appraisals, expressions, motivations, physiological responses and feelings. Marker color denotes the bootstrap ratio (BSR; salience divided by its bootstrap standard error, 5000 iterations); gray markers indicate unreliable saliences ( |BSR| *<* 2.0). **(c)** Corresponding brain saliences. For regional BOLD activity, cortical surface maps show reliable regional saliences ( |BSR| ≥ 2.0). For edges, triangles and the scaffold, 8 *×* 8 matrices give the density of reliable connections within and between seven canonical cortical resting-state networks and subcortical regions, with positive and negative saliences in the upper (red) and lower (blue) triangles; asterisks mark network pairs whose density of reliable connections exceeds chance (one-sided hypergeometric test, Benjamini–Hochberg FDR *<* 0.05). BOLD saliences concentrate in limbic and subcortical regions, whereas the connectivity modes embed these regions within a broader cortico-subcortical architecture, most extensively for the scaffold.

Scene definition establishes the fundamental units of this analysis, and certain metrics inherently depend on this choice. Because total covariance fluctuates across partitions and the number of connectivity modes scales with the number of scenes, we evaluate significance and covariance mode by mode rather than reporting a cumulative sum. Crucially, while these absolute totals vary, the dimensional contrast between neural representations remains qualitatively stable. Across distinct scene definitions, regional activity resolves exactly one significant mode, while both connectivity representations consistently resolve several. Furthermore, this structural dichotomy survives a conservative, film-restricted permutation scheme. By explicitly testing whether emotion–brain coupling exceeds what film-level differences alone can produce, this null model confirms that the multidimensional architecture of affective connectivity is a robust phenomenon (Supplementary Fig. S8 and Supplementary Note B). Importantly, this divergence is not driven by mere feature count [100]: despite operating within a similar high-dimensional space (7021 functional connections), hypercoherent triangles yield the fewest modes of any connectivity representation.

### Higher-order brain topology decodes emotional richness

Alignment within a single film does not guarantee prediction in a new one. Moving from description to out-of-sample prediction, we ask whether a model trained on 13 films can recover the continuous trajectory of emotional features in a held-out film. We decode each of the 50 emotion descriptors time-courses using kernel ridge regression (KRR). Because KRR operates on the similarity structure among observations rather than requiring explicit feature selection, it places the 119 regional BOLD signals and the 7021 edge-level features on a common footing [101]. To focus on sustained affective dynamics rather than transient frame-to-frame fluctuations, we apply identical Gaussian-weighted temporal smoothing to both neural and emotion time series (*σ* = 15 TRs, approximately 19.5 s) and score accuracy as the Pearson correlation between predicted and observed trajectories in the held-out film (see Methods; robustness to smoothing width in Supplementary Fig. S9).

At the full resolution of all 50 fine-grained affective descriptors, the homological scaffold emerges as the most accurate predictor for most emotion regressors (Fig. 4**a**-**b**). Averaged across films and emotion targets, scaffold-based prediction achieved the highest mean accuracy 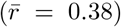, outperforming triangles 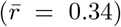, pair-wise edge time series 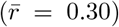, and regional BOLD activity 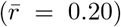. Because the 50 targets are partially correlated, we compare representations at the level of held-out films, with each film contributing its complete affective profile as a single multivariate outcome (see Methods). The scaffold shows the highest overall prediction accuracy. It exceeded regional BOLD activity in every film (Δ*r* = 0.181, *d*_*z*_ = 1.52, *p*_FDR_ *<* 0.001) and outperformed edge time series in 11 of 14 films (Δ*r* = 0.083, *p*_FDR_ = 0.022). Triangles and edge time series also improved substantially over regional BOLD, whereas differences among the interaction-based representations were smaller: the scaffold did not reliably exceed triangles, and edges and triangles performed similarly. As a conservative sensitivity analysis accounting for overlap among cross-validation folds, we additionally applied the Nadeau–Bengio correction; under this stricter test, only the scaffold–BOLD contrast remained significant. Thus, the results robustly support an advantage of higher-order and connectivity-based representations over regional activity, with the scaffold showing the strongest overall performance.

**Figure 4.**
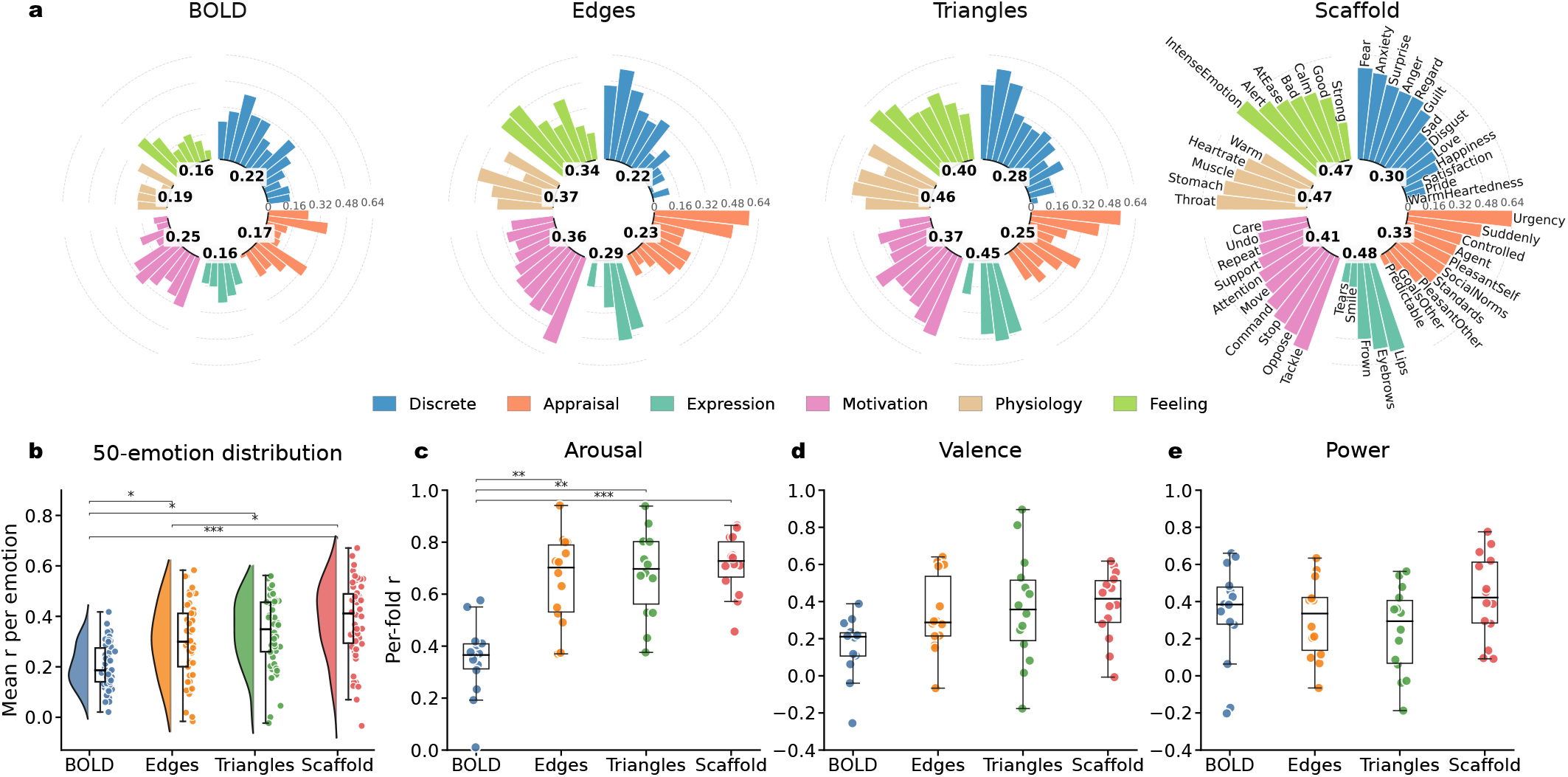
Emotion granularity shapes decoding performance across feature spaces. **(a)** Prediction accuracy for 50 fine-grained emotions across four brain representations: regional BOLD activity, edge time series (Edges), triangles, and the homological scaffold. Accuracy (Pearson’s *r*) was evaluated via kernel ridge regression (leave-one-film-out cross-validation, 15-TR Gaussian smoothing). Colors denote six emotion components; inner labels report median absolute *r*. For visual clarity, features are ordered within each component by Scaffold accuracy across all plots, with labels shown only for the Scaffold. **(b)** Aggregate decoding accuracy across all 50 emotion descriptors (*n* = 50). To account for interdependence among emotion trajectories, statistical brackets test across the 14 held-out films (exact sign-flip test, FDR-corrected; see Methods). At this fine-grained scale, the scaffold exceeded BOLD in every film (*d*_*z*_ = 1.52) and outperformed edges (*d*_*z*_ = 0.78, 11 of 14 films). Edges and triangles each exceeded BOLD (*d*_*z*_ = 0.81 and 0.89, respectively). **(c–e)** Decoding performance when emotions are compressed into broad dimensions of arousal, valence, and power (points are film folds, *n* = 14). For arousal, all connectivity-based representations outperformed regional BOLD, but the scaffold no longer showed an advantage over edges or triangles (Nadeau–Bengio corrected paired *t*-tests across folds, FDR-corrected; scaffold vs. BOLD: *d* = 2.67; edges vs. BOLD: *d* = 1.53; scaffold vs. edges: *d* = 0.44, n.s.). For valence and power, no representation showed a reliable predictive advantage. Boxplots show the median, interquartile range, and 1.5*×* interquartile range. Significance markers indicate FDR-corrected tests: ^:^*p <* 0.05, ^::^*p <* 0.01, ^:::^*p <* 0.001. Notice that in panels **(c)-(e)** a single ridge penalty was selected using arousal across all films and then held fixed across folds and dimensions (see Methods).

The picture changes when the 50 descriptors are summarized by the three canonical affective dimensions of valence, arousal, and power [92, 102] (see Methods) (Fig. 4**c–e**). Arousal is decoded well by all three connectivity representations, which significantly exceed regional BOLD performance (Nadeau–Bengio corrected resampled *t*-tests across folds [103], *p*_FDR_ *<* 0.05; scaffold versus BOLD Cohen’s *d* = 2.67, edges versus BOLD *d* = 1.53). However, the scaffold no longer outperforms the pairwise edge time series (*d* = 0.44, not significant). For valence and power, no neural representation predicts significantly better than any other (all *p*_FDR_ *>* 0.05).

Thus, the scaffold’s sensitivity does not reflect a globally superior decoding of all brain activity; this advantage dissolves once affective content collapses onto a broad, arousal-dominated signal that pairwise coupling already captures. Compressing neural representations rather than emotional ones yields a similar effect. The scaffold’s margin over edge time series is largest under the full-rank kernel model and diminishes as the neural feature space is reduced toward low-dimensional summaries, indicating that its predictive power rests on a highly distributed code (Supplementary Fig. S10).

### Distributed coding and the anatomy of arousal

Our predictive decoding results indicate that the sensitivity of the homological scaffold is not a generic improvement in prediction, but depends fundamentally on which affective content is being predicted. Higher-order topology is most informative when emotion is described as a rich, differentiated profile, whereas broad arousal is readily captured by pairwise connectivity. To understand this dissociation, we examine the dimensionality, overlap, and anatomical organization of these emotion-related representations (Fig. 5).

**Figure 5.**
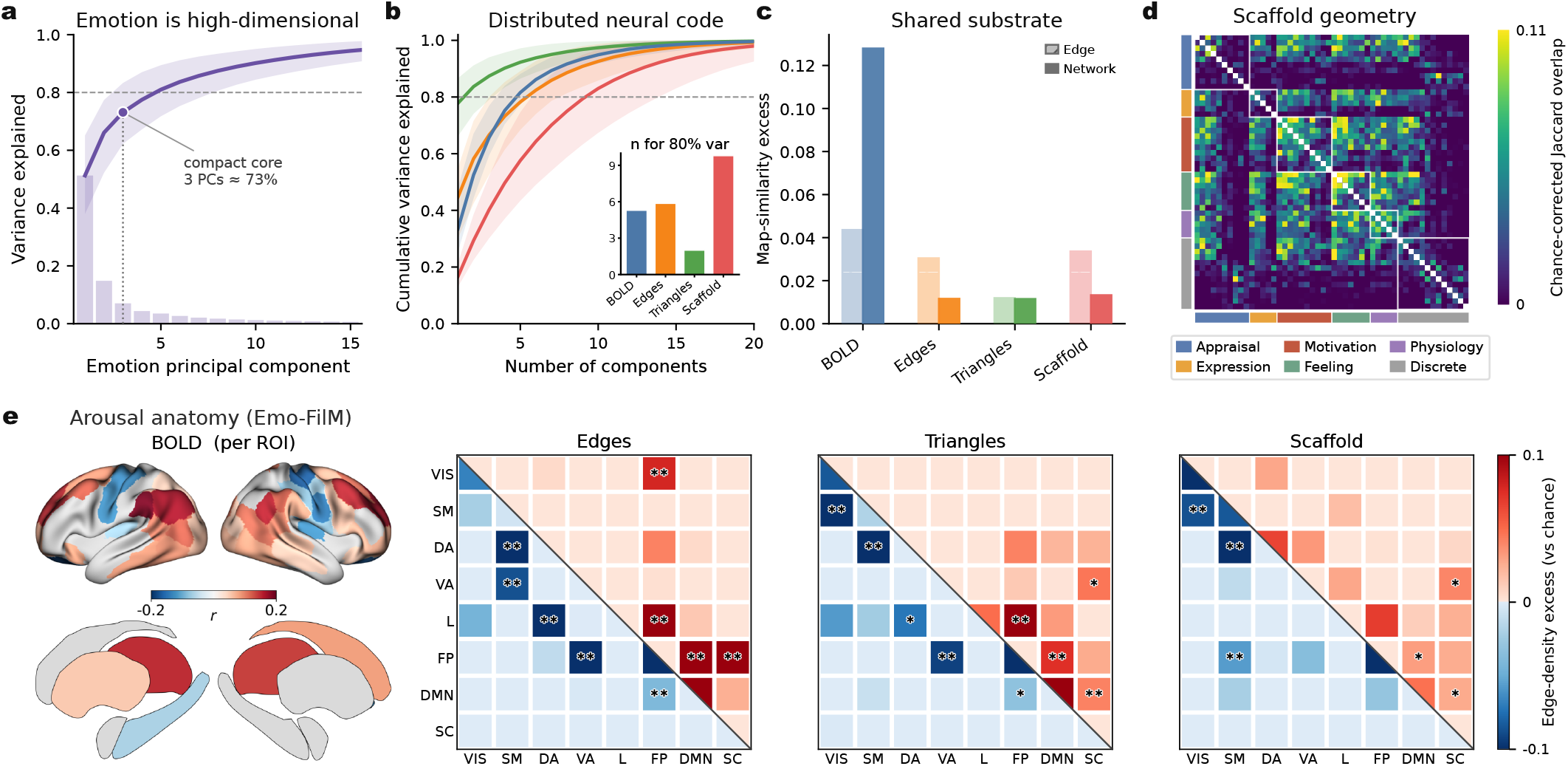
Dimensionality and spatial organization of emotion-related brain representations. **(a)** Principal-component decomposition of the 50 continuous emotion ratings, computed separately within each film. Bars show the across-film mean variance explained, the line shows cumulative variance, and the shaded band denotes variability across films (2.5-97.5 percentile range). The first three components capture ∼73% of the variance, but substantial residual structure extends across many additional components. **(b)** Cumulative variance explained by the principal components of each brain representation (shaded bands denote ±1 standard deviations across films). The inset reports the number of components required to explain 80% of the variance (5, 6, 10, and 2 for BOLD, edges, scaffold, and triangles, respectively). Triangles collapse onto a single dominant mode, whereas the scaffold shows the most distributed spectrum. **(c)** Similarity between emotion-predictive maps for each brain representation. For each pair of emotions, cosine similarity was computed between their selected feature maps in the native feature space (hatched bars) and after aggregation to the eight cortical network/subcortical systems (solid bars). Bars show mean cosine-similarity excess relative to a phase-randomized emotion-label null, preserving temporal autocorrelation and cross-emotion covariance. **(d)** Chance-corrected Jaccard overlap between scaffold edge sets selected for each pair of emotion components, ordered by component family. Overlap partly follows family-wise structure, while also extending across families, suggesting organization beyond strict emotion-family membership. **(e)** Anatomical distribution of arousal-related information in Emo-FilM. BOLD maps show regional decoding contributions on cortical and subcortical surfaces. ETS, scaffold, and triangles are summarized as 8 *×* 8 density-excess matrices over seven canonical resting-state networks and subcortical regions. Each cell represents the excess proportion of selected edges between a pair of systems relative to a hypergeometric chance baseline. The upper triangle shows positive arousal-predictive edges, whereas the lower triangle shows negative arousal-predictive edges. Asterisks denote significant network-pair enrichment after FDR correction (*:p*_FDR_ *<* 0.05, *: : p*_FDR_ *<* 0.01).

We first characterize the intrinsic dimensionality of the affective targets by applying principal component analysis (PCA) to the 50 continuous emotion-rating time courses of each film separately (see Methods for details). The first three components explain approximately 73% of the variance on average (61–84% across films). However, the valence, arousal, and power factors are largely captured by the first two components alone. Because the factors are naturally correlated whereas principal components are orthogonal, this correspondence operates as a shared subspace rather than a one-to-one mapping across individual axes. However, the remaining 27% of the variance does not collapse onto a single additional axis; rather, it is distributed across a tail of components (Fig. 5**a**). Thus, while the 50-emotion space contains a compact low-dimensional core, it also possesses substantial residual structure that preserves highly differentiated affective content.

We next ask whether the neural representations preserve or compress this distributed structure. Applying PCA to the participant-averaged feature time series within individual films, we determine the number of components required to explain 80% of the variance. Regional BOLD activity and edge co-fluctuations reach this threshold with approximately five to six components, whereas triangles are dominated by two modes. By contrast, the homological scaffold exhibits the broadest spectrum, requiring approximately ten components to explain 80% of the variance (Fig. 5**b**). This distributed geometry provides a natural explanation for the decoding results: the scaffold retains information across multiple weakly dominant modes. This architecture is advantageous for predicting the full 50-emotion profile, yet becomes less distinctive once the target space is reduced to broad dimensions like arousal.

This distributed code is not a generic affective substrate reused identically across all emotion regressors. To quantify the extent to which different emotions share neural features, we construct emotion-specific encoding maps by ranking features (i.e., regions or edges) according to their correlation with each emotion’s time course on the training films. This correlation-based feature selection closely builds upon the logic of canonical connectome-based predictive modeling (CPM) [104, 105]. However, rather than relying on a traditional uncorrected *p <* 0.01 significance threshold, we deliberately retain the strongest 10% of associations within each cross-validation fold to guarantee a stable, sparsity-controlled feature set, and subsequently intersect these selections across all 14 folds. Because the emotion ratings are themselves correlated and temporally autocorrelated, calculating raw overlap would overestimate shared neural structure. We therefore express map similarity as an excess over a phase-randomized emotion-label null that preserves both temporal autocorrelation and cross-emotion covariance while breaking the true brain–emotion correspondence (see Methods for details). Under this stringent correction, different emotions exhibit only a modest shared substrate, which is strongest at the network level for regional BOLD activity and substantially weaker for the connectivity representations (Fig. 5**c**). For the scaffold, the chance-corrected Jaccard overlap between selected edge sets organizes into family-wise blocks corresponding to appraisal, feeling, physiology, and discrete-emotion categories, with some occasional extensions across families, rather than forming a single global affective pattern (Fig. 5**d**). These results indicate that the scaffold’s sensitivity arises from a differentiated relational code: while some emotions reuse common architecture, much of their predictive information is carried by distinct configurations of network interactions.

Against this high-dimensional background, arousal provides a uniquely informative special case. It is the dimension most consistently decoded by all connectivity representations, and its spatial organization has previously been linked to broad frontoparietal and default-mode interactions during naturalistic viewing [59]. Because the raw backward weights of decoding models are not directly interpretable under severe feature collinearity [106], we localize arousal-related information using the identical forward/encoding mapping approach described above rather than the kernel-ridge-regression weights themselves (see Methods for details). Applying this stable, sparsity-controlled selection (intersecting the top 10% of positive and negative associations across all 14 folds; for regional BOLD, a per-fold *p <* 0.01 criterion, see Methods), we summarize the resulting arousal-predictive connectivity maps as a network-pair density excess relative to a hypergeometric baseline. This explicitly identifies network pairs enriched beyond what would be expected from their size alone (Fig. 5**e**).

Across connectivity representations, arousal reveals a partially shared network architecture: positive associations involve the default-mode network and its coupling with frontoparietal and subcortical systems — for edge time series and triangles most prominently a limbic–frontoparietal link — whereas negative associations are dominated by within-frontoparietal coupling together with somatomotor–attentional and visual interactions (Fig. 5**e**). This convergence is markedly weaker for valence and power (Supplementary Fig. S11). These data indicate that broad arousal follows a reproducible connectivity signature, aligning with existing results [59]. The encoding maps also closely matched the decoder’s forward activation patterns, unlike its raw regression weights (Supplementary Fig. S12) [106]. Figure 5 thus reveals complementary scales of affective organization: the scaffold differentiates fine-grained emotional content, whereas arousal recruits a more reproducible global network architecture.

### Cross-dataset validation

To test whether these findings capture a generalizable architecture of emotion rather than responses specific to Emo-FilM, we extend the analysis to three independent film fMRI datasets: Sherlock [93], Merlin [94], and Jojo Rabbit [107]. These datasets differ substantially in narrative structure, duration, acquisition parameters, and affective annotation methods, precluding direct temporal alignment across films. We therefore focus on valence and arousal, the affective dimensions shared across datasets, and compare their neural representations using representational similarity analysis (RSA) [108] (Fig. 6**a** for a schematic; see Methods). The datasets differ substantially in their affective distributions, with valence–arousal correlations ranging from *r* = −0.59 in Emo-FilM to *r* = 0.12 in Sherlock (Fig. 6**b**).

**Figure 6.**
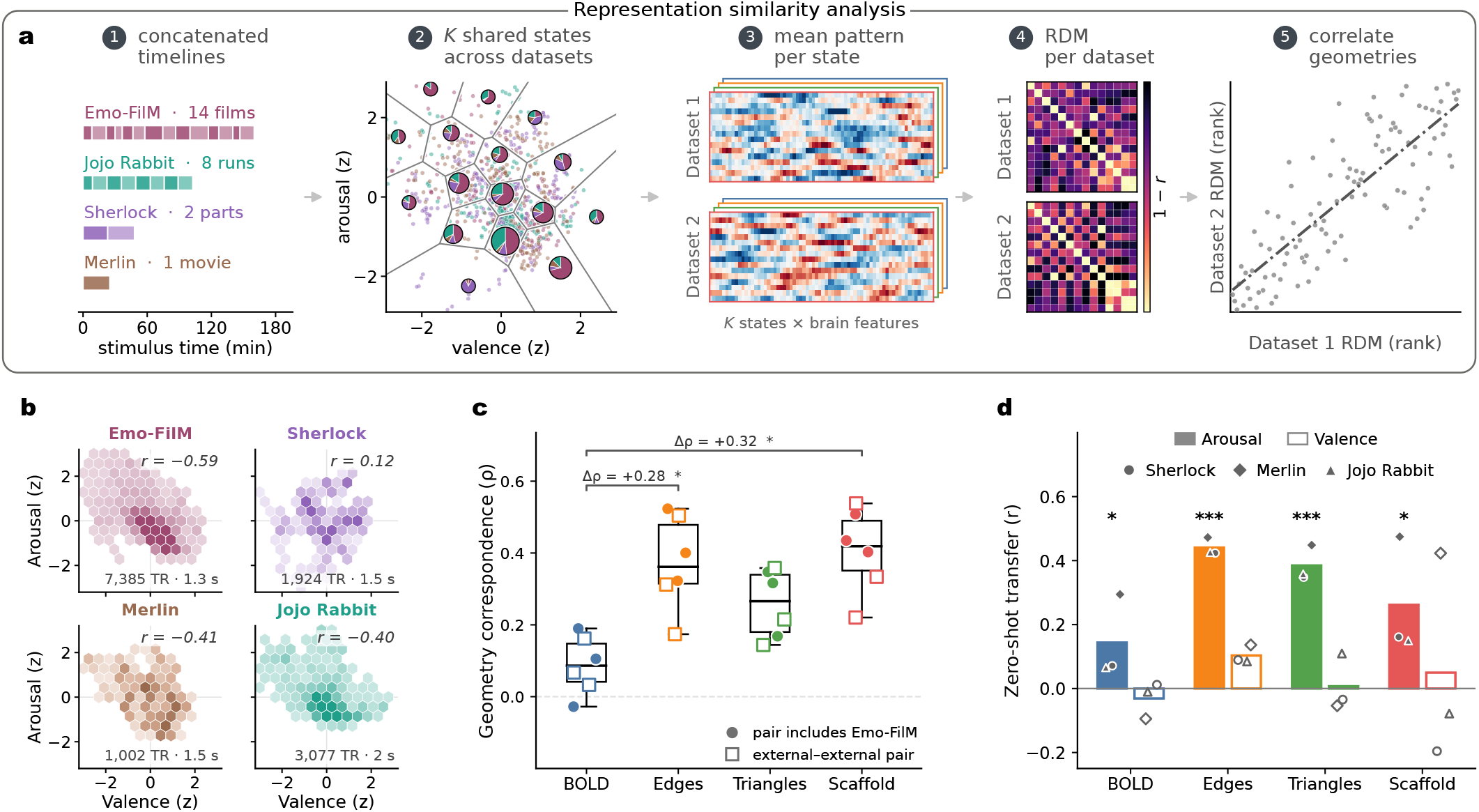
Cross-dataset generalization of affective representations and their geometry. **(a)** Cross-dataset representational similarity analysis. Because the four datasets differ in stimulus structure, duration, and acquisition, their neural time series cannot be aligned directly. Valence and arousal are therefore standardized within each dataset and partitioned into *K* = 16 shared affective states. For each dataset and neural representation, activity is averaged within each sufficiently sampled state, and pairwise correlation distances (1 *− r*) define a representational dissimilarity matrix (RDM). Cross-dataset correspondence is quantified as the Spearman correlation (*ρ*) between matched RDMs. **(b)** Distribution of standardized valence and arousal ratings across datasets. Hexagonal densities show state occupancy; annotations report the number of fMRI time points and TR for each dataset. The different distributions and correlations (*r* = −0.59 to 0.12) illustrate the affective domain shift across narratives. **(c)** Cross-dataset correspondence of affective-state geometry for regional BOLD, Edges, Triangles, and Scaffold representations. Points show the six dataset pairs (circles, comparisons involving Emo-FilM; squares, external–external comparisons), and box plots summarize their distribution. Edges and Scaffold showed greater correspondence than regional BOLD (Edges: Δ*ρ* = 0.28, *p*_*FDR*_ = 0.036; Scaffold: Δ*ρ* = 0.32, *p*_*FDR*_ = 0.024). **(d)** Zero-shot affect prediction from models trained exclusively on Emo-FilM and applied without refitting to Sherlock, Merlin, and Jojo Rabbit. Bars show the mean transfer across external datasets for arousal (filled) and valence (open), with symbols indicating individual datasets. Arousal generalized for edges (*r* = 0.44, *p*_*FDR*_ *<* 0.001), triangles (*r* = 0.38, *p*_*FDR*_ *<* 0.001), scaffold (*r* = 0.26, *p*_*FDR*_ = 0.016) and BOLD (*r* = 0.14, *p*_*FDR*_ = 0.032), whereas valence did not significantly transfer for any representation. Together, panels **(c**,**d)** assess the preservation of affective-state geometry and the zero-shot transfer of individual affective dimensions across datasets.

We first ask whether the relative geometry of affective brain states is preserved across film narratives. Within each dataset, time points are assigned to *K* = 16 shared states defined in standardized valence–arousal space. Neural activity is averaged within each sufficiently sampled state (i.e., at least 10 fMRI volumes), and pairwise correlation distances between state patterns yield a representational dissimilarity matrix (RDM) for each neural representation. Correlating RDMs across datasets reveals stronger geometric correspondence for connectivity-based representations than for regional BOLD (Fig. 6**c**).

At *K* = 16, both edges and scaffold show greater cross-dataset correspondence than BOLD (edges: Δ*ρ* = 0.28, *p*_FDR_ = 0.036; scaffold: Δ*ρ* = 0.32, *p*_FDR_ = 0.024). This hierarchy is robust across state resolutions from *K* = 12 to 32 (Supplementary Fig. S13).

We next test the more stringent requirement of direct zero-shot prediction. Kernel ridge regression models trained exclusively on Emo-FilM are applied, without refitting, to Sherlock, Merlin, and Jojo Rabbit (Fig. 6**d**). Arousal generalizes most strongly for edges (*r* = 0.44, *p*_FDR_ *<* 0.001), followed by triangles (*r* = 0.38, *p*_FDR_ *<* 0.001), scaffold (*r* = 0.26, *p*_FDR_ = 0.016) and regional BOLD (*r* = 0.14, *p*_FDR_ = 0.032). In contrast, valence does not significantly transfer for any representation (all *p*_FDR_ *>* 0.32). This finding is not an artifact of poor data quality, as valence ratings actually proved more reliable than arousal ones (split-half *r* = 0.94 versus 0.93 in Emo-FilM, 0.93 versus 0.84 in Sherlock, and 0.79 versus 0.73 in Merlin).

Although significant, scaffold zero-shot arousal transfer remains weaker than for the other connectivity representations, potentially reflecting the limited diversity of Emo-FilM training. We therefore repeated the analysis using leave-one-dataset-out training, fitting models on three narratives and testing on the fourth. Broader training improved arousal transfer across all connectivity representations, including the scaffold, while regional BOLD changed little (Supplementary Fig. S14**a-b**). It also yielded weak apparent valence transfer for BOLD and edges; however, after removing variance shared with arousal, only arousal remained transferable across all representations (Supplementary Fig. S14**c**). Thus, greater narrative diversity enhances the portability of arousal-related connectivity structure without revealing valence transfer beyond arousal.

Additional analyses indicate that the edge-based arousal signal is distributed rather than confined to a small fixed feature set (Supplementary Fig. S15). Removing the top 25–250 Emo-FilM-ranked edges leaves transfer essentially unchanged, whereas excluding 4000 of 7021 edges (57%) reduces performance (Δ*r* = *−*0.120, 95% CI −0.219, −0.035). Consistently, network-pair edge– arousal profiles are conserved across datasets (mean *r* = 0.507, range 0.22–0.70; phase-randomized *p* = 0.001), with prominent frontoparietal contributions. This correspondence is not evident at the regional scale (*r* = 0.162), and removing all frontoparietal connections only modestly reduces transfer (*r* = 0.441 to 0.434). Thus, arousal generalizes through a distributed yet redundant configuration of interactions between functional systems rather than a small anatomical core.

Together, these analyses reveal complementary forms of generalization. Arousal transfers significantly across all connectivity representations, although somewhat more strongly for edges and triangles than for the scaffold. By contrast, both edges and scaffold robustly preserve the cross-dataset geometry of affective states, indicating that the scaffold generalizes more through relational structure than through a fixed arousal mapping.

## DISCUSSION

A central challenge in affective neuroscience is understanding how the brain coordinates distributed systems to generate the rich, dynamic content of a large variety of emotional experience [4, 5, 23]. Our results reveal that, compared to regional activation alone, both pairwise connectivity and the mesoscale topological organization of brain networks capture a much richer picture of emotional experience during dynamic naturalistic conditions [1, 73, 75, 109, 110]. By comparing four distinct neural descriptions across extensive film viewing, we found that pairwise and higher-order interactions capture complementary facets of affect. The temporal homological scaffold — representing evolving higher-order topology — acts as a highly sensitive lens for multidimensional emotion. It best tracks the temporal recurrence of fine-grained affective states, resolves the most emotion–brain covariance modes, and most accurately decodes the full 50-feature affective profile used to differentiate discrete emotion categories. However, this sensitivity is bounded: its advantage is clearest for the full affective description, capturing intricate representational structure rather than acting as a universally portable decoder. Indeed, pairwise edges performed comparably or better under some regimes, indicating that these two connectivity descriptions are complementary rather than redundant, and both markedly outperform standard BOLD activity.

A natural reading of these results might be that modeling interactions beyond pairs is inherently superior. Our data suggest otherwise. Hypercoherent triangles also constitute a higher-order description, yet their variance collapses onto essentially a single dominant mode, rendering them one of the weakest of the three connectivity representations throughout our analyses. The scaffold, by contrast, was the most distributed of all four representations, requiring on the order of ten components to account for the majority of its variance (Fig. 5). Consistent with this, the scaffold’s margin over pairwise edges is largest under the full-rank kernel model and was not detectable under estimators that select or compress features (Supplementary Fig. S10), suggesting that its predictive information is spread across many modes. Each representation thus excels in a specific regime — edges for robust, localized interactions, and the scaffold for structure distributed across many weakly dominant yet recurrent and coherent modes. Therefore, what distinguishes a useful higher-order representation is not strictly the order of the interactions it encodes, but the breadth of the representational space those interactions span.

The scaffold enabled effective decoding across the full 50-feature space but disappeared once affect was compressed into broad dimensions of valence, arousal, and power (Fig. 4). We interpret this as a neural signature of emotional granularity rather than a generic improvement in predictive power [2, 9, 111]: the scaffold preserved a differentiated experiential content discarded by broad affective axes, providing a more ecologically faithful characterization of emotion as a multidimensional phenomenon. No single mode dominated emotional representation in either behavioral data or cerebral activity; the representation that best predicted fine-grained affect also showed the broadest dispersion of variance. At reduced behavioral dimensionality, pairwise metrics performed well, consistent with low-dimensional emotion topographies in hippocampal and prefrontal systems reported in other studies [50, 102]. The additional decodable richness of the full emotional space may reflect a complementary level of organization across macroscopic networks [18, 20, 42]. Furthermore, the scaffold closely tracked the recurrent temporal structure of emotional states (Fig. 2), enabling us to examine when and how cerebral activity emerges in response to emotional stimuli. This provides novel topological evidence for accounts of affect as arising from distributed, system-level interactions [18, 40, 112–114], particularly through transient network synchronization [19, 54, 56–58, 105, 115], and aligns with brain-wide rather than strictly localized accounts of higher-order cognition [116].

Across three independent films (Sherlock, Merlin, and Jojo Rabbit), the two canonical affective dimensions behaved entirely differently. Models trained on Emo-FilM and applied without refitting successfully transferred arousal to unseen films, whereas no representation transferred valence (Fig. 6) — a divergence consistent with recent findings [59]. Annotation quality does not explain this asymmetry, as valence ratings actually proved more reliable than arousal ratings in the datasets where per-rater data was available. Instead, the two dimensions appear fundamentally distinct in their neural architecture. Arousal rests on a redundant, frontoparietal-anchored network where decoding accuracy survived the removal of substantial portions of the connectome. This aligns with characterizations of arousal as a widely distributed, multi-system neural reference space [26, 55, 63, 117–119], possibly governing attentional orienting and alerting functions through interactions with other emotion-related systems. In contrast, although observers show high behavioral consensus when rating valence, its underlying neural instantiation remains deeply context-dependent [120–124]. More generally, valence itself may be an overly coarse construct: positive–negative classifications obscure emotions’ distinct functions and contextual variability [125], and some accounts question whether valence is represented along a single bipolar dimension rather than by partly separable positive and negative systems [126, 127]. Even positive affect may fractionate into distinct processes such as “liking” and “wanting” [18, 128]. Together, these conceptual and neural distinctions may explain why an approach based on shared group-level representations captures the more universal dynamics of arousal but struggles to generalize the more context-dependent organization of valence across narratives, despite high behavioral agreement. However, lack of cross-film transfer in valence decoding does not imply an absence of shared affective structure. When testing whether different films preserved the relative arrangement of affective states, connectivity representations agreed across datasets significantly better than regional activity (Fig. 6**c** and Supplementary Fig. S13) [129, 130]. Shared structure may therefore persist beyond fixed anatomical loci or reproducible connections, instead emerging from the relational configuration of emotional states in representational space. This explains the scaffold’s profile: its within-dataset advantage relies on a mesoscale topology dynamically reorganized by each film narrative, meaning rigid feature-to-emotion mappings cannot port directly even if underlying geometries are preserved. Consequently, representational richness and portability appear partly separable. Directly testing this *post hoc* interpretation, however, requires manipulating film narrative structure rather than relying on these four existing datasets.

Several limitations should guide interpretation. Validating across independent datasets is both a central strength and a defining constraint of this work. Because these external film datasets vary fundamentally in narrative content, acquisition parameters, and annotation methods, establishing a common target space required reducing our rich annotations to broad dimensions of valence and arousal. Consequently, we can test the generalization of broad affective dynamics, but we cannot confirm whether the full 50-dimensional profile transfers across narratives. Furthermore, with only three external datasets, cross-dataset agreement remains qualitative rather than a robust population-level estimate. Additionally, our analyses deliberately utilized group-average approaches to isolate shared, overarching principles of affective organization. While effective for this purpose, this inherently averages substantial interindividual variability of emotional experience [131, 132]. Finally, the scaffold and pairwise edge representations were not reliably separable across several measures, reinforcing that higher-order and pairwise descriptions act as complementary, rather than strictly redundant or competing analytical lenses.

Overall, comparing regional activity to higher-order topology reveals that emotion is distributed across systems rather than locally concentrated. Our results demonstrate a division of labor across scales. Pair-wise connectivity captures the generalizable dynamics of arousal, which has a foundational role throughout, as high-arousal emotions were consistently the most robustly predicted. In contrast, higher-order topology best describes fine-grained, context-bound affective profiles presumably underlying the rich variety of human emotion experience. Ultimately, it is the relational geometry of affective states — not their anatomical localization — that generalizes across independent populations. This multi-scale framework is compatible with componential models of emotion [2, 18, 19] and more broadly accord with network [4, 42], psychological-constructionist [20, 23], and semantic-space accounts [9, 111], demonstrating that each holds at the level of description where it is measured. Mapping emotion concurrently across multiple scales thus offers a promising route to characterize human emotional experience beyond any single stimulus set [38, 133–136].

## METHODS

### Datasets and participants

We analyzed four independent naturalistic film fMRI datasets to track emotion-related neural dynamics during audiovisual film narratives and test their reproducibility across varying stimuli, samples, and annotation schemes. We selected Emo-FilM as the primary dataset for its fine-grained, multicomponent emotion annotations. To test generalizability, we validated our findings against three additional datasets featuring full-length films and television shows annotated for valence and arousal. All original studies obtained institutional ethics approval and written informed consent. As this work consists of secondary analyses, sample sizes were not statistically pre-determined.

#### Dataset 1 (Emo-FilM)

The Emo-FilM dataset [91] contains fMRI data from 30 adults (18 female; aged 18-34) who viewed 14 emotional short films across four randomized sessions, for a total of 2h 40min per participant. Affective targets were defined by an independent cohort of 44 behavioral raters (23 female; aged 20-39), yielding 50 continuous features spanning discrete emotion labels and componential measures (appraisal, motivation, expression, physiology, and feeling). The study was approved by the Geneva Cantonal Commission for Ethics in Research. Data are available on OpenNeuro (fMRI: ds004892; annotations: ds004872). For details, see the original study [91].

#### Dataset 2 (Jojo Rabbit)

To evaluate generalization to full-length film, we used the *Jojo Rabbit* dataset from the REELMO resource [107]. fMRI data were acquired from 20 adults (10 female; mean age 27 years) watching the full film across two 1-hour sessions (totaling 40 hours of data). Affective targets were derived from independent behavioral ratings from 47 viewers tracking 20 emotion labels. The study was approved by the Comitato Etico Area Vasta Nord Ovest (protocol 1485/2017). Data are available on Figshare (28255745).

#### Dataset 3 (Sherlock)

We analyzed an open fMRI dataset of 16 participants viewing a 48-minute segment of the BBC series *Sherlock* [93], adopting the preprocessing framework from Ke *et al*. [59]. Group-level valence and arousal trajectories were generated by averaging continuous ratings from 60 independent behavioral participants (30 per dimension), provided at the fMRI temporal resolution. Procedures were approved by the University of Chicago Institutional Review Board.

#### Dataset 4 (Merlin)

We considered an open fMRI dataset of 18 native English speakers viewing a 25-minute excerpt of the BBC series *Merlin* [94]. Continuous valence and arousal ratings were obtained from behavioral data released by Ke *et al*. [59] and aligned to the fMRI data using the same pipeline as the *Sherlock* dataset to serve as external validation targets.

### fMRI preprocessing and parcellation

We utilized publicly available preprocessed fMRI data for all datasets, aligning any additional harmonization steps with the Emo-FilM pipeline. Dataset-specific details for the external datasets are documented in their respective original publications [93, 94, 107].

For Emo-FilM, imaging data were acquired on a 3T Siemens Magnetom TIM Trio scanner with a 32-channel head coil. T1-weighted anatomical images were captured using an MPRAGE sequence (TR = 1.9 s, TE = 2.27 ms, flip angle = 9°, voxel size 1 × 1 × 1 mm). Functional images were acquired via a multiband gradient-echo EPI sequence (TR = 1.3 s, TE = 30 ms, flip angle = 64°, multiband factor = 3, voxel size = 2.5 × 2.5 × 2.5 mm). Audiovisual stimuli (mean duration = 11.4 min; total 2h 40 min) were flanked by 90-s pre-and post-stimulus fixation periods.

The Emo-FilM functional data were preprocessed using FSL FEAT (v.6.00), which included motion correction, brain extraction, linear standard-space co-registration, spatial smoothing (6-mm FWHM Gaussian kernel), grand-mean intensity normalization by a single multiplicative factor, high-pass temporal filtering (Gaussian-weighted least-squares straight line fitting, with sigma=50s), and nuisance regression of white matter, cerebrospinal fluid, and six rigid-body motion parameters [91].

For *JoJo Rabbit* database, data were acquired on a 3T Philips Ingenia (32-channel head coil). Functional images: GRE-EPI pulse sequence with TR = 2000 ms, TE = 30 ms, flip angle = 75°, 39 slices, voxel size = 3 × 3 × 3 mm. Anatomical images: T1-weighted MPRAGE with voxel size = 1 × 1 × 1 mm. See [107] for more details.

For *Sherlock* database, data were acquired on a 3T Siemens Skyra (20-channel head coil). Functional images: EPI pulse sequence with TR = 1500 ms, TE = 28 ms, flip angle = 64°, 27 slices, voxel size = 3 × 3 × 4 mm. Anatomical images: T1-weighted MPRAGE with 0.89 mm^3^ voxel size. See [93] for more details.

For *Merlin* database, data were acquired on a 3T Siemens Skyra (20-channel head coil). Functional images: EPI pulse sequence with TR = 1500 ms, TE = 28 ms, flip angle = 64°, 27 slices, voxel size = 3 *×* 3 *×* 4 mm. Anatomical images: T1-weighted MPRAGE with 0.89 mm^3^ voxel size. See [94] for more details.

For all functional image preprocessing, we adopted the same approach as in the Emo-Film preprocessing pipeline (see above and [91]). The only exception was in the *Merlin* database, for which only the already brain-extracted image was available. This differs from the standard FSL registration pipeline, which uses both the head and brain-extracted images.

For each participant and run, regional BOLD time series were extracted by averaging voxel signals within *N* = 119 brain regions. This atlas comprised 100 cortical regions from the Schaefer parcellation [96], alongside 16 subcortical, 2 cerebellar regions, and the brainstem from the Human Connectome Project release (Atlas ROI2.nii.gz).

To isolate stimulus-evoked dynamics, time series were cropped to match narrative presentation windows, accounting for baseline periods and hemodynamic lag. For Emo-FilM, we discarded the initial 72 baseline volumes and an additional four volumes (5.2 s) to account for hemodynamic delay; the final analyzed segments included four volumes captured post-offset. For the *Jojo Rabbit, Sherlock*, and *Merlin* datasets, time series were cropped as in the original data releases, except that the audiovisual cartoon opening each *Sherlock* run was removed (see *External dataset annotations*). Finally, all regional time series were z-scored within each participant and run.

### Emotion annotations and temporal alignment

We related time-resolved neural representations to continuous affective annotations, resampling all emotion time series to match the temporal resolution and length of the processed fMRI data.

#### Emo-FilM annotations

For Emo-FilM, we recon-structed consensus time series from raw behavioral annotations sampled at 1 Hz. The feature set comprised 50 continuous items: 13 discrete emotion labels and 37 componential measures spanning appraisal, expression, motivation, physiology, and subjective feeling [91]. Following published preprocessing protocols [91, 92], ratings were z-scored across films within each rater, missing samples were linearly interpolated, and constant or extreme series ( |*z*| *>* 15) were removed. As done in Ref. [92], to maximize consensus, we iteratively excluded individual raters if their removal increased the mean pairwise Pearson correlation by *>* 0.20, maintaining 3–4 top-agreeing raters per item. The remaining ratings were averaged into a single consensus time series. Notice that the consensus annotations were independently validated in the fMRI cohort using post-scan ratings of excerpts from the same films. Agreement between these validation ratings and the consensus annotations averaged *r* = 0.41 across films and items, comparable to the inter-rater agreement in the original annotation sample (*r* = 0.39), supporting their robustness across independent participant samples [91].

#### Valence, arousal, and power dimensions

To extract broad affective dimensions from Emo-FilM, we concatenated and standardized the 50-feature matrices across all films, then applied factor analysis with an oblique Promax rotation, following recent work [92]. Regression-derived factor scores for the three retained components — labeled valence, arousal, and power based on loading profiles — were partitioned back into film-specific time series for targeted low-dimensional analyses. All primary analyses used the full 50-feature representation.

#### External dataset annotations

For cross-dataset validation, we relied only on existing or derived continuous valence and arousal trajectories from independent behavioral cohorts. For *Jojo Rabbit* [107], valence and arousal were derived from continuous ratings of 20 emotion categories provided by 47 viewers. We fitted a three-factor Promax model to these ratings (similarly to Morgenroth *et al*. [92]), identified the activation factor as arousal, and defined valence as the standardized pleasant-factor score minus the unpleasant-factor score. For *Sherlock* and *Merlin*, we used the published group-average valence and arousal ratings [59], sampled at one value per TR. In the scanner, each of the two *Sherlock* runs began with the same 30s audiovisual cartoon (*Let’s All Go to the Lobby* ) [93], which was not part of the rated video. We therefore discarded the first 26 volumes (39 s) of each run, so that the remaining 920 + 1,004 = 1,924 volumes corresponded one-to-one to the ratings. When necessary, we linearly downsampled external annotation time series to match the specific TR and volume count of their corresponding fMRI runs (e.g. in JojoRabbit). As in Emo-FilM, BOLD data were shifted forward relative to film onset to account for hemodynamic lag by 3 TRs, 3 TRs, and 2 TRs for Sherlock, Merlin, and Jojo Rabbit (i.e., corresponding to 4.5 s, 4.5 s, and 4.0 s, respectively).

### Time-resolved BOLD, edge and higher-order representations

To capture the dynamic and distributed organization of affective processing, we characterized brain activity at four complementary levels: regional BOLD activity (*BOLD* ), instantaneous pairwise co-fluctuations (*edges*), higher-order coherent triplets (*triangles*), and the topological organization of these interactions across the network (*scaffold* ). The latter two representations were derived using a recently developed topological framework for reconstructing instantaneous higher-order group structure from multivariate time series [1, 75].

Let *x*_*i*_(*t*) denote the BOLD signal of region *i* ∈ 1, …, *N* at time point *t* ∈ 1, …, *T*, where *N* = 119. As described above, signals were standardized separately within each participant and stimulus run:

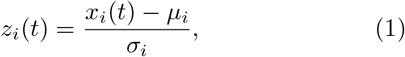

where *µ*_*i*_ and *σ*_*i*_ are the temporal mean and standard deviation of the signal from region *i*. The BOLD representation at each time point was defined as the vector of standardized regional activity,

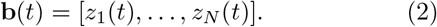

#### Edge co-fluctuations

While traditional functional connectivity evaluates time-averaged statistical dependencies, capturing the rapid shifts inherent to emotional experiences requires a temporally resolved approach. To achieve this, we computed edge time series to track instantaneous pairwise coordination [60, 61]. For each pair of regions (*i, j*), we computed an edge time series,

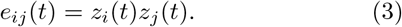

Positive and negative values indicate co-occurring deviations in the same or opposite directions, respectively. Because the temporal average of *e*_*ij*_(*t*) exactly recovers the Pearson correlation between regions *i* and *j*, this method provides a mathematically exact, moment-to-moment unwrapping of traditional functional connectivity without requiring sliding windows. Furthermore, these co-fluctuations can be interpreted as interaction terms, capturing shared variance beyond the isolated activity of either region [137]. At each time point, the full edge representation comprises the upper triangular elements of the *N × N* co-fluctuation matrix, yielding 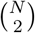 unique features.

#### Higher-order co-fluctuations and the triangle representation

To investigate functional architectures extending beyond dyads, we tracked time-resolved group interactions using a recent topological representation [1, 75]. For each region triplet (*i, j, k*), the instantaneous third-order co-fluctuation was computed as the element-wise product of their standardized BOLD signals:

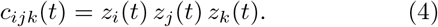

To allow for direct topological comparisons with pairwise co-fluctuations, this product time series was standardized across time:

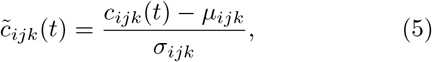

where *µ*_*ijk*_ and *σ*_*ijk*_ are the temporal mean and standard deviation of *c*_*ijk*_(*t*). We then remapped its sign so that positive values denote fully concordant triplets, in which all three signals were simultaneously above or below their means, whereas negative values denote discordant configurations, and the magnitude of the higher-order co-fluctuation is then defined as:

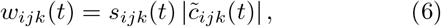

where *s*_*ijk*_(*t*) = +1 when sgn[*z*_*i*_(*t*)] = sgn[*z*_*j*_(*t*)] = sgn[*z*_*k*_(*t*)], and *s*_*ijk*_(*t*) = *−*1 otherwise. This transformation maps fully concordant three-region co-fluctuations to positive weights, effectively isolating the coherent contributions within the multivariate time series. At each time point *t*, the pairwise edges (*w*_*ij*_) and triplet triangles (*w*_*ijk*_) define a weighted simplicial complex [1, 64]. We evaluated its multiscale topology using persistent homology [138–140] by constructing a descending filtration, *S*_0_ ⊂ *S*_1_⊂*…* ⊂ *S*_*l*_, where links and triangles were progressively added by thresholding weights in descending order (see Ref. [1] for further details).

During filtration, we isolated higher-order interactions by identifying triplets that violated the simplicial closure condition. A triplet (*i, j, k*) was classified as a violating triangle if its higher-order weight strictly exceeded its weakest bounding edge:

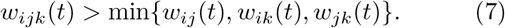

Because these violating triangles (Δ_*v*_) cannot be reduced to their constituent pairwise links at that filtration level, they serve as a localized topological signature of non-simplicial higher-order organization. The final *triangle* representation at each time point consists of the weights of these violating triangles across all 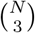 possible triplets, with non-violating triplets set to zero. To evaluate these interactions at the edge level, we applied a downward projection, assigning each edge (*i, j*) the average weight of all violating triangles containing it. That is, for each time point *t*, we have 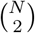 unique features.

#### Homological scaffold

To characterize how individual connections contribute to the mesoscale topology tracked along this same filtration sequence, we constructed a time-resolved frequency homological scaffold [1, 75]. Along the descending filtration, persistent homology tracks the evolution of one-dimensional topological features (*H*_1_) — closed edge paths that emerge as thresholds are lowered and are subsequently filled by triangles. While the persistence of these features captures mesoscale organization distinct from local edge strength or clique structures, persistent homology is coordinate-free and does not directly localize these loops to specific network components.

The homological scaffold [73, 141] resolves this by projecting the representative *H*_1_ generators identified during the filtration back onto their constituent edges, providing an edge-level localization of the network’s mesoscale topological organization. Specifically, at each time point *t*, each edge *e* is assigned a frequency scaffold weight equal to its total membership across the set of active cycle generators *G*(*t*):

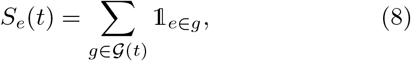

where 1_*e*∈*g*_ is the indicator function. Thus, if an edge belongs to *s* distinct generators, *g*_1_, …, *g*_*s*_, then *S*_*e*_(*t*) = *s*. Crucially, as a frequency-based mapping, *S*_*e*_(*t*) counts the structural recurrence of an edge across distinct generators rather than scaling by individual cycle longevity or persistence.

Because the resulting matrix *S*(*t*) ∈ ℝ^*N×N*^ is symmetric, extracting its upper triangular elements yields exactly 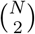 unique features per time point. This explicitly matches the dimensionality of our edge and downward-projected triangle representations, allowing direct comparative analyses across different hierarchical tiers of network organization. High scaffold weights highlight connections heavily embedded within the broader edge–triangle architecture, which links distinct coherent configurations, independent of their raw pairwise co-fluctuation magnitude. This framework extends the static homological scaffold [73] to frame-by-frame co-fluctuation patterns, tracking how topological embedding reconfigures over time, in line with evidence that scaffold architectures capture reproducible, individual-specific aspects of functional brain organization [79].

### Time-time correlation matrices and emotion-brain clustering analysis

To characterize the temporal organization of neural and emotional dynamics during movie viewing, we constructed time–time correlation matrices for each film. This framework, which compares multivariate configurations across all pairs of time points, has been previously validated as a robust method to assess how regional, edge-based, and higher-order fMRI representations recover task and resting-state temporal structure [75, 95]. Here, we adapted this approach to test whether recurring brain configurations align with recurring multivariate emotional states.

For the emotion data, each time point was represented by its 50-dimensional emotion profile, and the emotion time–time matrix indexed the Pearson correlation between profiles across all time points. Analogous matrices were constructed independently for regional BOLD activity, edge time series, projected triangle weights, and homological scaffold weights. Thus, each entry quantifies the similarity between the complete neural configuration at two time points.

To ensure robust statistical inference, we implemented a paired subsampling protocol (1000 iterations). In each iteration, 25 of the 30 participants were sampled without replacement, and their participant-level brain time–time matrices were averaged. To preserve the validity of paired comparisons, the same participant subset was used across all neural representations. Similarly to Santoro *et al*. [75], each time–time matrix was binarized into a temporal graph via percentile thresholding: we retained the top 5% of similarities for edges, triangles, and scaffold, and the extreme 5% (upper and lower 2.5% tails) for regional BOLD to capture both high similarity and high anti-similarity configurations.

Temporal states — representing stable narrative scenes — were identified using a consensus Louvain clustering pipeline [142, 143]. Given the stochastic nature of modularity optimization, we generated 50 initial Louvain partitions, constructed a nodal co-assignment matrix, thresholded this matrix at the 95th percentile, and obtained the consensus clustering partition. This consensus approach has been previously validated as a reliable strategy for segmenting complex temporal dynamics in naturalistic paradigms [143].

We quantified the correspondence between each brain-derived temporal partition and the emotion-derived partition using element-centric similarity (ECS) [98]. ECS compares the community relationships surrounding individual elements and is therefore insensitive to differences in the number or size of communities across partitions. Here, the elements were TRs; ECS values range from 0 for dissimilar partitions to 1 for identical partitions. Higher ECS values indicate that time points grouped by a neural representation also share similar multivariate emotional profiles.

Because participant-level scaffold estimates are unreliable and ECS is defined from the group-average community structure, sensitivity to participant composition was assessed by repeated subsampling. These subsamples quantify the stability of the group-level result rather than a sampling distribution and were therefore summarized descriptively rather than converted to *p* values. The primary claim — that the scaffold was the top-ranked representation across films — was tested with a one-sided exact binomial test on the number of films (out of 14) in which it achieved the highest mean ECS, under a null probability of 0.5. We report the exact *p* value, the corresponding proportion of films, and its one-sides 95% Clopper–Pearson confidence interval. For each film, subsampling stability was defined as the proportion of 1,000 subsamples in which scaffold ECS exceeded all three alternative representations; no multiple-comparison correction was applied to these descriptive estimates. As the same 30 participants contributed to all films, film-level observations are not fully independent, making the subsampling analysis a complementary robustness assessment at the participant-composition level.

### Emotion prediction and statistical inference

We predicted continuous affective time courses from four neural representations — regional BOLD activity, edge time series, projected triangles, and the homological scaffold — using kernel ridge regression (KRR) [101]. To isolate behaviorally relevant dynamics and place predictors and targets on a common timescale, we convolved all time series with a Gaussian kernel (*σ* = 15 TR ≈ 19.5 s; TR = 1.3 s) independently within each film to mitigate noise. Sensitivity analyses confirmed that the representational ordering was preserved for smoothing widths of 10 TRs and above (Supplementary Fig. S9)

We solved the KRR model in its dual form with a linear kernel:

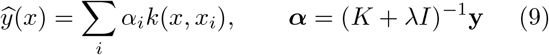

For the three-dimensional targets, a single ridge penalty was selected for each brain representation using arousal across the full set of 14 films and then held fixed across all folds and all three dimensions. This tuning procedure therefore differs from the strictly nested approach used for the item-level analysis in Fig. 4**a–b**. Because the same tuning strategy was applied consistently across representations, panels **c–e** are used to compare their relative decoding performance rather than to provide fully nested estimates of out-of-sample accuracy. This dual formulation accommodated both low- and high-dimensional representations, while the ridge penalty (*λ*) stabilized estimation against collinear features and temporal autocorrelation. Generalization was evaluated via leave-one-film-out cross-validation across the 14 short films (Fig. 4). Within each fold, predictors were mean-centred using the 13 training films, while targets were retained on their original scale. For the 50-item analysis, the regularization parameter (*λ* ∈ 10^*−*3^, …, 10^4^) was selected by nested leave-one-film-out cross-validation within the training set, maximizing mean validation accuracy. Prediction accuracy was quantified as the Pearson correlation (*r*) between the predicted and observed smoothed trajectories in the held-out film.

Statistical inference used the held-out film as the unit of replication. For the dimensional targets (valence, arousal, power), representations were compared with the Nadeau–Bengio corrected resampled *t*-test [103], which accounts for training-set overlap between leave-one-film-out folds.

The 50 emotion regressors required a different approach, because they are strongly intercorrelated (mean absolute correlation of 0.55 per film at the *σ* = 15 TR smoothing used for prediction), violating the independence assumed by most paired tests. As a result, we compared representations film by film. Each of the six possible comparisons contrasts two of the four neural representations, e.g., the scaffold against BOLD activity. For a given comparison and a given held-out film, we consider the difference in prediction accuracy between the two representations, Δ*r* = *r*_*scaffold*_ *−r*_*BOLD*_, separately for each of the 50 emotions, and averaged those 50 values into one number: the mean accuracy advantage of one representation over the other in that film. Repeating this for all films yields 14 values, one per held-out film, and these were the values tested. Significance was assessed with an exact sign-flip test [144]. Under the null hypothesis that the two representations predict equally well, each film’s advantage would be as likely to come out positive as negative, so reversing its sign would leave the data equally probable. We therefore recomputed the mean across films under all 2^14^ combinations of signs; the *p*-value is the proportion of those means whose absolute value equaled or exceeded the observed one. Flipping whole films rather than individual emotions holds the 50 correlated emotions together, so their covariance never enters the test. The Nadeau–Bengio corrected *t*-test on the same 14 values is reported alongside as a sensitivity analysis for the training data shared between folds; with only 14 folds, its variance correction is approximate and conservative. The two tests address different dependencies — the sign-flip test the correlation among the 50 emotions, the corrected *t*-test the overlap between training sets. All *p*-values were corrected using the Benjamini–Hochberg false-discovery-rate (FDR) procedure, with effect sizes reported as paired Cohen’s *d*.

### Dimensionality of affective and neural spaces

To assess intrinsic dimensionality, we applied principal-component analysis (PCA). For the emotion space, we decomposed the 50 unsmoothed regressors, separately within each film, and reported the variance captured by each component and the cross-film variability of the cumulative spectrum (Fig. 5**a**). For the brain representations, we performed PCA separately within each film on the smoothed feature time series, estimating the effective dimensionality (*n*_80_) as the average number of components required to explain 80% of the variance (Fig. 5**b**).

### Spatial organization and shared neural substrates

Because the raw backward weights (**w** = *X*^*⊤*^***α***) of regularized decoders are uninterpretable under severe feature collinearity, as shown by Haufe *et al*. [106], we mapped spatial architecture using independent, weight-free encoding maps. For each neural feature (i.e., ROIs or edges), we computed its Pearson correlation with the target time course across the training films. To ensure sparsity and cross-fold stability, we retained the strongest 10% of associations within each fold and intersected these selections across all 14 cross-validation folds. This non-circular approach parallels connectome-based predictive modeling [59, 104, 145] but employs a fixed selection fraction to maintain stable sparsity across folds.

To quantify the shared neural substrate between distinct emotions, we computed the cosine similarity between their binary selection maps (Fig. 5**c**) and the Jaccard overlap of their scaffold edge sets:

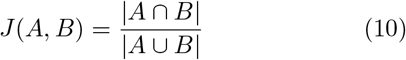

yielding the emotion-by-emotion overlap matrix shown in Fig. 5**d**. To prevent inflated overlap estimates driven by temporal autocorrelation and cross-emotion correlations, we expressed these metrics as an excess over a phase-randomized surrogate null. Using the Prichard and Theiler shared-phase method [146], this null preserved each channel’s power spectrum and the cross-emotion covariance while scrambling the temporal alignment between brain and emotion data, isolating spatial overlap from nuisance correlations.

### Arousal network enrichment and decoder correspondence

To localize arousal-related information, we ranked features within each training fold by their signed correlation with arousal and retained the upper and lower 10% tails separately, intersecting each across the 14 folds; separating the tails preserved the direction of the association at a fixed selection density. For regional BOLD, where the feature space is two orders of magnitude smaller, a fixed decile leaves too few regions to survive the intersection across folds; we therefore retained regions whose per-fold correlation exceeded a one-sided *p <* 0.01 threshold and intersected these across the 14 folds, and rendered the resulting fold-stable per-ROI correlations directly on cortical and subcortical surfaces. For the connectivity representations, selected edges were summarized within an 8 *×* 8 matrix spanning seven canonical resting-state networks and one subcortical system, with positive and negative associations displayed in the upper and lower triangles (Fig. 5**e**). Because network pairs vary in their total possible edges (*E*_*ab*_), we corrected for anatomical size using a density-excess metric:

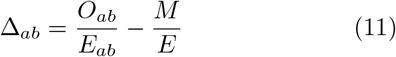

which subtracts the uniform chance baseline (*M/E*) from the observed proportion of selected edges (*O*_*ab*_*/E*_*ab*_), where *M* is the total number of selected edges and *E* the total number of possible edges. Enrichment was assessed with a one-sided hypergeometric test, with Benjamini– Hochberg correction applied across the 36 unique network pairs separately within each representation and each tail; cells surviving *p*_*FDR*_ *<* 0.05 and *p*_*FDR*_ *<* 0.01 are marked with one and two asterisks, respectively.

Finally, to verify that the correlation-based encoding maps captured the signal exploited by the KRR decoder, we compared them against the decoder’s forward activation pattern [106]:

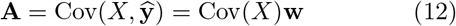

and against its raw backward weights (**w**). Correspondence was quantified within each fold by cosine similarity, Pearson correlation, and Jaccard overlap of the top-decile features, confirming the validity of the encoding-map approach (Supplementary Fig. S12).

### Partial least squares correlation (PLSC)

We applied partial least squares correlation (PLSC) [99] to identify latent patterns of covariation between the emotional content of each scene and its associated brain representation. Before the PLSC analysis, the 50-item emotion rating time series were smoothed using a Gaussian filter (*σ* = 15 TRs). Both the emotion and brain features were then averaged within emotion-derived temporal scenes. For each film, scenes were obtained by consensus Louvain clustering of the time–time correlation matrix of the unsmoothed 50-item ratings (95th-percentile threshold, 10 partitions), excluding communities shorter than 3 TRs. Scene-level emotion values used the *σ* = 15 TR smoothed ratings, whereas brain features used unsmoothed group averages. The resulting scene-level matrices were concatenated across 14 films, yielding 190 total scenes, and were subsequently z-scored column-wise.

We performed PLSC separately for each brain representation to relate the scene-averaged, z-scored emotion profiles (*X* ∈ ℝ^190*×*50^) to their corresponding neural features 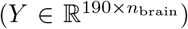. The brain feature space *n*_brain_ comprised either the mean BOLD activity across *N* regions, or the upper-triangular elements of the *N ×N* matrix for the three edge-based representations (ETS, projected triangles, and the homological scaffold), yielding 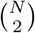 features per scene.

PLSC identifies orthogonal latent variables (LVs) that maximize the covariance between linear combinations of emotion features (behavioral) and linear combinations of brain features across scenes. Specifically, singular value decomposition (SVD) of the cross-covariance matrix between *X* and *Y* yielded these LVs, with corresponding emotion and brain saliences given by the right and left singular vectors. We quantified the proportion of shared emotion–brain covariance captured by each LV as its squared singular value divided by the sum of squared singular values across all components.

Each LV was evaluated against its permutation distribution (5000 permutations, scene labels shuffled). LVs with *p <* 0.05 were retained; these *p* values were not adjusted across LVs or representations. A film-restricted permutation (Supplementary Note B) provides a more conservative test.

To evaluate the reliability of individual brain and emotion weights, we utilized bootstrap resampling of scenes with replacement (5000 iterations). Stability was quantified using bootstrap ratios (BSRs). Emotion-side BSRs were computed from PLSC saliences, whereas brain-side BSRs were computed from feature loadings, defined as correlations with the emotion latent scores, across 5000 bootstrap samples. Features were considered robust if they met a defined BSR magnitude threshold (e.g., |*BSR*| ≥ 2). This is analogous to considering salience weights as significant when their 95% bootstrap confidence intervals did not include zero.

For a compressed functional network representation, reliable edges were mapped to an 8 *×* 8 matrix representing the seven resting-state networks [147] plus a subcortical network. To determine whether an LV contained more reliable edges than expected by chance within a specific network pair, we computed a one-sided hypergeometric p-value for each cell, followed by an FDR correction across the 72 unique network-pair-by-sign tests within each LV.

All PLSC analyses were implemented using the behavioral pls function from the pyls Python package (https://github.com/netneurolab/pypyls).

### Cross-dataset validation

#### Zero-shot prediction and model transfer

To evaluate whether the affective neural architectures identified in Emo-FilM generalize to independent narratives, models were evaluated without refitting on three external datasets: Sherlock, Merlin, and Jojo Rabbit. Affect targets were constructed and standardized independently for each dataset, with Jojo Rabbit targets derived from its native set of 20 emotion indicators. To align timescales across varying acquisition parameters, neural and affective features were smoothed using Gaussian kernels matched in physical time ( ≈ 20 s standard deviation), strictly preventing smoothing across acquisition boundaries. Within Emo-FilM, the scale-free ridge parameter (*α* = *λ*tr(*K*)*/n*), with *λ* selected from a common logarithmic grid spanning 10^*−*3^ to 10^4^, and all centering/scaling parameters were selected exclusively using training-fold statistics before the final model was applied to the external data. External features were standardized using the mean and variance of each complete external dataset, without using affect labels, before applying the trained model. Zero-shot performance was quantified as Pearson’s correlation between predicted and observed trajectories, averaged equally across the three external datasets. Significance was assessed by phase-randomizing the target series separately within each film, run, or movie part. Only the observed time points were randomized; temporary samples added during smoothing were excluded, preventing boundary artefacts and mixing between separate acquisitions. Probabilities were then combined across the external datasets and FDR-corrected across the tested representations and affective dimensions. The corrected arousal results were BOLD *p*_*FDR*_ = 0.032, Edges *p*_*FDR*_ = 0.00003, Triangles *p*_*FDR*_ = 0.00009, and Scaffold *p*_*FDR*_ = 0.016.

In a complementary leave-one-dataset-out analysis, each dataset was held out in turn, and models were trained on the concatenated, within-dataset z-scored time series from the other three, with contribution proportional to dataset duration (Emo-FilM: 60–72% of training samples). Because valence and arousal were correlated in three datasets, we additionally computed partial correlations after regressing the other affective dimension from both prediction and target. Significance was assessed using 1000 phase-randomized surrogates of the residualized target and FDR-corrected across representations and dimensions.

#### Ranked-edge perturbation

To test the anatomical redundancy of the arousal-predictive network, the 7021 edge features in Emo-FilM were ranked by the absolute magnitude of their training-set association with arousal. For predefined thresholds *k*, we compared a model retaining only the top *k* edges against one refitted on the remaining complement. These perturbed models were evaluated zero-shot on the external datasets, with uncertainty estimated via boundary-respecting paired temporal resampling. Practical equivalence to the full-edge model was defined *a priori* as a 95% confidence interval falling entirely within ±0.02 correlation.

#### Cross-dataset network organization

To determine whether arousal recruited the same largescale network interactions across datasets, we summarized each edge–arousal map over the 36 possible pairs of functional systems. For each dataset, we correlated every edge with arousal, standardized these correlations across the 7021 edges, and averaged them within each of the 36 network pairs, yielding one signed profile per dataset; profiles were then correlated across the six dataset pairs. For display only, Supplementary Fig. S15**b** shows positive and negative associations separately. Significance was assessed using two complementary controls. First, we disrupted the timing of the arousal trajectory separately within each film, run, or movie part while preserving its slow temporal structure. Second, we reassigned cortical regions among functional networks while preserving network sizes and the non-cortical system. These controls tested whether correspondence depended both on the temporal relationship with arousal and on the organization of the cortex into canonical functional networks.

#### Cross-dataset geometry of affective states

Because the four narratives differed in content, duration, and timing, their neural responses could not be aligned. We therefore used representational similarity analysis (RSA) [108, 148], which compares the relationships among neural states rather than their position in a shared timeline. This approach has been used in nauralistic neuroimaging to identify common event representations across individuals and across perception and memory [93, 94].

Valence and arousal were standardized within each dataset; the four standardized clouds were then pooled and partitioned once by *k*-means into *K* = 16 common affective states, and every time point of every dataset was assigned to its nearest state centroid. For each dataset and neural representation, we averaged the neural pattern associated with every state and measured the dissimilarity between all pairs of states, producing a representational dissimilarity matrix (RDM). We then correlated these matrices across datasets. High correspondence indicated that the relative arrangement of affective states was preserved across narratives, even when their neural time series and moment-to-moment affective trajectories could not be aligned directly.

Because each dataset contributed to three of the six dataset pairs, inference was based on their joint correspondence rather than treating the six correlations as independent observations. Uncertainty was estimated by repeatedly resampling films or runs and contiguous temporal segments within them, using identical resamples for every neural representation. Finally, to ensure that the result did not depend on the choice of (K=16), we repeated the complete RSA across state-space resolutions (see supplementary Fig. S13). The principal pattern of cross-dataset correspondence remained stable across this range.

#### Arousal adjustment and sensitivity analyses

To isolate cross-dataset geometric correspondence from broad arousal effects, neural RDMs were adjusted for measured differences in mean arousal. For each pair of datasets, the adjustment covariate was defined as the average of their respective state-pair absolute arousal-difference matrices. The neural RDM vectors and this covariate were rank-transformed; the covariate was then regressed from each neural RDM, and the resulting residual vectors were correlated via a partial-Spearman procedure. Notice that while this isolates correspondence from measured mean arousal, it does not remove all arousal-related structure

## Code availability

All Python code required to reproduce the principal findings will be made publicly available in a GitHub repository upon publication. Subcortical visualizations were obtained using the Python version of the `subcortical_visualization package` [149].

## Data availability

The neuroimaging and emotion data analyzed in this study were obtained from public repositories, namely, Emo-FilM fMRI data: ds004892; annotations: ds004872; Jojo Rabbit fMRI data and annotations: 28255745; Sher-lock fMRI data: [Princeton data common discovery](https://datacommons.princeton.edu/discovery/catalog/doi-10-34770-9ndy-8c50); Merlin fMRI data: ds001110. Emotion annotations for Sherlock and Merlin are available here: [emotion annotations](https://github.com/jinke828/AffectPrediction/tree/main/beh_ratings). The preprocessed fMRI data will be available on Zenodo upon publication.

## Acknowledgment

This research was funded by the Swiss National Science Foundation (Sinergia grant CRSII5 180319 to P.V. and D.V.D.V.) and benefited from resources of the Swiss Center of Affective Sciences at UNIGE. Data collection was conducted on the imaging platform at the Brain and Behavior Lab (BBL) and benefited from support of the BBL technical staff and the Center for Biomedical Imaging (CIBM). A.S. has received funding from the European Union’s Horizon 2020 research and innovation programme under the Marie Sklodowska-Curie grant agreement no. 101208090. A.S. thanks D. Orsenigo and M. Nurisso for helpful discussions. We thank S. Romanato for very preliminary analyses on audio-video signals.

AI tools — OpenAI ChatGPT (GPT-5.6), Anthropic Claude (Claude Opus 5), and Google Gemini (Gemini Pro) — were used to assist with code development and debugging the authors’ code, preliminary figure templates, and language editing. All analyses, outputs, and scientific decisions were produced or verified by the authors, who take full responsibility for the manuscript. AI was not used to generate or modify research data.

## Supplementary Information

**Figure S1.**
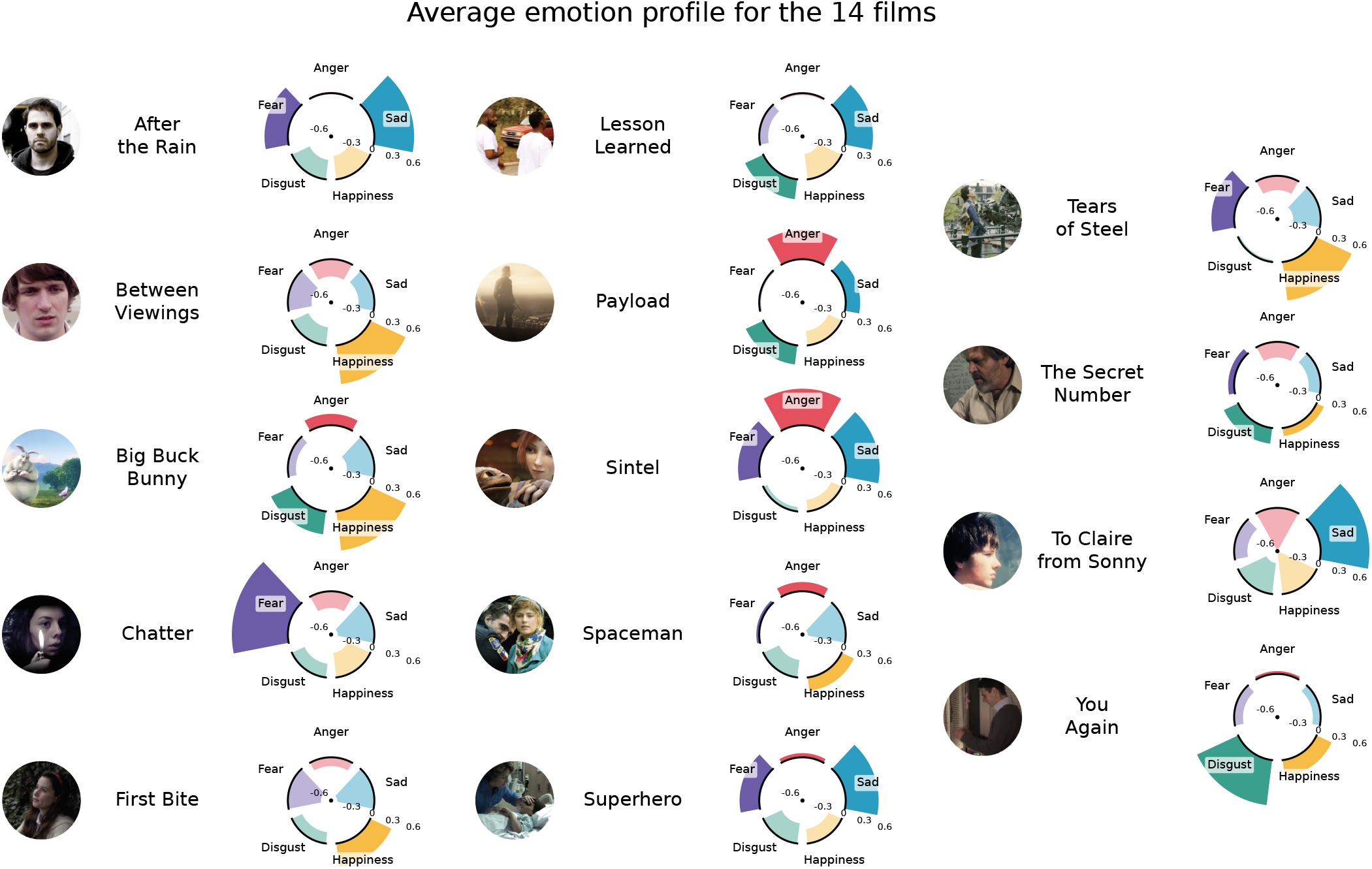
Film-level emotional profiles across the 14 Emo-FilM films. Radial plots summarize five illustrative discrete-emotion annotations — anger, sadness, happiness, disgust, and fear — selected from the 13 discrete labels among the 50 continuous affective features. For each film and emotion, the radial value represents the temporal mean of the corresponding consensus time course, derived from three or four independent annotations. Ratings were standardized across films within each annotator before consensus estimation; values above and below zero therefore indicate relatively higher and lower intensity for that emotion. The black arcs mark zero. These static profiles highlight differences in the overall emotional composition of the films while discarding their moment-to-moment dynamics.

**Figure S2.**
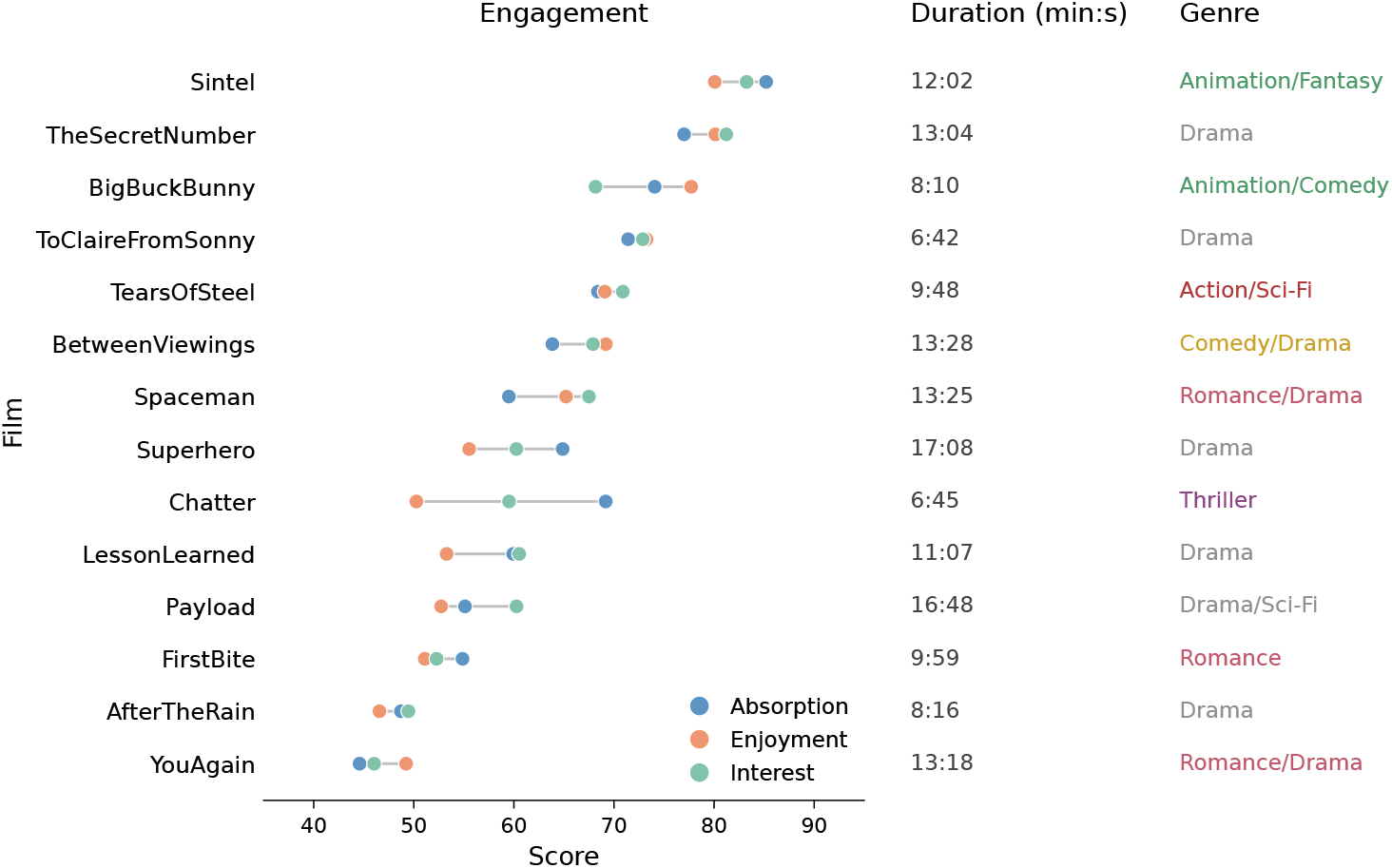
Engagement scores across films in the Emo-FilM dataset. Engagement profiles for the 14 films included in the Emo-FilM dataset, spanning a total of two hours and forty minutes of naturalistic stimuli. Each film is characterized by three distinct scores quantifying subjective engagement: absorption (blue), enjoyment (orange), and interest (green). These scores were rated on a continuous scale from 0 to 100, and colored points indicate the score for each engagement dimension, with gray line segments connecting the three scores within each film to illustrate their relative distribution. To facilitate comparison, the films are ordered according to their overall engagement-score profiles. Film duration (approximately 6-17 minutes) and genre are displayed alongside each film for additional context. The visualization highlights substantial heterogeneity in both the magnitude and the relative balance of absorption, enjoyment, and interest across the diverse film selection, reflecting the use of these naturalistic stimuli to evoke a rich and varied spectrum of emotional experiences.

**Figure S3.**
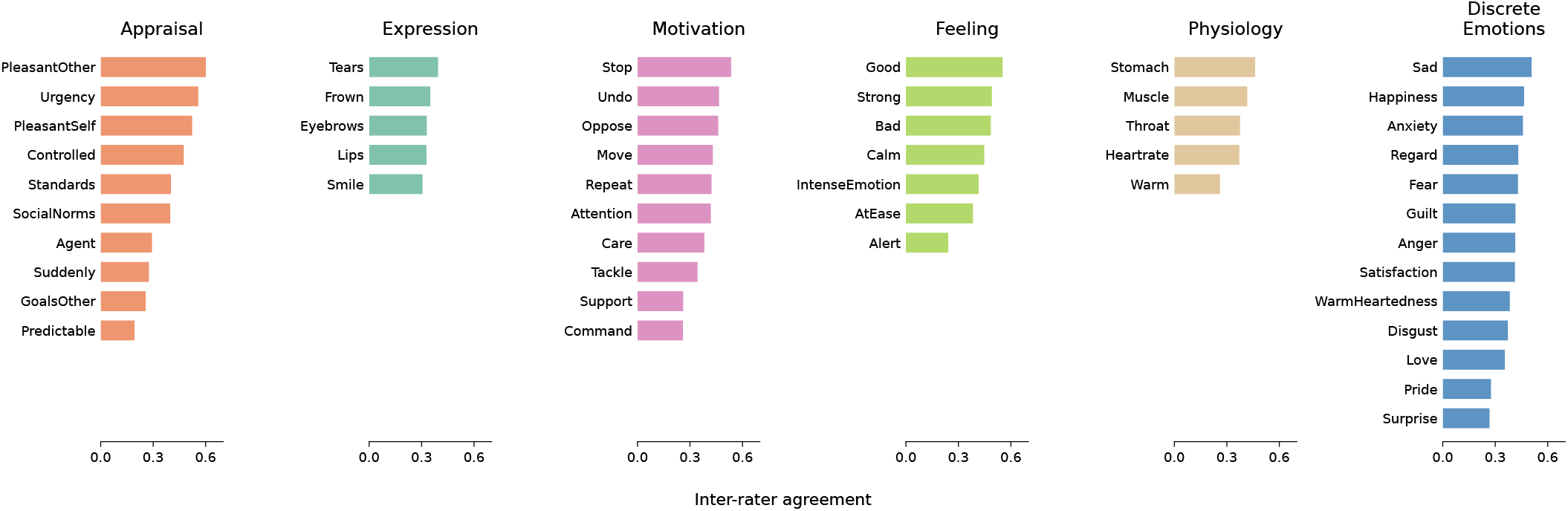
Emotion annotation items and inter-rater agreement. The 50 emotion regressors retained in the Emo-FilM dataset are grouped according to their respective emotion components and discrete emotion categories. Based on the Component Process Model (CPM), the 37 emotion components comprise *Appraisal, Expression, Motivation, Feeling*, and *Physiology*, complemented by 13 *Discrete Emotions*. Bars indicate the mean inter-rater agreement for each item, quantified as the Pearson correlation coefficient (*r*) between all pairs of the annotators retained for the consensus of that item, averaged over pairs and then over film. Annotations were collected from a total cohort of 44 independent raters, yielding an overall mean inter-rater agreement of 0.39 across the dataset. This final set of 50 items represents a robust consensus, obtained after strictly excluding items that demonstrated poor agreement across the evaluated films (see [150]). Together, these dense annotations provide a highly granular, multi-level assessment of dynamic emotional phenomena over time.

**Figure S4.**
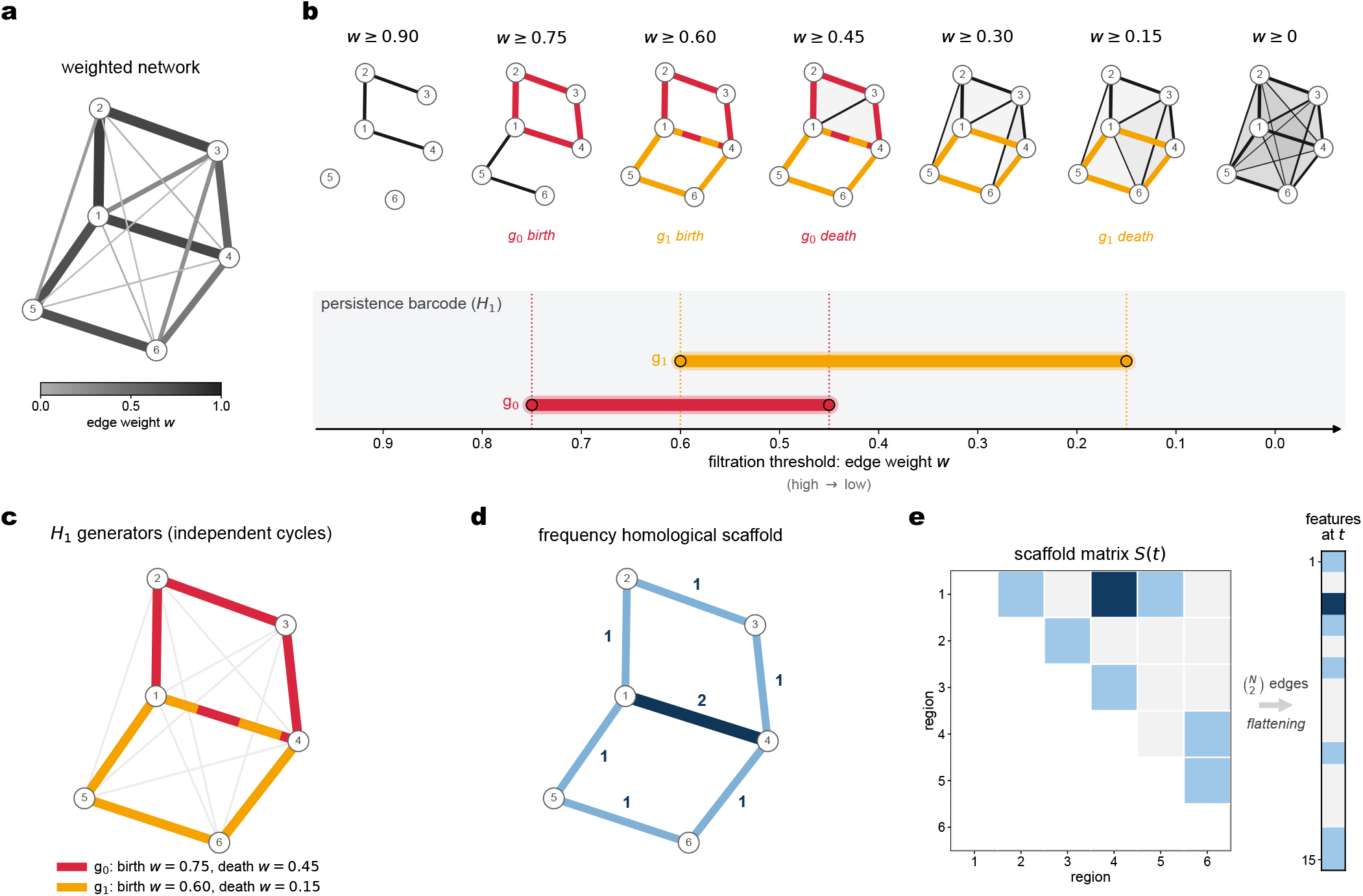
Schematic illustration of the homological scaffold construction and its mesoscopic view. **(a)** A sample weighted network comprising six nodes (1–6), with edge widths proportional to connection weights *w*. **(b)** Persistent homology scans through this network at every scale at once: as the threshold lowers (*w* = 0.90*→* 0), surviving edges are assembled into a simplicial complex where filled triangles (grey) mark groups of nodes that are already mutually connected. One-dimensional holes (*H*_1_)—closed paths enclosing unfilled regions—are born when their final edge enters the filtration and die when triangles tile them over. Two appear here: *g*_0_ (1–2–3–4), born at *w* = 0.75 and dying at *w* = 0.45, and *g*_1_ (1–4–6–5), born at *w* = 0.60 and dying at *w* = 0.15. The persistence barcode records these birth–death intervals, with long bars denoting cycles that survive across many scales and are therefore robust to the choice of threshold. **(c)** Representative generators returned by the filtration, drawn on the original layout. Each highlights a distinct mesoscale route that circumvents dense, fully connected cores. Generators represent homology classes and are not unique; those shown are the ones returned by the algorithm. Here, edge 1–4 belongs to both generators (*g*_0_ and *g*_1_) and is therefore drawn as a single line carrying both colours. **(d)** Superimposing the generators yields the frequency homological scaffold. Every edge is weighted by the number of distinct generators that traverse it, so the shared edge 1–4 carries *S*_*e*_ = 2 while the remaining scaffold edges carry *S*_*e*_ = 1; edges belonging to no generator drop out entirely. The scaffold is thus a sparse, cycle-based summary that scores a connection by its embedding in the mesoscale organization of the network rather than by its own weight. **(e)** The upper triangle of the scaffold matrix *S*(*t*) serves as the feature vector that enters all analyses reported in the main text: 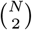 values per time point, which equates to 7,021 features for the *N* = 119 parcellation. For clarity, the construction shown here relies on a single static network, where triangles are filled as soon as their three edges are present. However, in the main text, the same procedure is applied instead to instantaneous edge and triangle co-fluctuation patterns at each time point. There, each simplex (i.e., edges and triangles) enters the filtration by its own weight, yielding one scaffold per TR [1].

**Figure S5.**
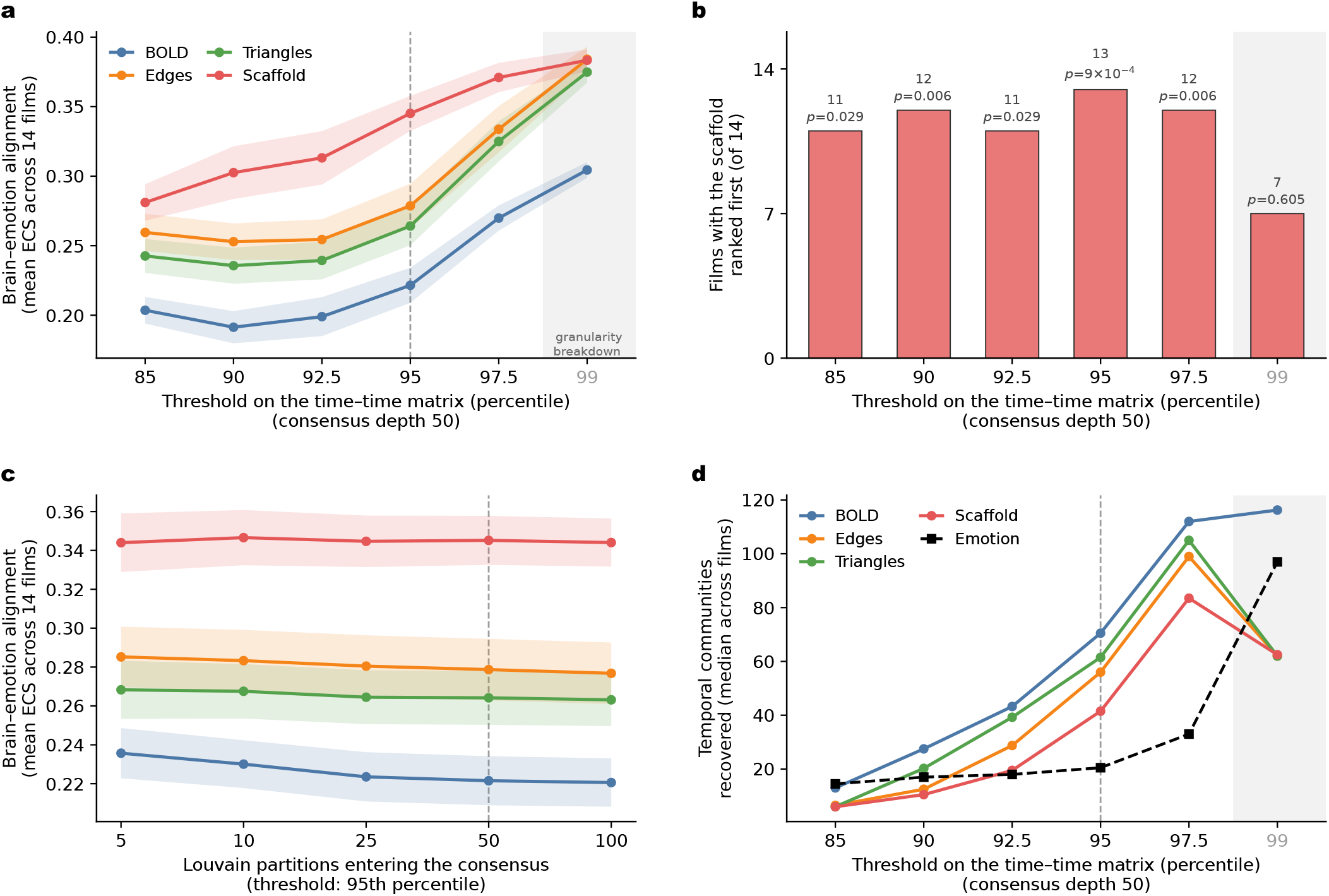
The homological scaffold alignment advantage is robust to clustering parameters. We varied the percentile threshold used to construct the time–time and consensus matrices and the number of Louvain partitions entering the consensus. **(a)** Mean element-centric similarity (ECS) across 14 films as a function of percentile threshold, with consensus clustering obtained from 50 iterations ( *±* standard error on the mean across films). The scaffold remains the best-performing representation from the 85th to 97.5th percentile. **(b)** Number of films in which the scaffold ranks first. It leads in 11–13/14 films across this range (one-sided exact binomial tests, all *p* ≤ 0.029), but not at the 99th percentile. **(c)** Mean ECS as a function of consensus depth at the 95th-percentile threshold. Results are essentially unchanged across depths, with scaffold ECS varying by only 0.003 and ranking first in 13–14/14 films. **(d)** Number of recovered temporal communities. Extreme thresholds produce mismatched partition granularity: low thresholds yield overly coarse brain partitions, whereas at the 99th percentile the emotion partition fragments while brain partitions coarsen, inflating ECS and collapsing the rank order. The 95th percentile provides comparable and interpretable temporal resolution across modalities and was the one originally proposed in Ref. [75]. Estimates are means over 100 subsamples (25/30 participants), with film as the unit of inference. Dashed lines mark the main-analysis parameters; the shaded region marks the degenerate 99th-percentile regime.

### A. Control for low-level audiovisual features

To verify that the homological scaffold’s temporal correspondence with emotion reflects true affective content rather than a generic sensitivity to recurring stimulus structure, we repeated the element-centric similarity (ECS) analysis using the films’ low-level audiovisual properties in place of the emotion annotations. Low-level luminance and loudness often covary with emotional content in film (for instance, fear-eliciting scenes may be intentionally darker or louder), making them critical nuisance regressors to consider [46].

#### Feature definitions and temporal alignment

We evaluated three specific low-level targets: an auditory set (5 features), a visual set (6 features), and their concatenation (11 features). Five auditory features were extracted from each film’s soundtrack using standard music-information-retrieval definitions [151, 152], following the descriptor set established for content-based audio classification [153]: the root-mean-square amplitude, zero-crossing rate, spectral flux, mean mel-frequency cepstral coefficient, and mean chroma.

Six visual features were extracted from the video stream at each film’s native frame rate (e.g., mean brightness, saturation, and hue on the left and right halves of each frame via the HSV representation). Brightness and saturation were explicitly included because they carry most of the affective variance attributable to color, with hue contributing considerably less [154]. To retain a coarse sensitivity to spatial layout, these visual features were computed separately over the left and right halves of each frame. Feature sets of this kind routinely underpin affective video content analysis [155, 156]. To align with the fMRI data, these features were smoothed via a boxcar moving average, sampled at equally spaced indices matching the cropped scan length, and *z*-scored across time. This ensured the low-level series shared the identical timeline, initial volume crop, and hemodynamic offset as the BOLD and emotion annotations. The targets were then processed through the exact same clustering pipeline as the emotion data (i.e., time-by-time Pearson correlation, 95th-percentile thresholding, and consensus Louvain clustering).

#### Dimensionality control

A potential confound in this comparison is that the emotion target contains 50 dimensions, whereas the low-level targets contain only 5 to 11. Time-by-time correlations computed over fewer columns are inherently noisier, which predictably fragments the Louvain partition (yielding a median of 47–58 communities across films for low-level features, compared to 20.5 for the emotion annotation). To ensure the emotion advantage reflects affective content rather than an artifact of target dimensionality, we rebuilt the emotion target from 20 random subsets of 11, 6, and 5 regressors and rescored the analysis. As expected, reducing dimensionality fragmented the emotion partition and attenuated the ECS advantage. Crucially, however, it did not abolish it: at perfectly matched dimensionalities, the emotion advantage remained 2.5 to 4.0 times larger than the corresponding low-level advantage, outperforming the low-level models in all 20 random subsets across nearly every film.

**Figure S6.**
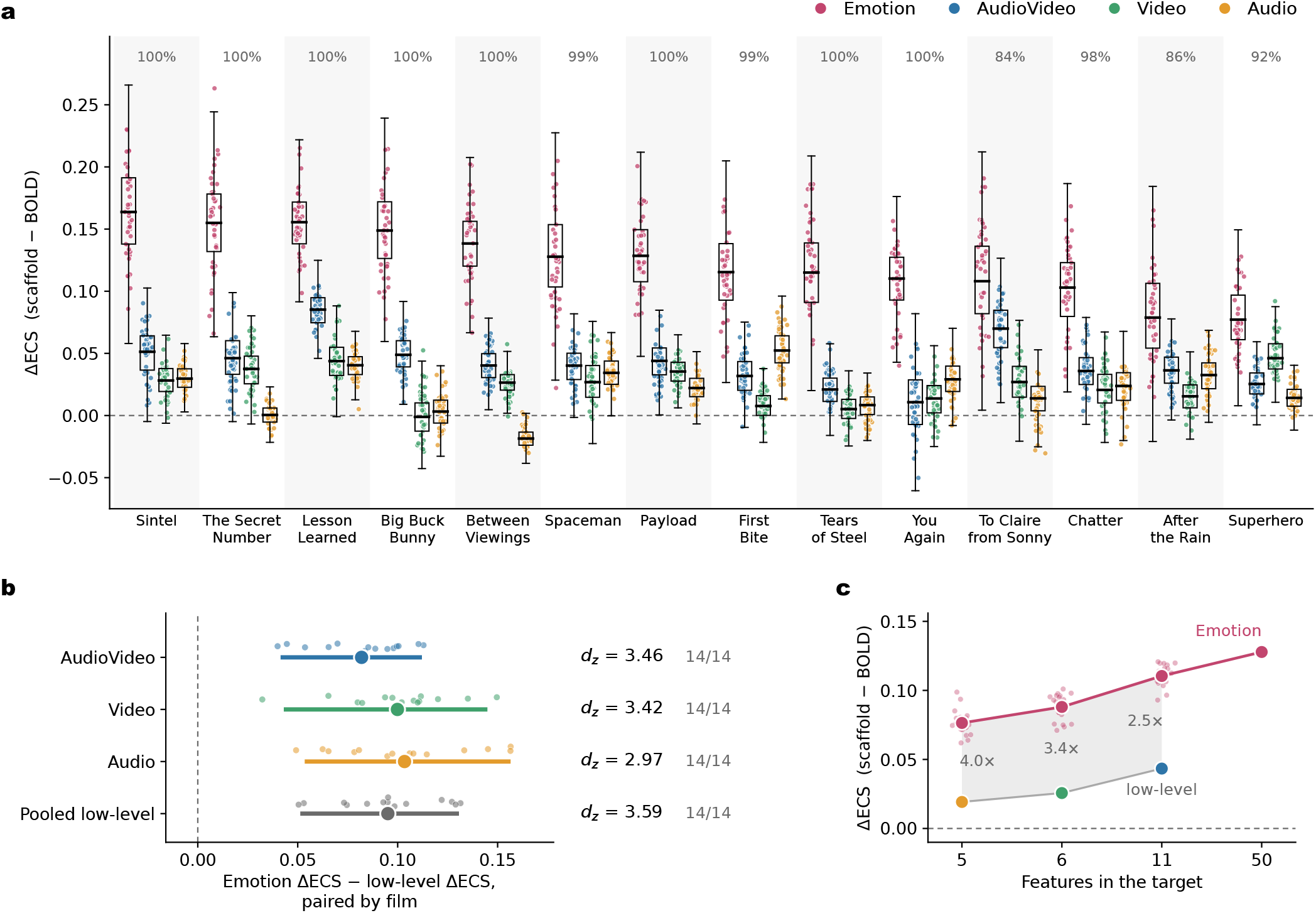
The scaffold’s advantage over regional activity tracks affective content rather than low-level audiovisual streams. For each film, we compared the temporal community structure of a brain representation with a target time series, scoring the match via element-centric similarity (ECS). Throughout, we plot ΔECS (ECS_scaffold_ *−* ECS_BOLD_) to illustrate how much better the homological scaffold recovers a target’s recurrence structure compared to regional activity alone. Four target annotations, all processed identically, were compared: the 50-item emotion annotation and three sets of low-level stimulus descriptors—*Audio* (5 soundtrack features), *Video* (6 image features), and *AudioVideo* (all 11 features). **(a)** ΔECS for each film evaluated across 1,000 paired participant subsamples (drawing 25 of the 30 participants per iteration). Boxes denote the median and interquartile range over all draws; films are ordered sequentially by their emotion ΔECS. The percentages above each column indicate the share of subsamples where the emotion ΔECS exceeded all three low-level baselines (median 99.7%). Because these denote sampling without replacement, they reflect robustness to participant composition rather than statistical *p*-values. **(b)** The same contrast computed with the film as the unit of replication (*n* = 14). Small points represent individual films, the large point indicates their mean, and the bar shows a 95% percentile interval (*d*_*z*_ specifies Cohen’s *d* for paired samples). Averaged over all films, the scaffold’s advantage for the emotion target (0.125) is higher than the audiovisual (0.041), visual (0.024), and auditory (0.020) feature sets, leading in every single film evaluated. **(c)** Control for target dimensionality. Rebuilding the emotion target from 20 random subsets of 11, 6, or 5 regressors (small points, with their mean in bold) predictably attenuates the overall ΔECS, but leaves it 2.5 to 4.0 times higher than the size-matched low-level values. Ultimately, higher-order topology closely follows the narrative’s emotional landscape, rather than responding generically to recurring audiovisual structures.

**Figure S7.**
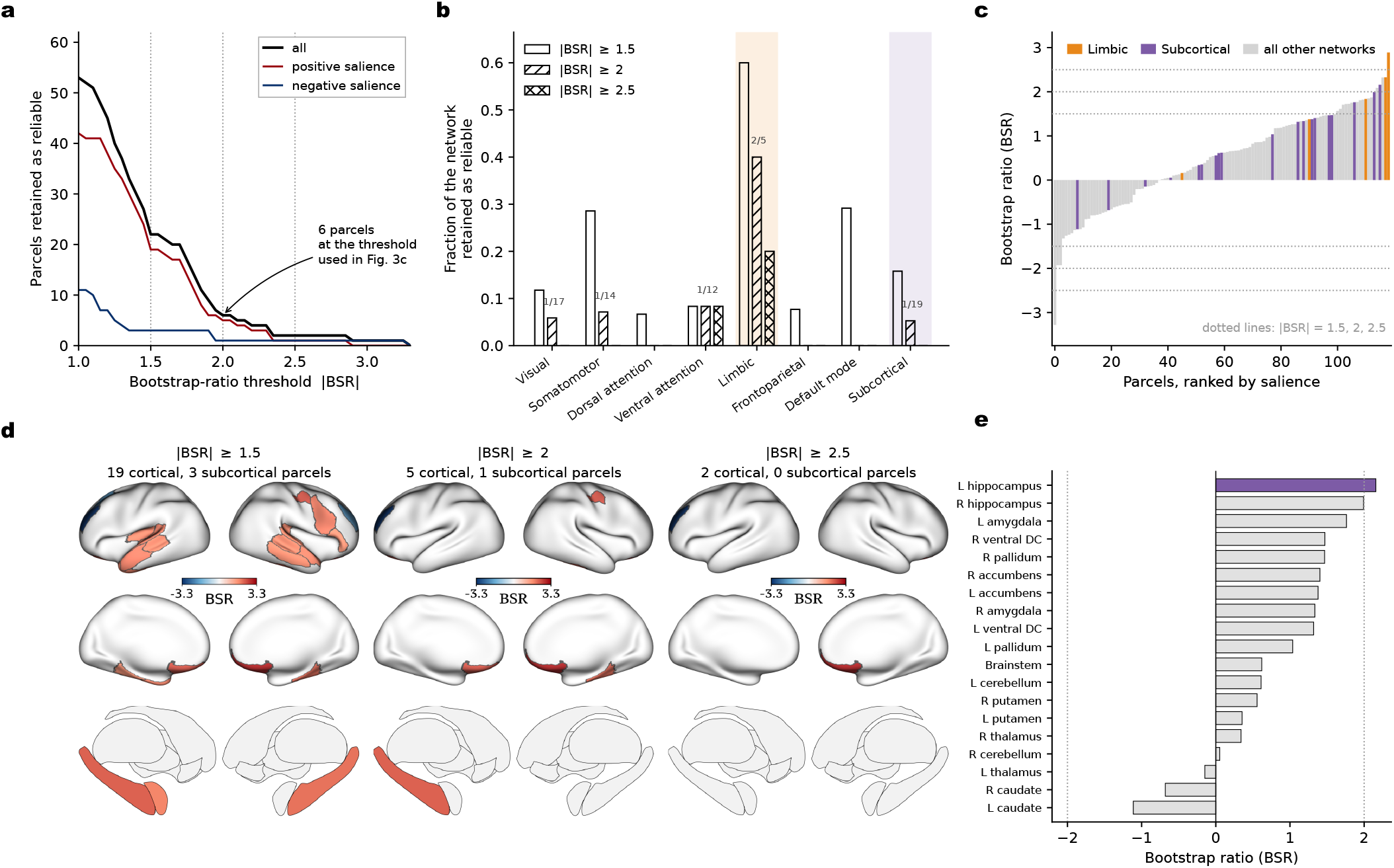
Regional distribution of BOLD saliences across bootstrap-ratio thresholds. Brain saliences for the single significant regional-BOLD latent variable (Fig. 3), shown across a range of bootstrap-ratio criteria. Bootstrap ratios (BSR) are used here to describe the stability and spatial distribution of the salience estimates, rather than as parcel-wise significance tests. **(a)** Number of parcels exceeding the BSR criterion, separated into positive and negative saliences. Dotted lines mark |*BSR*| = 1.5, 2.0, and 2.5; at the threshold used in Fig. 3c ( |*BSR*| ≥ 2.0), 6 of 119 parcels are retained. **(b)** Fraction of parcels retained within each canonical network at three different thresholds. Retention is expressed as a proportion of network size to prevent large networks from dominating; counts above hatched bars denote parcels retained out of total network size at |*BSR*| ≥ 2.0. The limbic (orange) and subcortical (violet) networks are shaded; these hues are distinct from the red and blue encoding salience sign. The five-parcel limbic network retains 40% (2/5), roughly five times any other network, whereas frontoparietal and default-mode retain none. **(c)** Ranked Bootstrap ratios for all 119 parcels, with subcortical and limbic regions highlighted. **(d)** Cortical surfaces (upper) and subcortical structures (lower, medial view) at each threshold on a common color scale. **(e)** Bootstrap ratios for all 19 subcortical parcels; those exceeding |*BSR*| ≥ 2.0 are filled violet. The left hippocampus is the only subcortical structure exceeding |*BSR*| ≥ 2.0 (|*BSR*| = +2.16); the right hippocampus falls just below it (+1.99), followed by the left amygdala (+1.76). The right amygdala (+1.34) ranks below the right ventral DC (+1.47), the right pallidum (+1.47) and the right accumbens (+1.40).

### B. Robustness of the emotion–brain dimensionality (PLSC analysis)

Multivariate cross-decompositions such as PLSC can be sensitive to analytic choices when features greatly outnumber observations [100, 157], and permutation-based significance in PLS can depend on the resampling scheme [158]. We therefore verified that the dimensional dichotomy reported in Fig. 3 is robust along two independent axes—the definition of the scene units and the permutation null (Supplementary Fig. S8).

#### Scene definition

Besides the emotion-derived consensus Louvain scenes (190 scenes), we re-derived scenes in two ways that make no reference to emotion. First, because roughly one fifth of the Louvain scenes span fewer than 20 TRs and therefore yield noisy scene-mean estimates while still entering the model as equally weighted observations, we recomputed the analysis after discarding these short scenes, retaining only Louvain communities of at least 15, 20, or 30 TRs (164, 150, and 123 scenes; the community detection itself was left unchanged and only the smallest communities were set aside). Second, we replaced the Louvain partition entirely with fixed, non-overlapping temporal windows of 15 TRs (486 scenes), 30 TRs (240 scenes), and an *n*-matched 38-TR window. Across all seven definitions, regional BOLD resolved exactly one significant emotion–brain mode, whereas both connectivity representations resolved several (Supplementary Fig. S8**a**). The exact number of connectivity modes did vary with scene granularity—for instance, the standard permutation returned more scaffold modes at finer temporal resolution, an expected consequence of the larger number of observations—but the qualitative one-versus-several dichotomy never changed.

#### Permutation null

Scenes drawn from the same film are not freely exchangeable: they share subject, narrative context, and low-level audiovisual statistics. We therefore complemented the standard permutation (which reassigns scenes across all films) with a *film-restricted* permutation, in which emotion scene labels were shuffled only within each film [158], testing specifically whether emotion–brain coupling exceeds what film-level differences alone provide. Unlike the standard permutation—whose number of significant modes grows with the number of scenes—the film-restricted null yielded a scene-count-invariant estimate: regional BOLD resolved a single mode under every scene definition, while both edge time series and the scaffold robustly resolved three (Supplementary Fig. S8**a**). The weakest modes retained under the standard permutation (for example, the scaffold’s fourth mode) did not survive the film-restricted null and thus reflect between-film structure, whereas the core dichotomy—one regional mode versus multiple connectivity modes—was fully preserved (Supplementary Fig. S8**b**).

**Figure S8.**
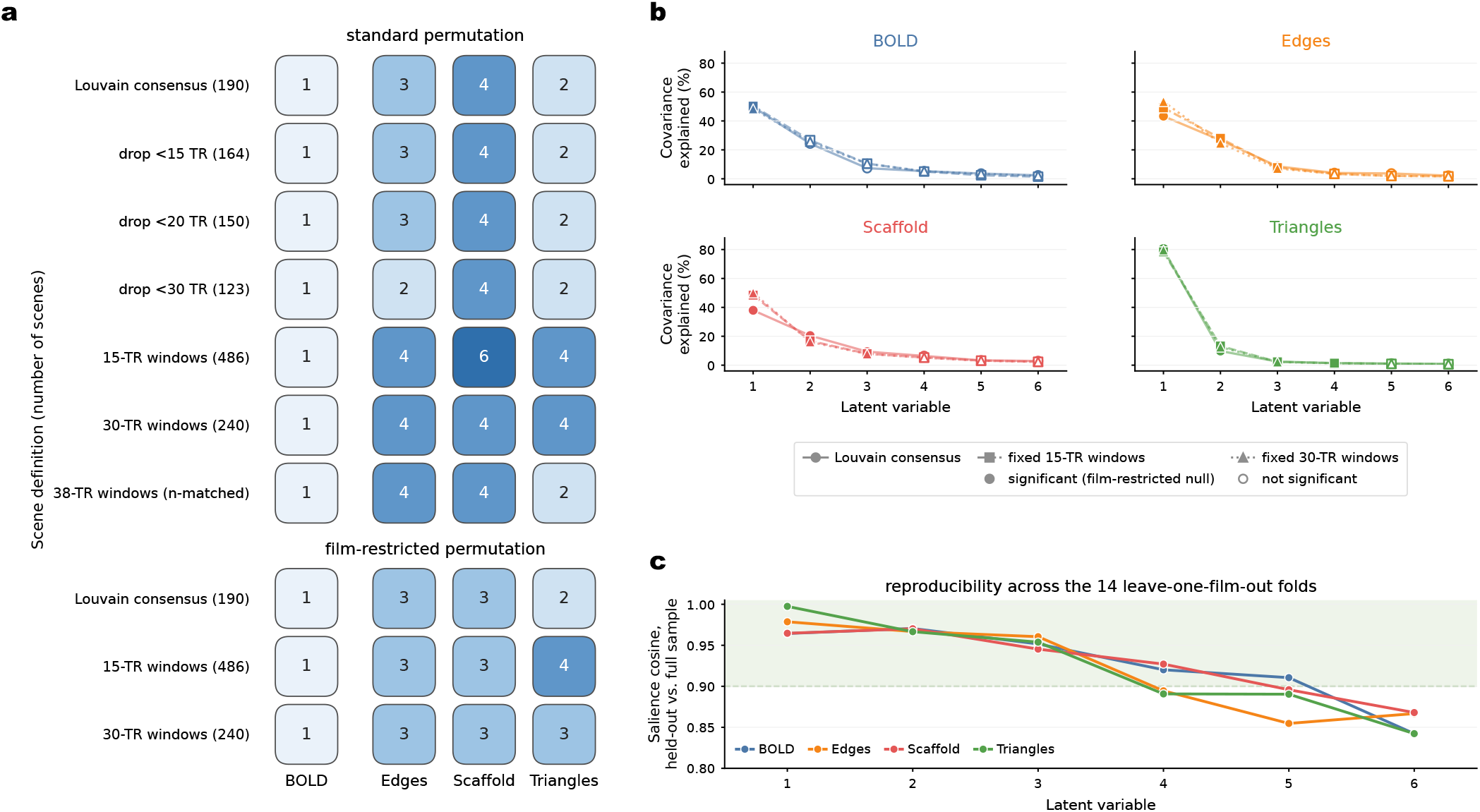
Robustness of the emotion–brain dimensionality to scene definition and permutation null. **(a)** Number of significant PLSC latent variables (permutation *p <* 0.05) for each brain representation across seven scene definitions under the standard permutation (top block: consensus Louvain scenes; Louvain scenes after removing communities shorter than 15/20/30 TRs; fixed non-overlapping 15-, 30-, and *n*-matched 38-TR windows) and under a film-restricted permutation (bottom block; Louvain, 15-, and 30-TR windows). Regional BOLD resolves a single mode throughout, whereas the connectivity representations resolve several; the exact connectivity count varies with scene granularity under the standard permutation but stabilizes at three (for both edge time series and the scaffold) under the film-restricted null. **(b)** Explained-covariance spectra (first six LVs) for the three principal scene definitions; filled markers denote LVs significant under the film-restricted permutation, open markers non-significant. **(c)** Reproducibility of the brain saliences: for each latent variable, the cosine similarity between the brain salience estimated with one film held out and that of the full-sample decomposition, averaged over the 14 leave-one-film-out folds (shaded band, cosine *>* 0.9). The leading modes of every representation are highly reproducible despite the large number of connectivity features relative to scenes.

**Figure S9.**
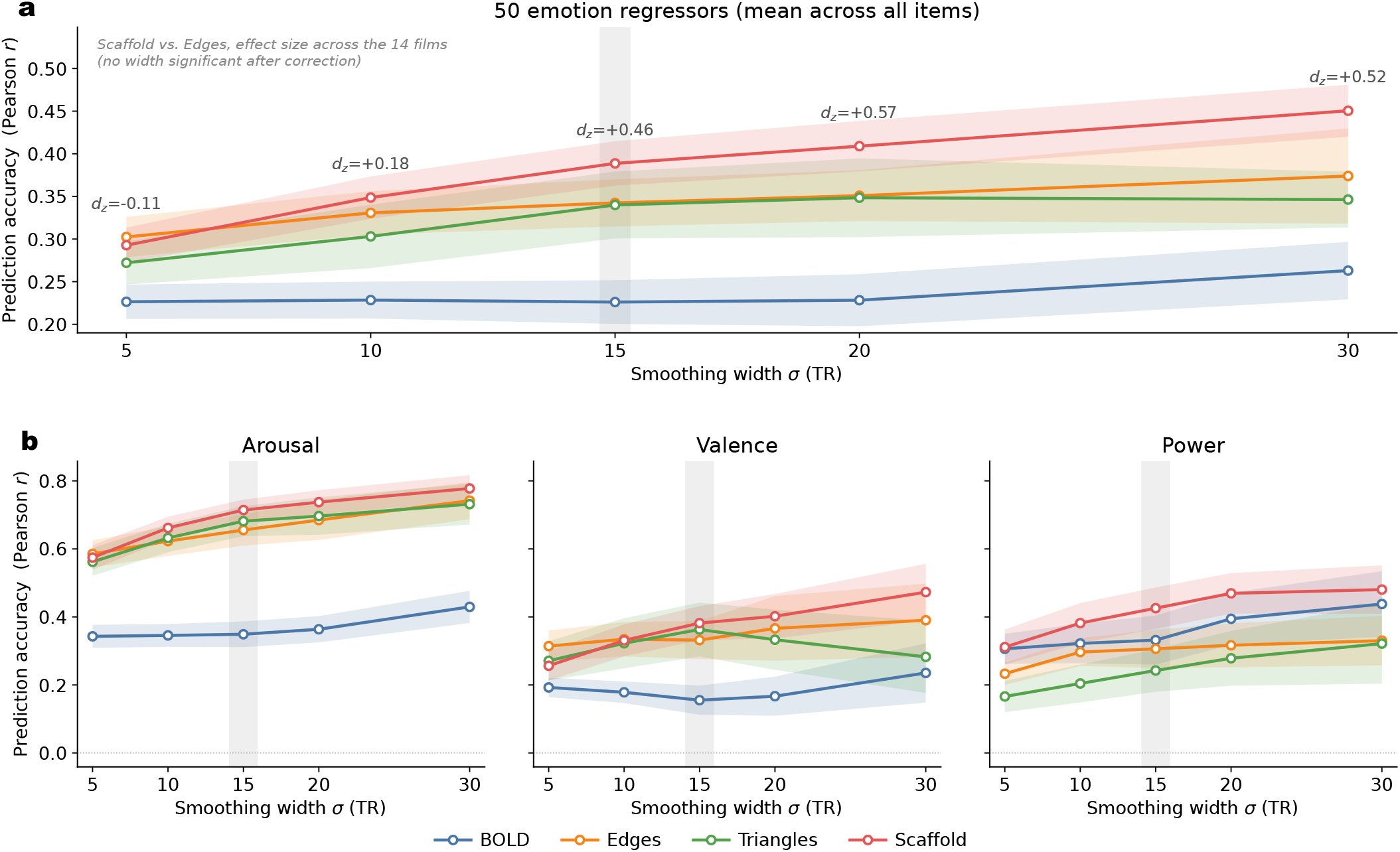
Emotion-prediction accuracy is robust to the temporal smoothing width. Leave-one-film-out kernel-ridge-regression accuracy for the four brain representations across Gaussian smoothing widths *σ* (TR = 1.3 s). Accuracy is reported as the mean Pearson correlation *r* across the 14 held-out films, with shaded bands denoting ±1 standard error across films. The gray band marks *σ* = 15 TR, the smoothing width used in the main analyses. **(a)** Prediction accuracy averaged across the 50 emotion regressors. Values above the curves give the film-level effect size of the scaffold-minus-edges difference (*d*_*z*_ over the 14 held-out films; exact sign-flip test, FDR-corrected across the five widths).. Across the full emotional space, performance increased with smoothing for all representations, but the relative ordering was preserved: the homological scaffold provided the highest accuracy from *σ* = 10 TR onward, whereas regional BOLD remained the least predictive representation across all widths. **(b)** Prediction accuracy for the three affective dimensions: arousal, valence, and power. Connectivity-based representations consistently outperformed regional BOLD for arousal and showed broadly similar performance to one another, whereas valence and power showed weaker and less differentiated prediction across representations. Overall, the qualitative pattern was stable across medium and large smoothing widths. Notice that unlike the fully nested procedure used in Fig. 4, this supplementary sensitivity analysis selects a single ridge penalty for each representation and smoothing width based on performance across all 14 leave-one-film-out folds. Consequently, the held-out films contribute to hyperparameter selection. The effect sizes and associated *p*-values shown here should therefore be interpreted descriptively as part of the smoothing-sensitivity analysis, rather than as independent confirmatory tests of the main Fig. 4 result.

### C. Emotion prediction robustness and dimensionality analyses

#### Dimensional constraints on predictive readouts

Item-level emotion decoding in our main analysis relied on kernel ridge regression (KRR) with a linear kernel, solved in dual form, where the ridge penalty *λ* ∈ {10^*−*3^, 10^*−*2^, …, 10^4^} was tuned per fold via nested leave-one-film-out cross-validation. To confirm that the observed differences between neural representations reflect their intrinsic predictive capacity rather than artifacts of this specific estimator, we repeated the 50-emotion decoding using complementary methods that impose progressively stronger dimensionality constraints on the readout (Supplementary Fig. S10**a**). This spectrum of estimators represented a gradient of decreasing model complexity, ranging from approaches that retain nearly the full feature set to aggressively compressed two-dimensional summaries. Specifically, we evaluated the connectome-based support-vector readout (CPM-SVR) [59, 105], sparse *ℓ*_1_-regularized regression (LASSO) [159], partial least squares regression (PLSR) [99], and canonical connectome-based predictive modeling (CPM) [104, 145]. All models were evaluated using identical participant-averaged predictors—comprising regional BOLD activity alongside the three edge-level connectivity representations—and targeted the same smoothed continuous emotion trajectories. For PLSR (`scikit-learn PLSRegression` with no internal scaling), predictions were restricted to a low-rank projection by tuning the number of latent components over the set {1, 2, 3, 5, 8, 12, 20} . Conversely, the canonical CPM pipeline severely compressed the connectome by thresholding features at the top decile of training-set Pearson correlations ( |*r*| at or above the 90th percentile) and summing them into strictly two positive and negative network-strength signals. These two summaries were then used to fit an ordinary least squares linear model with an intercept (*ŷ*(*t*) = *β*_0_ + *β*_+_ *S*_+_(*t*) + *β*_*−*_ *S*_*−*_(*t*)). We also tested the exact CPM-SVR readout used by Ke *et al*. [59], applying a non-linear radial basis function support-vector regression to all connections surviving a *p <* 0.01 training-set threshold, evaluating strict generalization to new, unseen films.

#### Temporal integration and effective dimensionality

Although canonical CPM was originally developed for sliding-window functional connectivity, no sliding window was required here because all three connectivity representations are inherently resolved at the level of individual TRs (1.3 s). Thus, the only temporal integration applied was the identical Gaussian smoothing (*σ* = 15 TR) used in the main analysis, which isolated the relevant slow behavioral timescales while matching the temporal grain of the neural and affective data. To explicitly estimate the effective dimensionality of the predictive code within this temporally smoothed data, we fit an independent LASSO regression (`scikit-learn` Lasso via coordinate descent, tolerance 10^*−*3^, maximum 3000 iterations) for each emotion along a warm-started penalty path of *α* ∈ {0.02, 0.01, 0.005, 0.002, 0.001} .

By fixing the penalty to the value that maximized the mean leave-one-film-out accuracy, we used the mean number of non-zero coefficients as a direct measure of how distributed the predictive code is for each distinct neural representation (Supplementary Fig. S10**b**).

#### Cross-validation and statistical inference

Across all estimators, generalization was strictly assessed using a leave-one-film-out cross-validation scheme across the 14 narratives. Feature selection for CPM and CPM-SVR was performed within each training set; for PLSR and LASSO, the number of components or the penalty was chosen from the complete 14-fold accuracy curve, so these two rungs are descriptive capacity analyses. Prediction accuracy was quantified as the Pearson correlation (*r*) between the predicted and observed emotion time courses in the held-out film. Because correlation-based prediction of autocorrelated time series can be artificially inflated, CPM accuracy was rigorously evaluated against a whole-film circular-shift null distribution. This null procedure preserved each item’s temporal autocorrelation while safely destroying its temporal correspondence to the brain, allowing us to calculate robust empirical significance and confirm that all models capture genuine brain–emotion alignment (Supplementary Fig. S10**c**). Finally, statistical comparisons mirrored the main text: representations were compared with the held-out film as the unit of replication (the accuracy difference averaged over the 50 items within each film, *n* = 14). We report the film-level effect size *d*_*z*_ and the number of films favoring the scaffold. Exact sign-flip tests, BH-corrected across the five estimators, were not significant for any estimator (KRR *p*_*FDR*_ = 0.073; all others *p*_*FDR*_ = 0.58), so these comparisons are presented descriptively. The 50 emotion items are strongly intercorrelated, so a paired test across items would count one shared signal many times; because these comparisons additionally span five estimators with only 14 films each, they are presented descriptively. Dimensional comparisons used the Nadeau–Bengio corrected resampled *t*-test [103] to account for training-set overlap.

**Figure S10.**
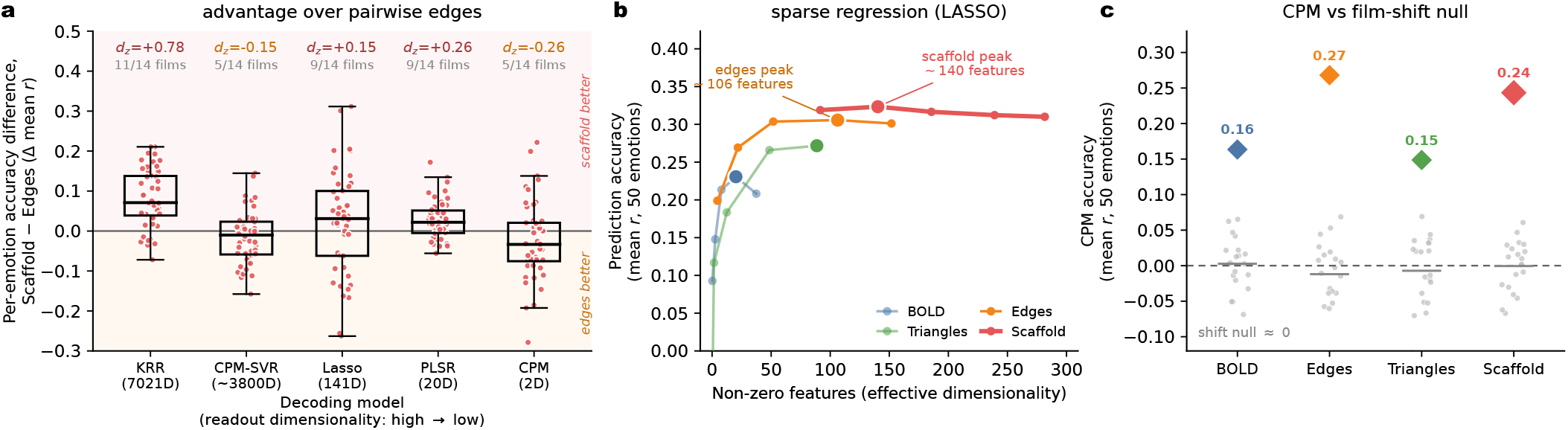
The scaffold’s predictive advantage reflects a high-dimensional, distributed code. Evaluation of the 50-emotion decoding of Fig. 4 across alternative predictive models. **(a)** Per-emotion accuracy difference between the scaffold and pairwise edge time series across five estimators, ordered from left to right by decreasing effective dimensionality. The largest scaffold’s advantage over edges is present under kernel ridge regression (KRR), which uses the full 7021-dimensional feature space (11 of 14 films, *d*_*z*_ = 0.78), and is absent for every estimator that reduces or thresholds the features, including the non-linear CPM-SVR approach [59]. This indicates that the scaffold’s decoding power relies on a distributed long tail of weak connections, which are systematically discarded by feature selection. Boxes indicate the interquartile range with a median center line, and whiskers extend to 1.5 times the interquartile range. Boxes show the 50 per-emotion differences; values above the boxes give the film-level effect size *d*_*z*_ and the number of films in which the scaffold was higher. Because the 50 emotion items are not independent, no significance markers are drawn. **(b)** Sparse linear regression (LASSO) prediction accuracy as a function of the number of retained non-zero features. Edge time series reach their peak accuracy at roughly 106 non-zero features, whereas the scaffold needs about 140. This confirms that the scaffold’s predictive information is distributed across many weakly informative connections rather than concentrated in a few. **(c)** Canonical predictive modeling (CPM) accuracy compared to a whole-film circular-shift null. By compressing the connectivity profile into just two scalar components, CPM eliminates the scaffold’s advantage (edges higher in 9 of 14 films, *d*_*z*_ = −0.26). Nevertheless, every neural representation performs well above this autocorrelation-preserving null. This confirms that our models capture genuine brain–emotion alignment and that cross-representation comparisons remain unbiased by differing chance baselines. Notice that for the CPM-SVR approach, as in Ke *et al*. [59], the solver was capped at 1000 iterations, which it reached for every fit.

**Figure S11.**
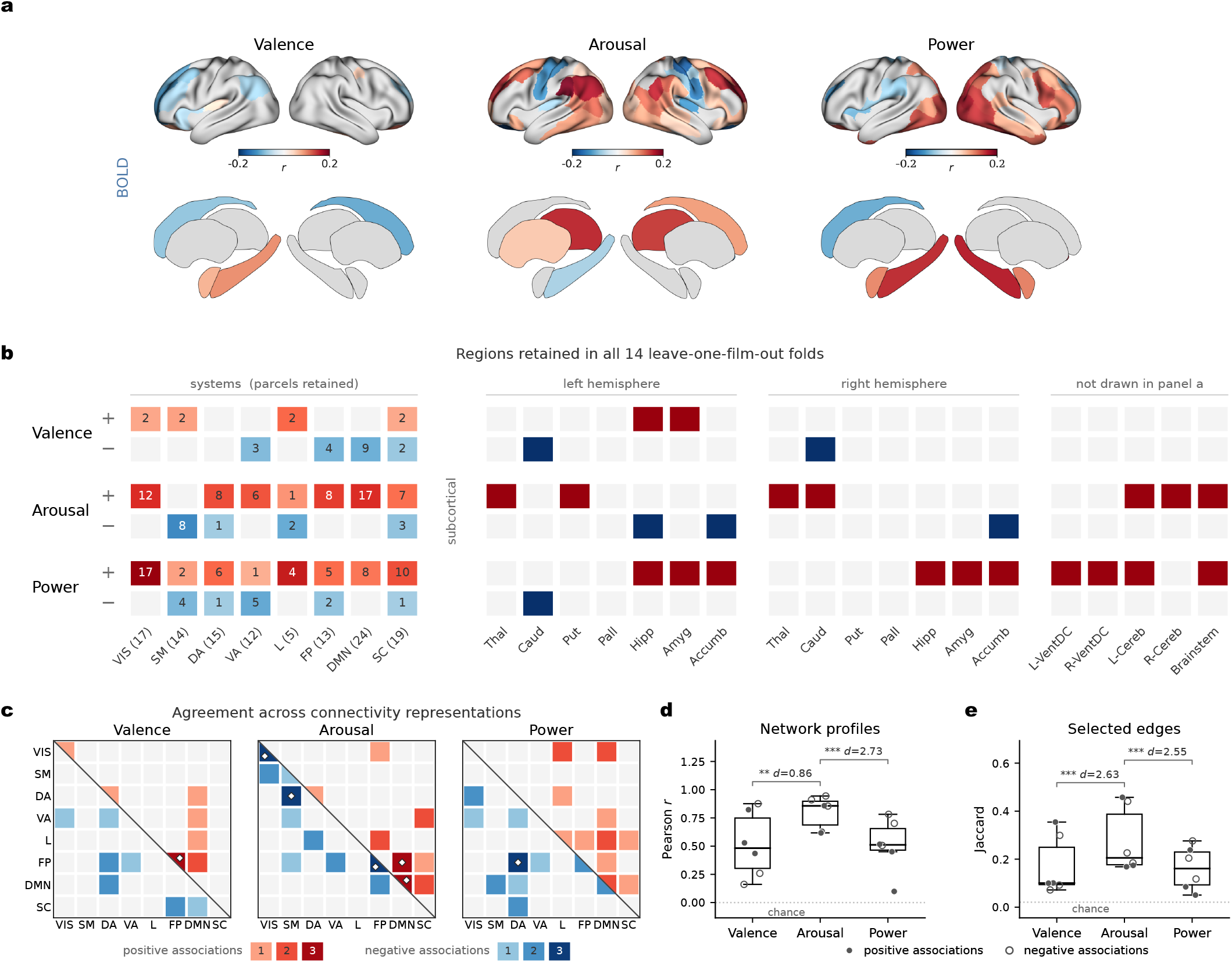
Network and regional mapping reveals that cross-representation convergence is specific to arousal in the Emo-FilM dataset. **(a)** Regional BOLD maps, rendered on the cortical surface and on subcortical structures (medial view). Color indicates the fold-averaged correlation with the affective dimension (valence, arousal, power), exclusively for regions retained across all 14 leave-one-film-out folds at a one-sided *p <* 0.01 threshold (matching the selection rule in Fig. 5**e**). The subcortical atlas used for rendering covers only 14 structures, meaning the ventral DC, cerebellum, and brainstem are excluded from this visualization. **(b)** The explicit region list underlying the maps in **(a)**. Rows denote regions exhibiting consistent negative ( *−* ) and positive (+) associations with each dimension. Columns represent the Yeo networks (with their total significant parcel count in brackets), followed by the 7 subcortical structures of the left and right hemispheres, and finally the five structures omitted from the visual rendering in **(a)**. Each network cell is shaded according to the fraction of its parcels retained across all 14 folds, with the absolute count printed inside. **(c)** Connectivity representation. Features were ranked within each training fold by their signed correlation with the target dimension. The strongest positive and negative associations (specifically, the top and bottom 10% of features) were retained separately and intersected across the 14 folds. The surviving edges were then summarized over the seven Yeo networks and the subcortical system. We tested the enrichment of each network pair relative to a one-sided hypergeometric baseline, applying Benjamini–Hochberg FDR correction across the 36 pairs separately for each representation and direction of association. Each cell is shaded to indicate how many of the three connectivity representations (edges, scaffold, triangles) achieved significance (*p*_FDR_ *<* 0.05) for that specific pair. The upper triangle displays positive associations, while the lower triangle displays negative associations, with white diamonds highlighting network pairs successfully recovered by all three representations. **(d)** Correlation between the 36 network-profile values of two representations (yielding one point per representation pair and association direction; filled = positive associations, open = negative associations), with boxes denoting the median and quartiles. **(e)** Jaccard overlap between the selected edge sets for these same representation pairs. Dotted lines indicate the baseline overlap expected purely by chance, given the selection sizes. For both **(d)** and **(e)**, significance was determined via a delete-one-film jackknife comparison of the 14-film intersection, FDR-corrected across the four comparisons (*: p*_*FDR*_ *<* 0.05,*: : : p*_*FDR*_ *<* 0.001), and reported with Cohen’s *d*.

### D. Correspondence between the decoder and the encoding map

Our predictive models operate as backward (discriminative) decoders, learning a mapping from brain features to emotional targets. As established by Haufe et al. [106], the weight vector of such models cannot be directly interpreted as a spatial map of the underlying neural signal. To cancel shared noise, these models often assign large weights to features unrelated to the target (suppressors) and near-zero weights to features that are highly correlated with it. This spatial distortion is particularly acute for connectivity representations, where the 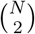 edge features are highly collinear, and is further compounded by ridge regularization, which distributes weights arbitrarily across correlated variables. Consequently, the raw backward weight map fails to provide a valid neuroanatomical description of the decoded substrate.

To resolve this, a backward model can be made interpretable by computing its forward (activation) pattern, *A* = Cov(*X*) *w*. For a single target, this quantity equals the covariance between each feature and the model’s prediction, 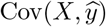, effectively capturing how strongly each feature actually co-varies with the decoded signal. The encoding map used for our spatial localization — defined as each feature’s correlation with the target — approximates this interpretable forward quantity; the residual departure from unity (cosine *<* 1) reflects the ridge shrinkage that separates Cov(*X*)*w* from the raw feature–target covariance. Computed directly from the data and independently of any fitted model, this encoding map represents the standard approach to visualizing a linear decoder’s spatial representation and coincides with the feature-selection logic of connectome-based predictive modeling [59, 104, 145]. To empirically verify that our encoding map represents the signal exploited by the decoder, we compared the two approaches within every neural representation and cross-validation fold using identical mean-centered training data. From the fitted linear-kernel KRR model, we recovered the primal weights *w* = *X*^*⊤*^***α***, the in-fold prediction 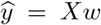, and the forward pattern 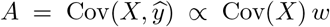. The encoding map was then compared with both the forward pattern *A* and the raw backward weights *w* through three metrics: cosine similarity of the full maps, Pearson correlation of the network-aggregated 8 8 patterns, and the Jaccard overlap of the top-10% selected features. Because the cross-validation estimates originate from overlapping training sets, forward-versus-backward differences were evaluated using the Nadeau–Bengio corrected resampled *t*-test [103], with *p*-values FDR-corrected across representations and effect sizes reported as paired Cohen’s *d*. As controls, the primal weights recomputed here reproduced the weights of the decoding analysis (cosine 0.95–0.99), and the recomputed encoding maps reconstructed the feature selections of Fig. 5**e** exactly (Jaccard = 1.00 for both the positive and the negative selections of every connectivity representation).

The results reported in Fig. S12 confirmed our rationale: the forward pattern reproduced the encoding map (cosine similarity 0.91–0.99; top-10% edge overlap 0.51–0.70), whereas the raw weights were markedly less aligned (cosine 0.24–0.70; overlap 0.08–0.24). This divergence was most pronounced for the triangle and scaffold representations. Therefore, our correlation-based spatial localization provides a non-circular description of the signal the decoder exploits, obtained without reference to its raw weights.

**Figure S12.**
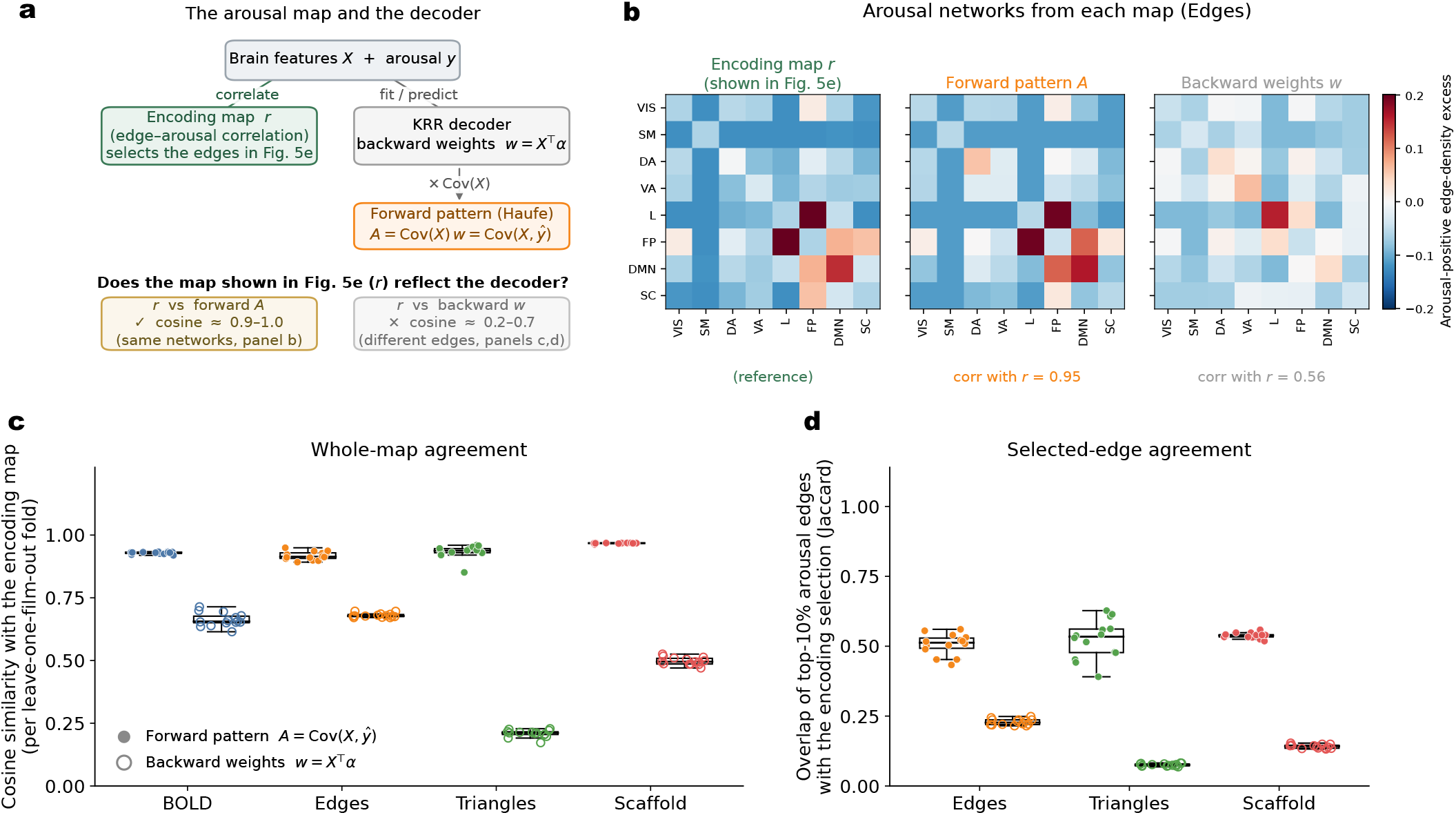
The arousal map reflects the decoder’s forward pattern, not its raw weights. **(a)** Schematic of the spatial maps evaluated. For brain features *X* and arousal time course *y*, spatial localization relies on an *encoding map r*, defined as each feature’s correlation with arousal. Kernel ridge regression, however, fits *backward* weights *w* = *X*^*⊤*^***α***. Its interpretable *forward* (activation) pattern is derived via the Haufe transform, *A* = Cov(*X*) *w* = Cov(*X, y*). We compare the independent encoding map (*r*) against both the decoder’s forward pattern (*A*) and its raw weights (*w*). **(b)** The arousal-positive network enrichment (8 *×* 8 edge-density excess over seven cortical networks and one subcortical system; edge representation), reconstructed from each of the three maps. The encoding map and the forward pattern highlight identical systems—limbic– frontoparietal, frontoparietal–default-mode, and frontoparietal–subcortical (spatial correlation *r* = 0.95), whereas the raw weights give a diffuse, weaker pattern (*r* = 0.56). **(c)** Per-fold cosine similarity of the forward pattern (filled markers) and the backward weights (open markers) with the encoding map across the four neural representations. The forward pattern is near-collinear with the encoding map (cosine 0.92–0.97), while the backward weights align far less (0.21–0.69), particularly for the sparser triangle and scaffold representations. **(d)** Overlap (Jaccard) of the top-10% arousal edges selected by each decoder map with those selected by the encoding map. The forward pattern recovers most of the encoding selection (0.50–0.71); the backward weights recover little (0.05–0.24).

**Figure S13.**
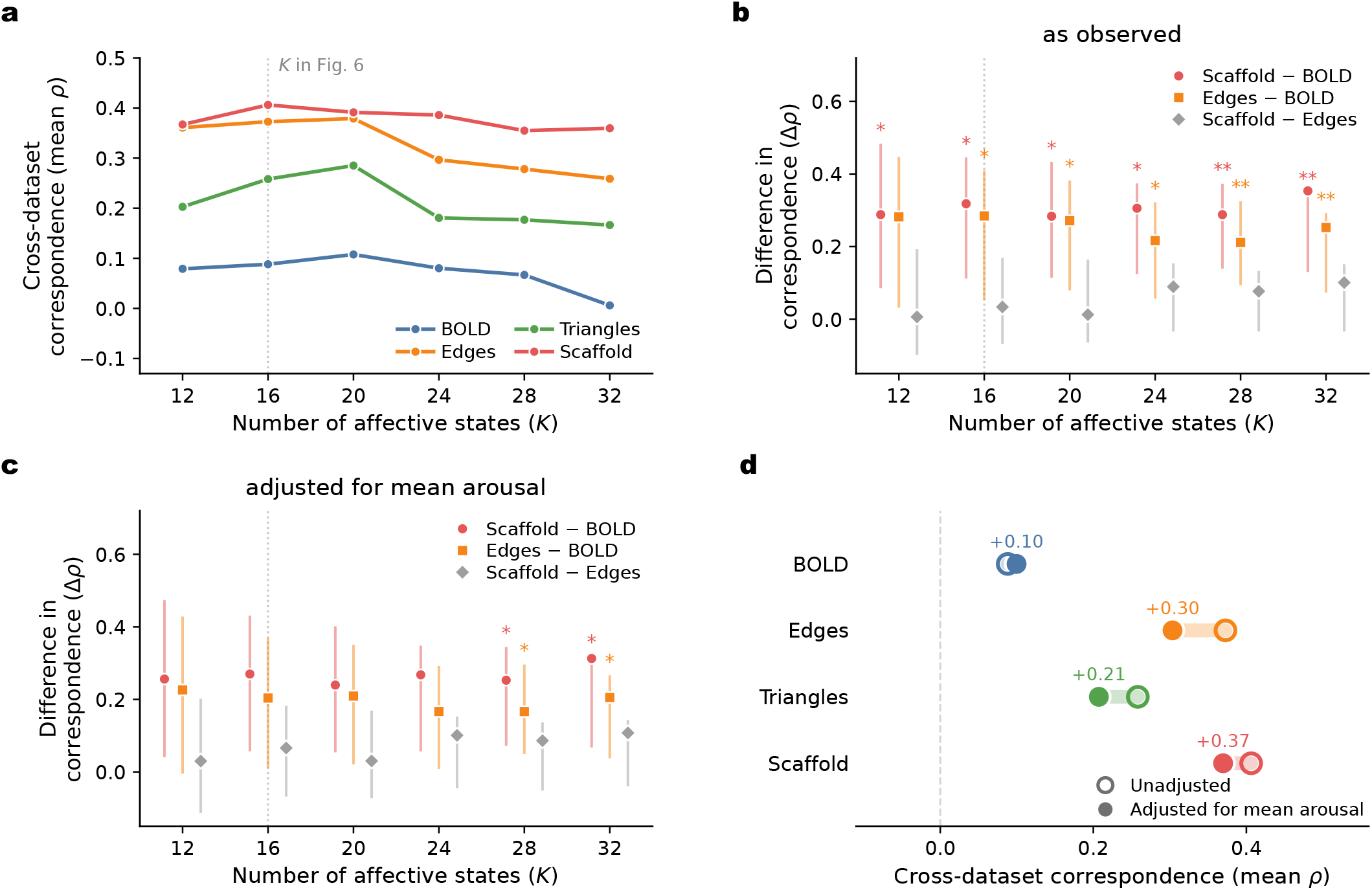
Cross-dataset affective-state geometry across state-space resolutions. Affective states were defined by a single *k*-means partition of the four datasets’ pooled, standardized valence–arousal clouds into *K* clusters, each time point being assigned to its nearest centroid; the main analysis uses *K* = 16 (dotted line). States sampled by fewer than 10 volumes in a dataset are not considered from that dataset’s RDM. All values are averaged across the six dataset pairs. Intervals and *p* values were obtained using hierarchical resampling, with acquisition units resampled with replacement and 90-s circular moving blocks within units; state means, RDMs, and affect-distance covariates were recomputed in every replicate (1,000 bootstraps, 1,000 state-label permutations, FDR-BH correction across six contrasts). **(a)** Cross-dataset geometric correspondence for each neural representation. The same ordering is preserved across resolutions, with scaffold and edges consistently showing the strongest correspondence and regional BOLD remaining close to zero. **(b)** Pairwise representation contrasts. Scaffold exceeds regional BOLD at every resolution (Δ*ρ* = 0.28–0.35, all *q* ≤ 0.048), while edges exceeds BOLD from *K* = 16 onward. Scaffold and edges do not differ significantly at any resolution. **(c)** The same contrasts after accounting for state-pair differences in mean arousal. This adjustment attenuates both connectivity-based effects (Scaffold–BOLD Δ*ρ* = 0.24–0.31; Edges–BOLD Δ*ρ* = 0.17–0.23), with significance retained only at the finest resolutions (*K* = 28 and 32). Scaffold and Edges remain statistically indistinguishable across all resolutions. **(d)** Effect of mean-arousal adjustment at *K* = 16. Open circles show the unadjusted means, filled circles the adjusted means.

**Figure S14.**
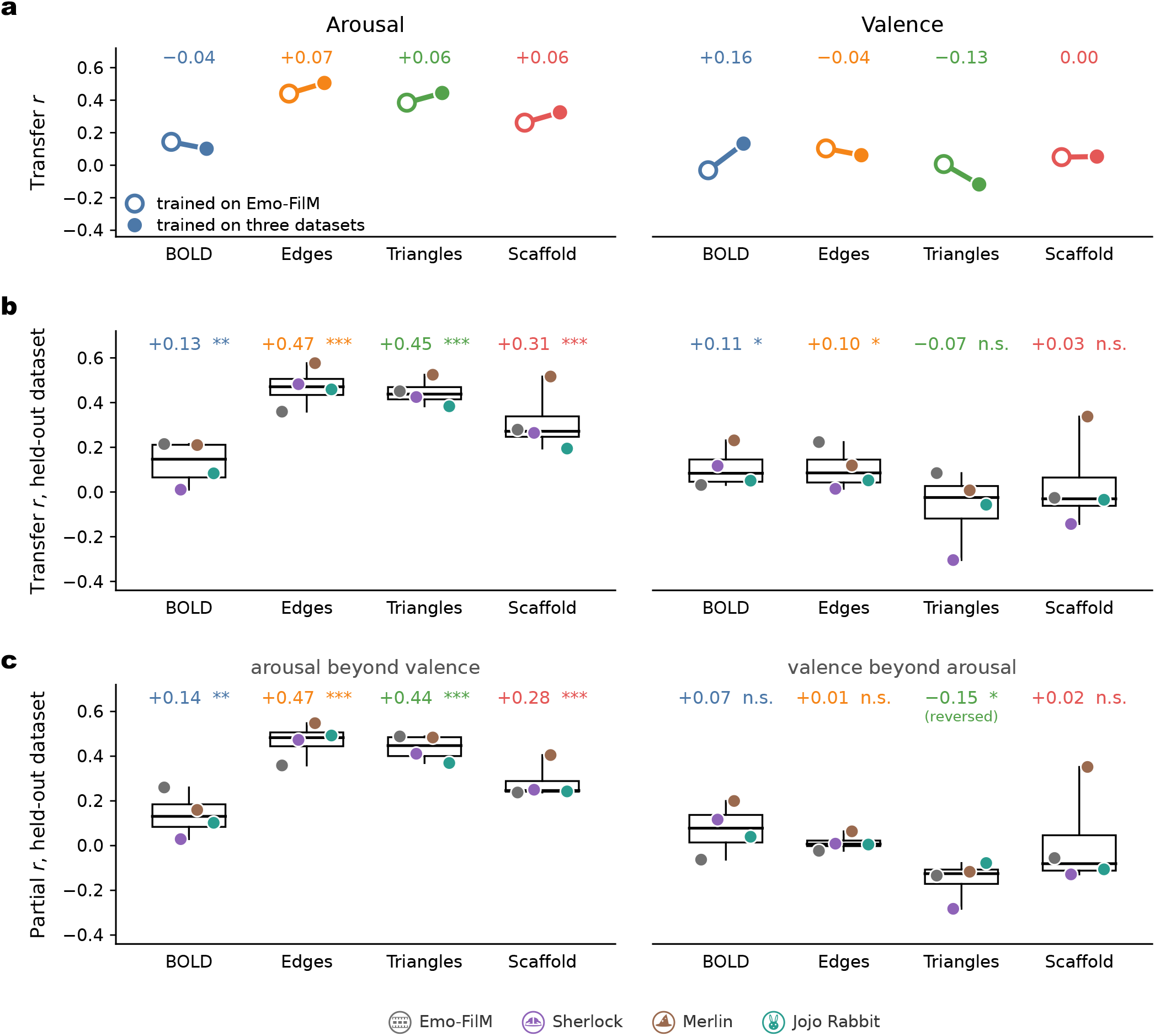
Cross-narrative training strengthens arousal but not valence generalization. To assess whether broader training improves cross-dataset generalization, we compared zero-shot transfer, in which models were trained only on Emo-FilM and applied without refitting to the external datasets, with leave-one-dataset-out (LODO) validation, in which models were trained on three datasets and tested on the fourth. Transfer was quantified as the Pearson correlation between predicted and observed affective trajectories. **(a)** Change from zero-shot to LODO across the three external datasets (Sherlock, Merlin and Jojo Rabbit). Lines connect the zero-shot mean (open circles) and LODO mean (filled circles) for the same held-out datasets; annotations indicate the corresponding change. Arousal transfer increases for all three connectivity representations (+0.06 to +0.07) but not for regional BOLD ( −0.04). Valence shows no consistent improvement (BOLD +0.16, edges ™0.04, triangles −0.13, scaffold 0.00). **(b)** LODO transfer across all four held-out datasets. Points denote individual folds and boxplots summarize the median, interquartile range, and full range; values above each box indicate the mean across folds. Significance was assessed against 1,000 phase-randomized surrogates per held-out dataset, preserving temporal autocorrelation, and FDR-corrected across the eight representation-by-dimension tests. Arousal transfers significantly for every representation, including scaffold (*r* = 0.31, *p*_FDR_ *<* 0.001) and regional BOLD (*r* = 0.13, *p*_FDR_ = 0.006). Valence reaches significance only for regional BOLD (*r* = 0.11, *p*_FDR_ = 0.035) and edges (*r* = 0.10, *p*_FDR_ = 0.040). **(c)** The same LODO predictions evaluated by partial correlation after removing the alternate affective dimension from both prediction and target in the held-out dataset. Significance was assessed against phase-randomized surrogates of the residualized target and corrected as in **(b)**. Arousal remains significant for every representation after controlling for valence (all *p*_FDR_ *≤* 0.003), whereas valence shows no positive transfer beyond arousal (regional BOLD *r* = 0.07, *p*_FDR_ = 0.27; edges *r* = 0.01, *p*_FDR_ = 0.99; scaffold *r* = 0.02, *p*_FDR_ = 0.53). Triangles show a significant negative partial association (*r* = −0.15, *p*_FDR_ = 0.021; labelled “reversed”). Thus, the apparent valence transfer in **(b)** is attributable to variance shared with arousal.

**Figure S15.**
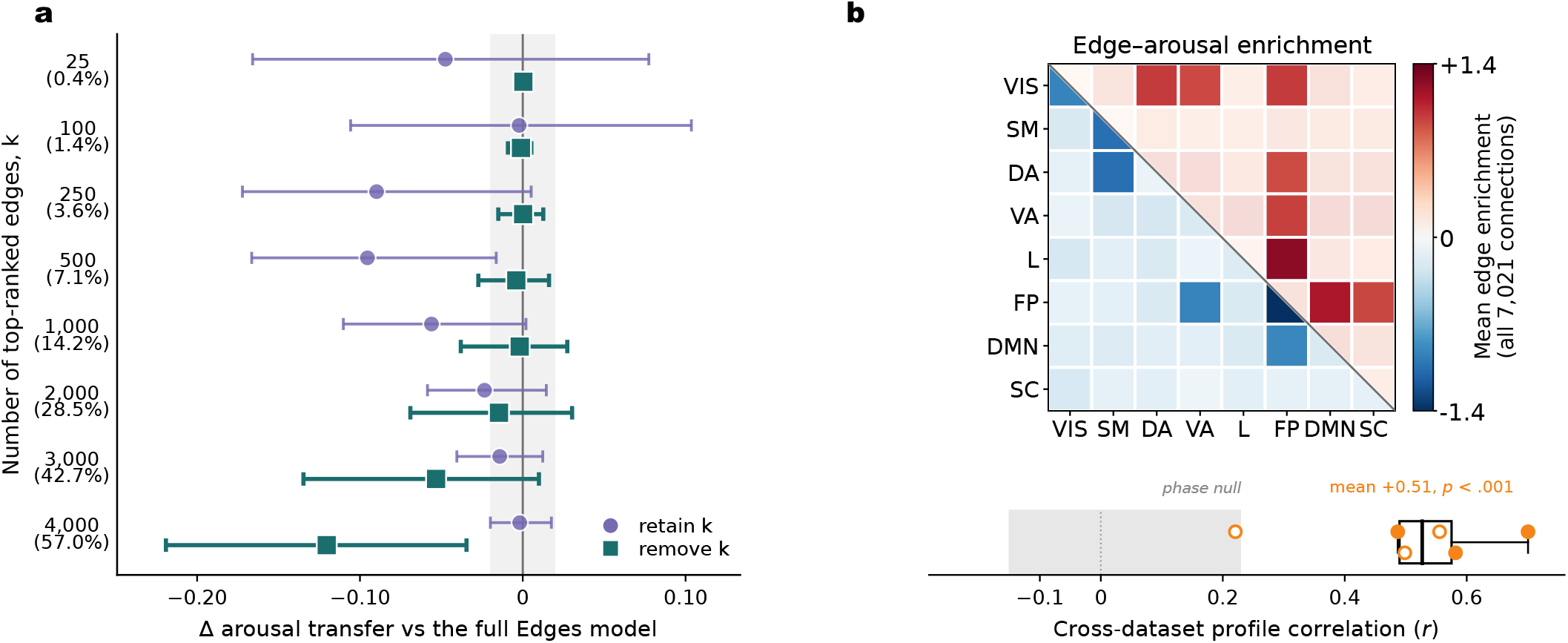
Edge-based arousal transfer reflects a distributed and reproducible network-level architecture. All analyses use pairwise edge time series and the arousal dimension. **(a)** Dependence of zero-shot arousal transfer on the predictive edge set. Performance is shown relative to the full 7,021-edge model when the top *k* Emo-FilM ranked edges are either retained (violet) or removed before refitting on the complementary set (teal). Whiskers denote paired 95% confidence intervals obtained from the same moving-block resamples used for the full and reduced models; the shaded region marks the pre-specified practical-equivalence range ( ± 0.02 *r*). Removing the top 25–250 edges leaves performance practically equivalent to the full model, whereas removing 4,000 edges (57%) reduces transfer (Δ*r* = −0.120, 95% CI [ −0.219, −0.035]), indicating that predictive information is broadly distributed rather than concentrated in a small set of connections. **(b)** Cross-dataset consensus architecture of arousal-predictive coupling, aggregated into Yeo network-pair blocks. The upper and lower triangles show positively and negatively predictive coupling, respectively, in the uncentred four-dataset mean of the sign-split profiles. Cross-dataset similarity is quantified on the corresponding signed 36-block profiles, with one point per dataset pair (filled, pairs including Emo-FilM; open, external–external pairs) and the boxplot summarizing all six comparisons. The grey band denotes the 5–95% interval of 1,000 phase-randomized surrogates. All six pairwise correlations are positive (*r* = 0.22–0.70; mean *r* = 0.51, phase-randomized *p <* 0.001), including the three external–external pairs (mean *r* = 0.43), so the shared architecture is not driven solely by comparisons involving the training dataset. Notice that the matrix shows the sign-split profile (72 cells, positive and negative coupling kept apart); the correlations tested are computed on the signed 36-block profile, the quantity reported in the main text.

## References

[1] A. Santoro, F. Battiston, G. Petri, and E. Amico, “Higher-order organization of multivariate time series,” Nature Physics 19, 221–229 (2023).

[2] K. R. Scherer, “What are emotions? And how can they be measured?” Social Science Information 44, 695–729 (2005).

[3] L. F. Barrett, B. Mesquita, K. N. Ochsner, and J. J. Gross, “The Experience of Emotion,” Annual Review of measured mean arousal, it does not remove all arousalrelated structure. Psychology 58, 373–403 (2007).

[4] L. Pessoa, “A Network Model of the Emotional Brain,” Trends in Cognitive Sciences 21, 357–371 (2017).

[5] R. Adolphs, L. Mlodinow, and L. F. Barrett, “What is an emotion?” Current Biology 29, R1060–R1064 (2019).

[6] A. Moors, P. C. Ellsworth, K. R. Scherer, and N. H. Frijda, “Appraisal Theories of Emotion: State of the Art and Future Development,” Emotion Review 5, 119–124 (2013).

[7] P. Kuppens, “It’s About Time: A Special Section on Affect Dynamics,” Emotion Review 7, 297–300 (2015).

[8] L. F. Barrett, “The theory of constructed emotion: An active inference account of interoception and categorization,” Soc. Cogn. Affect. Neurosci. 12, 1–23 (2017).

[9] A. S. Cowen and D. Keltner, “Semantic Space Theory: A Computational Approach to Emotion,” Trends in Cognitive Sciences 25, 124–136 (2021).

[10] P. Ekman and W. V. Friesen, Unmasking the Face: A Guide to Recognizing Emotions from Facial Clues (ISHK, 1975).

[11] P. Ekman, “An argument for basic emotions,” Cognition and Emotion 6, 169–200 (1992).

[12] J. A. Russell and A. Mehrabian, “Evidence for a threefactor theory of emotions,” Journal of Research in Personality 11, 273–294 (1977).

[13] J. A. Russell, “A circumplex model of affect.” Journal of Personality and Social Psychology 39, 1161–1178 (1980).

[14] A. Mehrabian, “Pleasure-arousal-dominance: A general framework for describing and measuring individual differences in Temperament,” Current Psychology 14, 261–292 (1996).

[15] L. F. Barrett and J. A. Russell, “The Structure of Current Affect: Controversies and Emerging Consensus,” Current Directions in Psychological Science 8, 10–14 (1999).

[16] J. A. Russell, “Core affect and the psychological construction of emotion,” Psychological Review 110, 145– 172 (2003).

[17] J. Posner, J. A. Russell, and B. S. Peterson, “The circumplex model of affect: An integrative approach to affective neuroscience, cognitive development, and psychopathology,” Development and Psychopathology 17, 715–734 (2005).

[18] G. Mohammadi, D. Van De Ville, and P. Vuilleumier, “Brain networks subserving functional core processes of emotions identified with componential modeling,” Cerebral Cortex 33, 7993–8010 (2023).

[19] J. Leitão, B. Meuleman, D. V. D. Ville, and P. Vuilleumier, “Computational imaging during video game playing shows dynamic synchronization of cortical and subcortical networks of emotions,” PLOS Biology 18, e3000900 (2020).

[20] K. A. Lindquist, T. D. Wager, H. Kober, E. Bliss-Moreau, and L. F. Barrett, “The brain basis of emotion: A meta-analytic review,” Behavioral and Brain Sciences 35, 121–143 (2012).

[21] L. F. Barrett, “Solving the Emotion Paradox: Categorization and the Experience of Emotion,” Personality and Social Psychology Review 10, 20–46 (2006).

[22] L. F. Barrett, “Emotions are real.” Emotion 12, 413– 429 (2012).

[23] L. F. Barrett, “The Conceptual Act Theory: A Précis,” Emotion Review 6, 292–297 (2014).

[24] M. V. Peelen, A. P. Atkinson, and P. Vuilleumier, “Supramodal Representations of Perceived Emotions in the Human Brain,” Journal of Neuroscience 30, 10127– 10134 (2010).

[25] S. Hamann, “Mapping discrete and dimensional emotions onto the brain: Controversies and consensus,” Trends in Cognitive Sciences 16, 458–466 (2012).

[26] L. Nummenmaa, H. Saarimäki, E. Glerean, A. Gotsopoulos, I.P. Jääskeläinen, R. Hari, and M. Sams, “Emotional speech synchronizes brains across listeners and engages large-scale dynamic brain networks,” NeuroImage 102, 498–509 (2014).

[27] Y. Jacob, G. Gilam, T. Lin, G. Raz, and T. Hendler, “Anger Modulates Influence Hierarchies Within and Between Emotional Reactivity and Regulation Networks,” Frontiers in Behavioral Neuroscience 12 (2018), 10.3389/fnbeh.2018.00060.

[28] D. Ventura, L. Heredia, M. Torrente, and P. Vicens, “Automated Emotional Facial Expression Assessment and Emotional Elicitation through Film Clip Stimuli,” Studia Psychologica 62, 350–363 (2020).

[29] N. K. Logothetis, J. Pauls, M. Augath, T. Trinath, and A. Oeltermann, “Neurophysiological investigation of the basis of the fMRI signal,” Nature 412, 150–157 (2001).

[30] N. K. Logothetis, “The underpinnings of the BOLD functional magnetic resonance imaging signal,” Journal of Neuroscience 23, 3963–3971 (2003).

[31] N. K. Logothetis and B. A. Wandell, “Interpreting the BOLD signal,” Annu. Rev. Physiol. 66, 735–769 (2004).

[32] P. J. Drew, “Vascular and neural basis of the BOLD signal,” Current Opinion in Neurobiology Computational Neuroscience, 58, 61–69 (2019).

[33] K. L. Phan, T. Wager, S. F. Taylor, and I. Liberzon, “Functional Neuroanatomy of Emotion: A Meta-Analysis of Emotion Activation Studies in PET and fMRI,” NeuroImage 16, 331–348 (2002).

[34] K. L. Phan, S. F. Taylor, R. C. Welsh, L. R. Decker, D. C. Noll, T. E. Nichols, J. C. Britton, and I. Liberzon, “Activation of the medial prefrontal cortex and extended amygdala by individual ratings of emotional arousal: A fMRI study,” Biological Psychiatry 53, 211– 215 (2003).

[35] H. Kober, L. F. Barrett, J. Joseph, E. Bliss-Moreau, K. Lindquist, and T. D. Wager, “Functional grouping and cortical–subcortical interactions in emotion: A meta-analysis of neuroimaging studies,” NeuroImage 42, 998–1031 (2008).

[36] P. A. Kragel and K. S. LaBar, “Decoding the Nature of Emotion in the Brain,” Trends in Cognitive Sciences 20, 444–455 (2016).

[37] A. J. Shackman and T. D. Wager, “The emotional brain: Fundamental questions and strategies for future research,” Neuroscience letters 693, 68–74 (2019).

[38] E. Meaux and P. Vuilleumier, “Facing mixed emotions: Analytic and holistic perception of facial emotion expressions engages separate brain networks,” NeuroImage 141, 154–173 (2016).

[39] T. D. Wager, J. Kang, T. D. Johnson, T. E. Nichols, A. B. Satpute, and L. F. Barrett, “A Bayesian Model of Category-Specific Emotional Brain Responses,” PLOS Computational Biology 11, e1004066 (2015).

[40] H. Saarimäki, L. F. Ejtehadian, E. Glerean, I.P. Jääskeläinen, P. Vuilleumier, M. Sams, and L. Nummenmaa, “Distributed affective space represents multiple emotion categories across the human brain,” Social Cognitive and Affective Neuroscience 13, 471–482 (2018).

[41] N. Koide-Majima, T. Nakai, and S. Nishimoto, “Distinct dimensions of emotion in the human brain and their representation on the cortical surface,” NeuroImage 222, 117258 (2020).

[42] L. Pessoa, “Understanding emotion with brain networks,” Current Opinion in Behavioral Sciences Emotion-Cognition Interactions, 19, 19–25 (2018).

[43] H. Saarimäki, “Naturalistic Stimuli in Affective Neuroimaging: A Review,” Frontiers in Human Neuroscience 15 (2021), 10.3389/fnhum.2021.675068.

[44] I.P. Jääskeläinen, M. Sams, E. Glerean, and J. Ahveninen, “Movies and narratives as naturalistic stimuli in neuroimaging,” NeuroImage 224, 117445 (2021).

[45] E. S. Finn, “Is it time to put rest to rest?” Trends in Cognitive Sciences 25, 1021–1032 (2021).

[46] E. Morgenroth, L. Vilaclara, M. Muszynski, J. Gaviria, P. Vuilleumier, and D. Van De Ville, “Probing neurodynamics of experienced emotions—a Hitchhiker’s guide to film fMRI,” Social Cognitive and Affective Neuroscience 18, nsad063 (2023).

[47] U. Hasson, E. Yang, I. Vallines, D. J. Heeger, and N. Rubin, “A Hierarchy of Temporal Receptive Windows in Human Cortex,” Journal of Neuroscience 28, 2539–2550 (2008).

[48] U. Hasson, R. Malach, and D. J. Heeger, “Reliability of cortical activity during natural stimulation,” Trends in Cognitive Sciences 14, 40–48 (2010).

[49] S. Sonkusare, M. Breakspear, and C. Guo, “Naturalistic Stimuli in Neuroscience: Critically Acclaimed,” Trends in Cognitive Sciences 23, 699–714 (2019).

[50] G. Lettieri, G. Handjaras, E. Ricciardi, A. Leo, P. Papale, M. Betta, P. Pietrini, and L. Cecchetti, “Emotionotopy in the human right temporo-parietal cortex,” Nature Communications 10, 5568 (2019).

[51] E. S. Finn and P. A. Bandettini, “Movie-watching outperforms rest for functional connectivity-based prediction of behavior,” NeuroImage 235, 117963 (2021).

[52] R. M. Hutchison, T. Womelsdorf, E. A. Allen, P. A. Bandettini, V. D. Calhoun, M. Corbetta, S. Della Penna, J. H. Duyn, G. H. Glover, J. Gonzalez-Castillo, D. A. Handwerker, S. Keilholz, V. Kiviniemi, D. A. Leopold, F. de Pasquale, O. Sporns, M. Walter, and C. Chang, “Dynamic functional connectivity: Promise, issues, and interpretations,” NeuroImage Mapping the Connectome, 80, 360–378 (2013).

[53] M. G. Preti, T. A. Bolton, and D. Van De Ville, “The dynamic functional connectome: State-of-the-art and perspectives,” NeuroImage Functional Architecture of the Brain, 160, 41–54 (2017).

[54] M. J. Tobia, K. Hayashi, G. Ballard, I. H. Gotlib, and C. E. Waugh, “Dynamic functional connectivity and individual differences in emotions during social stress,” Human Brain Mapping 38, 6185–6205 (2017).

[55] L. Nummenmaa, E. Glerean, M. Viinikainen, I.P. Jääskeläinen, R. Hari, and M. Sams, “Emotions promote social interaction by synchronizing brain activity across individuals,” Proceedings of the National Academy of Sciences of the United States of America 109, 9599–9604 (2012).

[56] G. Raz, A. Touroutoglou, C. Wilson-Mendenhall, G. Gilam, T. Lin, T. Gonen, Y. Jacob, S. Atzil, R. Admon, M. Bleich-Cohen, A. Maron-Katz, T. Hendler, and L. F. Barrett, “Functional connectivity dynamics during film viewing reveal common networks for different emotional experiences,” Cognitive, Affective, & Behavioral Neuroscience 16, 709–723 (2016).

[57] J. Gaviria, G. Rey, T. Bolton, D. V. D. Ville, and P. Vuilleumier, “Dynamic functional brain networks underlying the temporal inertia of negative emotions,” NeuroImage 240, 118377 (2021).

[58] Y. Sun, J. Ma, M. Huang, Y. Yi, Y. Wang, Y. Gu, Y. Lin, L. M. W. Li, and Z. Dai, “Functional connectivity dynamics as a function of the fluctuation of tension during film watching,” Brain Imaging and Behavior 16, 1260–1274 (2022).

[59] J. Ke, H. Song, Z. Bai, M. D. Rosenberg, and Y. C. Leong, “Dynamic brain connectivity predicts emotional arousal during naturalistic movie-watching,” PLOS Computational Biology 21, e1012994 (2025).

[60] F. Z. Esfahlani, Y. Jo, J. Faskowitz, L. Byrge, D. P. Kennedy, O. Sporns, and R. F. Betzel, “High-amplitude cofluctuations in cortical activity drive functional connectivity,” Proceedings of the National Academy of Sciences 117, 28393–28401 (2020).

[61] J. Faskowitz, F. Z. Esfahlani, Y. Jo, O. Sporns, and R. F. Betzel, “Edge-centric functional network representations of human cerebral cortex reveal overlapping system-level architecture,” Nature Neuroscience 23, 1644–1654 (2020).

[62] J. C. Tanner, J. Faskowitz, L. Byrge, D. P. Kennedy, O. Sporns, and R. F. Betzel, “Synchronous high-amplitude co-fluctuations of functional brain networks during movie-watching,” Imaging Neuroscience 1, imag–1–00026 (2023).

[63] J. Tanner, J. Faskowitz, D. P. Kennedy, and R. F. Betzel, “Dynamic adaptation to novelty in the brain is related to arousal and intelligence,” Cerebral Cortex 36, bhag025 (2026).

[64] F. Battiston, G. Cencetti, I. Iacopini, V. Latora, M. Lucas, A. Patania, J. G. Young, and G. Petri, “Networks beyond pairwise interactions: Structure and dynamics,” Physics Reports 874, 1–92 (2020).

[65] L. Torres, A. S. Blevins, D. Bassett, and T. Eliassi-Rad, “The Why, How, and When of Representations for Complex Systems,” SIAM Review 63, 435–485 (2021).

[66] F. Battiston, E. Amico, A. Barrat, G. Bianconi, G. Ferraz de Arruda, B. Franceschiello, I. Iacopini, S. Kéfi, V. Latora, Y. Moreno, M. M. Murray, T. P. Peixoto, F. Vaccarino, and G. Petri, “The physics of higherorder interactions in complex systems,” Nature Physics 17, 1093–1098 (2021).

[67] C. Giusti, R. Ghrist, and D. S. Bassett, “Two’s company, three (or more) is a simplex: Algebraic-topological tools for understanding higher-order structure in neural data,” Journal of computational neuroscience 41, 1–14 (2016).

[68] F. E. Rosas, P. A. Mediano, M. Gastpar, and H. J. Jensen, “Quantifying high-order interdependencies via multivariate extensions of the mutual information,” Physical Review E 100, 32305 (2019).

[69] F. E. Rosas, P. A. M. Mediano, A. I. Luppi, T. F. Varley, J. T. Lizier, S. Stramaglia, H. J. Jensen, and D. Marinazzo, “Disentangling high-order mechanisms and high-order behaviours in complex systems,” Nature Physics 18, 476–477 (2022).

[70] A. I. Luppi, P. A. Mediano, F. E. Rosas, N. Holland, T. D. Fryer, J. T. O’Brien, J. B. Rowe, D. K. Menon, D. Bor, and E. A. Stamatakis, “A synergistic core for human brain evolution and cognition,” Nature Neuroscience 25, 771–782 (2022).

[71] T. F. Varley, M. Pope, J. Faskowitz, and O. Sporns, “Multivariate information theory uncovers synergistic subsystems of the human cerebral cortex,” Communications Biology 6, 1–12 (2023).

[72] T. F. Varley, M. Pope, M. Grazia, Joshua, and O. Sporns, “Partial entropy decomposition reveals higher-order information structures in human brain activity,” Proceedings of the National Academy of Sciences 120, e2300888120 (2023).

[73] G. Petri, P. Expert, F. E. Turkheimer, R. L. Carhart-Harris, D. Nutt, P. Hellyer, and F. Vaccarino, “Homological scaffolds of brain functional networks,” Journal of the Royal Society Interface 10, 186–198 (2014).

[74] A. E. Sizemore, J. E. Phillips-Cremins, R. Ghrist, and D. S. Bassett, “The importance of the whole: Topological data analysis for the network neuroscientist,” Network Neuroscience (Cambridge, Mass.) 3, 656–673 (2019).

[75] A. Santoro, F. Battiston, M. Lucas, G. Petri, and E. Amico, “Higher-order connectomics of human brain function reveals local topological signatures of task decoding, individual identification, and behavior,” Nature Communications 15, 10244 (2024).

[76] A. I. Luppi, P. A. M. Mediano, F. E. Rosas, J. Allanson, J. D. Pickard, R. L. Carhart-Harris, G. B. Williams, M. M. Craig, P. Finoia, A. M. Owen, L. Naci, D. K. Menon, D. Bor, and E. A. Stamatakis, “A synergistic workspace for human consciousness revealed by integrated information decomposition,” (2023), 10.7554/eLife.88173.1.

[77] A. I. Luppi, D. Golkowski, A. Ranft, R. Ilg, D. Jordan, D. Bzdok, A. M. Owen, L. Naci, E. A. Stamatakis, E. Amico, and B. Misic, “General anaesthesia decreases the uniqueness of brain functional connectivity across individuals and species,” Nature Human Behaviour 9, 987–1004 (2025).

[78] E. Combrisson, R. Basanisi, M. Neri, G. Auzias, G. Petri, D. Marinazzo, S. Panzeri, and A. Brovelli, “Higher-order and distributed synergistic functional interactions encode information gain in goal-directed learning,” bioRxiv, 2024–09 (2024).

[79] S. Poetto, H. Merritt, A. Santoro, G. Rabuffo, D. A. Battaglia, F. Vaccarino, M. Saggar, A. Brovelli, and G. Petri, “The topological architecture of brain identity,” bioRxiv (2025), 10.1101/2025.06.20.660792.

[80] T. F. Varley, O. Sporns, N. J. Stevenson, P. Yrjölä, M. G. Welch, M. M. Myers, S. Vanhatalo, and A. Tokariev, “Emergence of a synergistic scaffold in the brains of human infants,” Communications Biology 8, 1–13 (2025).

[81] M. Gatica, R. Cofré, P. A. Mediano, F. E. Rosas, P. Orio, I. Diez, S. P. Swinnen, and J. M. Cortes, “High-Order Interdependencies in the Aging Brain,” Brain Connectivity 00, 1–11 (2021).

[82] M. Gatica, F. E. Rosas, P. A. Mediano, I. Diez, S. P. Swinnen, P. Orio, R. Cofré, and J. M. Corts, “High-order functional redundancy in ageing explained via alterations in the connectome in a whole-brain model,” PLoS Computational Biology 18 (2022), 10.1371/journal.pcbi.1010431.

[83] M. Gatica, C. Atkinson-Clement, P. A. M. Mediano, M. Alkhawashki, J. Ross, J. Sallet, and M. Kaiser, “Transcranial ultrasound stimulation effect in the redundant and synergistic networks consistent across macaques,” Network Neuroscience, 1–30 (2024).

[84] M. Gatica, C. Atkinson-Clement, C. Coronel-Oliveros, M. Alkhawashki, P. A. M. Mediano, E. Tagliazucchi, F. E. Rosas, M. Kaiser, and G. Petri, “Understanding the high-order network plasticity mechanisms of ultrasound neuromodulation,” bioRxiv (2025), 10.1101/2025.01.11.632528.

[85] R. Herzog, F. E. Rosas, R. Whelan, S. Fittipaldi, H. Santamaria-Garcia, J. Cruzat, A. Birba, S. Moguilner, E. Tagliazucchi, P. Prado, et al., “Genuine highorder interactions in brain networks and neurodegeneration,” Neurobiology of Disease 175, 105918 (2022).

[86] L. Arbabyazd, S. Petkoski, M. Breakspear, A. Solodkin, D. Battaglia, and V. Jirsa, “State-switching and high-order spatiotemporal organization of dynamic functional connectivity are disrupted by alzheimer’s disease,” Network Neuroscience 7, 1420–1451 (2023).

[87] Q. Li, V. D. Calhoun, A. R. Ballem, S. Yu, J. Malo, and A. Iraji, “Aberrant high-order dependencies in schizophrenia resting-state functional mri networks,” arXiv preprint 2310.17445 (2023).

[88] I. Pirovano, Y. Antonacci, A. Mastropietro, C. Bara, L. Sparacino, E. Guanziroli, F. Molteni, M. Tettamanti, L. Faes, and G. Rizzo, “Rehabilitation modulates highorder interactions among large-scale brain networks in subacute stroke,” IEEE Transactions on Neural Systems and Rehabilitation Engineering 31, 4549–4560 (2023).

[89] H. Wang, Y. Liu, and Y. Ding, “Identifying diagnostic biomarkers for autism spectrum disorder from higherorder interactions using the ped algorithm,” Neuroin-formatics 22, 285–296 (2024).

[90] A. Santoro, M. Neri, S. Poetto, D. Orsenigo, M. Diano, M. Gatica, and G. Petri, “Charting higher-order models of brain function beyond pairwise interactions,” Nat. Commun. (2026), 10.1038/s41467-026-75959-w.

[91] E. Morgenroth, S. Moia, L. Vilaclara, R. Fournier, M. Muszynski, M. Ploumitsakou, M. Almató-Bellavista, P. Vuilleumier, and D. Van De Ville, “Emo-film: A multimodal dataset for affective neuroscience using naturalistic stimuli,” Scientific Data 12 (2025), 10.1038/s41597-025-04803-5.

[92] E. Morgenroth, R. Somarathna, D. Van De Ville, G. Mohammadi, and P. Vuilleumier, “Dissecting appraisal and multicomponential features of emotion: Evidence from multilevel annotation during naturalistic stimulation,” Emotion (2026), 10.1037/emo0001619.

[93] J. Chen, Y. C. Leong, C. J. Honey, C. H. Yong, K. A. Norman, and U. Hasson, “Shared memories reveal shared structure in neural activity across individuals,” Nature Neuroscience 20, 115–125 (2017).

[94] A. Zadbood, J. Chen, Y. Leong, K. Norman, and U. Hasson, “How We Transmit Memories to Other Brains: Constructing Shared Neural Representations Via Communication,” Cerebral Cortex 27, 4988–5000 (2017).

[95] J.-P. Eckmann, S. O. Kamphorst, and D. Ruelle, “Recurrence Plots of Dynamical Systems,” Europhysics Letters 4, 973 (1987).

[96] A. Schaefer, R. Kong, E. M. Gordon, T. O. Laumann, X.-N. Zuo, A. J. Holmes, S. B. Eickhoff, and B. T. T. Yeo, “Local-Global Parcellation of the Human Cerebral Cortex from Intrinsic Functional Connectivity MRI,” Cerebral Cortex 28, 3095–3114 (2018).

[97] M. F. Glasser, S. N. Sotiropoulos, J. A. Wilson, T. S. Coalson, B. Fischl, J. L. Andersson, J. Xu, S. Jbabdi, M. Webster, J. R. Polimeni, D. C. Van Essen, and M. Jenkinson, “The minimal preprocessing pipelines for the Human Connectome Project,” NeuroImage Mapping the Connectome, 80, 105–124 (2013).

[98] A. J. Gates, I. B. Wood, W. P. Hetrick, and Y.-Y. Ahn, “Element-centric clustering comparison unifies overlaps and hierarchy,” Scientific Reports 9, 8574 (2019).

[99] A. Krishnan, L. J. Williams, A. R. McIntosh, and H. Abdi, “Partial Least Squares (PLS) methods for neuroimaging: A tutorial and review,” NeuroImage Multivariate Decoding and Brain Reading, 56, 455–475 (2011).

[100] M. Helmer, S. Warrington, A.-R. Mohammadi-Nejad, J. L. Ji, A. Howell, B. Rosand, A. Anticevic, S. N. Sotiropoulos, and J. D. Murray, “On the stability of canonical correlation analysis and partial least squares with application to brain-behavior associations,” Communications Biology 7, 217 (2024).

[101] T. He, R. Kong, A. J. Holmes, M. Nguyen, M. R. Sabuncu, S. B. Eickhoff, D. Bzdok, J. Feng, and B. T. T. Yeo, “Deep neural networks and kernel regression achieve comparable accuracies for functional connectivity prediction of behavior and demographics,” NeuroImage 206, 116276 (2020).

[102] Y. Ma and P. A. Kragel, “Map-like representations of emotion knowledge in hippocampal-prefrontal systems,” Nature Communications 17, 1518 (2026).

[103] C. Nadeau and Y. Bengio, “Inference for the Generalization Error,” Machine Learning 52, 239–281 (2003).

[104] E. S. Finn, X. Shen, D. Scheinost, M. D. Rosenberg, J. Huang, M. M. Chun, X. Papademetris, and R. T. Constable, “Functional connectome fingerprinting: Identifying individuals using patterns of brain connectivity,” Nature Neuroscience 18, 1664–1671 (2015).

[105] H. Song, E. S. Finn, and M. D. Rosenberg, “Neural signatures of attentional engagement during narratives and its consequences for event memory,” Proceedings of the National Academy of Sciences 118, e2021905118 (2021).

[106] S. Haufe, F. Meinecke, K. Görgen, S. Dähne, J.-D. Haynes, B. Blankertz, and F. Bießmann, “On the interpretation of weight vectors of linear models in multivariate neuroimaging,” NeuroImage 87, 96–110 (2014).

[107] E. Sampaolo, G. Handjaras, G. Lettieri, and L. Cecchetti, “Lights, Camera, Emotion: REELMO’s 1060 Hours of Affective Reports to Explore Emotions in Naturalistic Contexts,” Scientific Data 12, 797 (2025).

[108] N. Kriegeskorte, M. Mur, and P. A. Bandettini, “Representational similarity analysis - connecting the branches of systems neuroscience,” Front. Syst. Neurosci. 2 (2008), 10.3389/neuro.06.004.2008.

[109] F. Battiston and G. Petri, Higher-Order Systems (Springer, 2022).

[110] G. Bianconi, Higher-Order Networks. An Introduction to Simplicial Complexes (Cambridge Elements, 2021).

[111] A. S. Cowen and D. Keltner, “Self-report captures 27 distinct categories of emotion bridged by continuous gradients,” Proceedings of the National Academy of Sciences of the United States of America 114, E7900– E7909 (2017).

[112] A. E. Skerry and R. Saxe, “Neural Representations of Emotion Are Organized around Abstract Event Features,” Current Biology 25, 1945–1954 (2015).

[113] T. Horikawa, A. S. Cowen, D. Keltner, and Y. Kamitani, “The Neural Representation of Visually Evoked Emotion Is High-Dimensional, Categorical, and Distributed across Transmodal Brain Regions,” iScience 23, 101060 (2020).

[114] P. A. Kragel, L. Koban, L. F. Barrett, and T. D. Wager, “Representation, Pattern Information, and Brain Signatures: From Neurons to Neuroimaging,” Neuron 99, 257–273 (2018).

[115] J. N. van der Meer, M. Breakspear, L. J. Chang, S. Sonkusare, and L. Cocchi, “Movie viewing elicits rich and reliable brain state dynamics,” Nature Communications 11, 5004 (2020).

[116] C. Rosen and D. J. Freedman, “How distributed is the brain-wide network that is recruited for cognition? Nature Reviews Neuroscience,” Nat. Rev. Neurosci. 27, 138–150 (2026).

[117] A. B. Satpute, P. A. Kragel, L. F. Barrett, T. D. Wager, and M. Bianciardi, “Deconstructing arousal into wakeful, autonomic and affective varieties,” Neuroscience Letters Functional Neuroimaging of the Emotional Brain, 693, 19–28 (2019).

[118] X. Kong, S. Li, and G. Gong, “Arousal modulates functional connectivity through structured and hemispherically asymmetric community architecture during wakefulness,” eLife 15, RP110294 (2026).

[119] J. M. Shine, P. G. Bissett, P. T. Bell, O. Koyejo, J. H. Balsters, K. J. Gorgolewski, C. A. Moodie, and R. A. Poldrack, “The dynamics of functional brain networks: integrated network states during cognitive task performance,” Neuron 92, 544–554 (2016).

[120] H. Kim, L. H. Somerville, T. Johnstone, S. Polis, A. L. Alexander, L. M. Shin, and P. J. Whalen, “Contextual Modulation of Amygdala Responsivity to Surprised Faces,” Journal of Cognitive Neuroscience 16, 1730– 1745 (2004).

[121] H. Okon-Singer, L. Lichtenstein-Vidne, and N. Cohen, “Dynamic modulation of emotional processing,” Biological Psychology Specificity, Methodology and Psychopathology of Emotional Attention, 92, 480–491 (2013).

[122] S. Iyer, E. Collier, T. W. Broom, E. S. Finn, and M. L. Meyer, “Individuals who see the good in the bad engage distinctive default network coordination during postencoding rest,” Proceedings of the National Academy of Sciences 121, e2306295121 (2024).

[123] E. Qiao-Tasserit, C. Corradi-Dell’Acqua, and P. Vuilleumier, “The good, the bad, and the suffering. Transient emotional episodes modulate the neural circuits of pain and empathy,” Neuropsychologia Special Issue: The Neuroscience of Empathy, 116, 99–116 (2018).

[124] P. Vrtička, D. Sander, and P. Vuilleumier, “Effects of emotion regulation strategy on brain responses to the valence and social content of visual scenes,” Neuropsychologia 49, 1067–1082 (2011).

[125] G. A. van Kleef and D. A. Sauter, “What Makes Emotions Positive or Negative? Repurposing Valence at the Level of Meta-Emotions,” Affect. Sci. (2026), 10.1007/s42761-026-00385-6.

[126] J. T. Cacioppo and G. G. Berntson, “Relationship between attitudes and evaluative space: A critical review, with emphasis on the separability of positive and negative substrates,” Psychological Bulletin 115, 401–423 (1994).

[127] B. Knutson, K. Katovich, and G. Suri, “Inferring affect from fMRI data,” Trends Cogn. Sci. 18, 422–428 (2014).

[128] K. C. Berridge and P. Dayan, “Liking,” Curr. Biol. 31, R1555–R1557 (2021).

[129] E. S. Finn, E. Glerean, A. Y. Khojandi, D. Nielson, P. J. Molfese, D. A. Handwerker, and P. A. Bandettini, “Idiosynchrony: From shared responses to individual differences during naturalistic neuroimaging,” NeuroImage 215, 116828 (2020).

[130] M. Nguyen, T. Vanderwal, and U. Hasson, “Shared understanding of narratives is correlated with shared neural responses,” NeuroImage 184, 161–170 (2019).

[131] L. F. Barrett, “Variety is the spice of life: A psychological construction approach to understanding variability in emotion,” Cogn. Emot. 23, 1284–1306 (2009).

[132] K. A. Lindquist, “Emotions Emerge from More Basic Psychological Ingredients: A Modern Psychological Constructionist Model,” Emot. Rev. 5, 356–368 (2013).

[133] S. B. Eickhoff, M. Milham, and T. Vanderwal, “Towards clinical applications of movie fMRI,” NeuroImage 217, 116860 (2020).

[134] M. D. Rosenberg and E. S. Finn, “How to establish robust brain–behavior relationships without thousands of individuals,” Nature Neuroscience 25, 835–837 (2022).

[135] L. Pessoa, “The entangled brain,” Journal of Cognitive Neuroscience 35, 349–360 (2023).

[136] F. Antonelli, F. Bernardi, A. Koul, G. Novembre, and F. Papaleo, “Emotions in multi-brain dynamics: A promising research frontier,” Neuroscience & Biobehavioral Reviews 168, 105965 (2025).

[137] H. Merritt, A. Mejia, and R. Betzel, “The dual interpretation of edge time series: Time-varying connectivity versus statistical interaction,” iScience 29 (2026), 10.1016/j.isci.2026.115949.

[138] H. Edelsbrunner, D. Letscher, and A. Zomorodian, “Topological persistence and simplification,” in Proceedings 41st Annual Symposium on Foundations of Computer Science (2000) pp. 454–463.

[139] A. Zomorodian and G. Carlsson, “Computing Persistent Homology,” Discrete & Computational Geometry 33, 249–274 (2005).

[140] L. Caputi, A. Pidnebesna, and J. Hlinka, “Promises and pitfalls of topological data analysis for brain connectivity analysis,” NeuroImage 238, 118245 (2021).

[141] L.-D. Lord, P. Expert, H. M. Fernandes, G. Petri, T. J. Van Hartevelt, F. Vaccarino, G. Deco, F. Turkheimer, and M. L. Kringelbach, “Insights into Brain Architectures from the Homological Scaffolds of Functional Connectivity Networks,” Frontiers in Systems Neuroscience 10 (2016), 10.3389/fnsys.2016.00085.

[142] V. D. Blondel, J.-L. Guillaume, R. Lambiotte, and E. Lefebvre, “Fast unfolding of communities in large networks,” Journal of Statistical Mechanics: Theory and Experiment 2008, P10008 (2008).

[143] A. Lancichinetti and S. Fortunato, “Consensus clustering in complex networks,” Scientific Reports 2, 1–7 (2012).

[144] A. M. Winkler, G. R. Ridgway, M. A. Webster, S. M. Smith, and T. E. Nichols, “Permutation inference for the general linear model,” NeuroImage 92, 381–397 (2014).

[145] X. Shen, E. S. Finn, D. Scheinost, M. D. Rosenberg, M. M. Chun, X. Papademetris, and R. T. Constable, “Using connectome-based predictive modeling to predict individual behavior from brain connectivity,” Nature Protocols 12, 506–518 (2017).

[146] D. Prichard and J. Theiler, “Generating surrogate data for time series with several simultaneously measured variables,” Physical Review Letters 73, 951–954 (1994).

[147] B. T. Thomas Yeo, F. M. Krienen, J. Sepulcre, M. R. Sabuncu, D. Lashkari, M. Hollinshead, J. L. Roffman, J. W. Smoller, L. Zöllei, J. R. Polimeni, B. Fischl, H. Liu, and R. L. Buckner, “The organization of the human cerebral cortex estimated by intrinsic functional connectivity,” Journal of Neurophysiology 106, 1125– 1165 (2011).

[148] H. Popal, Y. Wang, and I. R. Olson, “A Guide to Representational Similarity Analysis for Social Neuroscience,” Social Cognitive and Affective Neuroscience 14, 1243– 1253 (2019).

[149] A. Bryant, “Subcortex visualization: A toolbox for custom data visualization in the subcortex and cerebellum,” Aperture Neuro 6 (2026), 10.52294/001c.165884.

[150] E. Morgenroth, S. Moia, L. Vilaclara, R. Fournier, M. Muszynski, M. Ploumitsakou, M. Almató-Bellavista, P. Vuilleumier, and D. Van De Ville, “Emo-FilM: A multimodal dataset for affective neuroscience using naturalistic stimuli,” (2024).

[151] G. Peeters et al., “A large set of audio features for sound description (similarity and classification) in the cuidado project,” CUIDADO Ist Project Report 54, 1– 25 (2004).

[152] B. McFee, C. Raffel, D. Liang, D. Ellis, M. McVicar, E. Battenberg, and O. Nieto, “Librosa: Audio and Music Signal Analysis in Python,” in Python in Science Conference (Austin, Texas, 2015) pp. 18–24.

[153] G. Tzanetakis and P. Cook, “Musical genre classification of audio signals,” IEEE Trans. Speech Audio Process. 10, 293–302 (2002).

[154] P. Valdez and A. Mehrabian, “Effects of color on emotions,” J. Exp. Psychol. Gen. 123, 394–409 (1994).

[155] A. Hanjalic and LQ. Xu, “Affective video content representation and modeling,” IEEE Trans. Multimed. 7, 143–154 (2005).

[156] Y. Baveye, E. Dellandréa, C. Chamaret, and L. Chen, “LIRIS-ACCEDE: A Video Database for Affective Content Analysis,” IEEE Trans. Affect. Comput. 6, 43–55 (2015).

[157] P. Mergny and L. Zdeborová, “Spectral Thresholds in Correlated Spiked Models and Fundamental Limits of Partial Least Squares,” (2025), 2510.17561 [math.ST].

[158] M. Danyluik, Y. Zeighami, A. Mukora, M. Lepage, J. Shah, R. Joober, B. Misic, Y. Iturria-Medina, and M. M. Chakravarty, “Evaluating permutation-based inference for partial least squares analysis of neuroimaging data,” Imaging Neurosci. 3, imag a 00434 (2025).

[159] R. Tibshirani, “Regression Shrinkage and Selection Via the Lasso,” Journal of the Royal Statistical Society: Series B (Methodological) 58, 267–288 (1996).

